# Lightning Pose: improved animal pose estimation via semi-supervised learning, Bayesian ensembling, and cloud-native open-source tools

**DOI:** 10.1101/2023.04.28.538703

**Authors:** Dan Biderman, Matthew R Whiteway, Cole Hurwitz, Nicholas Greenspan, Robert S Lee, Ankit Vishnubhotla, Richard Warren, Federico Pedraja, Dillon Noone, Michael Schartner, Julia M Huntenburg, Anup Khanal, Guido T Meijer, Jean-Paul Noel, Alejandro Pan-Vazquez, Karolina Z Socha, Anne E Urai, The International Brain Laboratory, John P Cunningham, Nathaniel B Sawtell, Liam Paninski

## Abstract

Contemporary pose estimation methods enable precise measurements of behavior via supervised deep learning with hand-labeled video frames. Although effective in many cases, the supervised approach requires extensive labeling and often produces outputs that are unreliable for downstream analyses. Here, we introduce “Lightning Pose,” an efficient pose estimation package with three algorithmic contributions. First, in addition to training on a few labeled video frames, we use many unlabeled videos and penalize the network whenever its predictions violate motion continuity, multiple-view geometry, and posture plausibility (semi-supervised learning). Second, we introduce a network architecture that resolves occlusions by predicting pose on any given frame using surrounding unlabeled frames. Third, we refine the pose predictions post-hoc by combining ensembling and Kalman smoothing. Together, these components render pose trajectories more accurate and scientifically usable. We release a cloud application that allows users to label data, train networks, and predict new videos directly from the browser.

## 1 Introduction

Behavior is our window into the processes that underlie animal intelligence, ranging from early sensory processing to complex social interaction [1]. Methods for automatically quantifying behavior from video [2–4] have opened the door to high-throughput experiments that compare animal behavior across pharmacological [5] and disease [6] conditions. Moreover, when behavior is carefully monitored, motor signals are revealed in unexpected brain areas, even regions classically defined to be purely sensory [7, 8].

Pose estimation methods based on fully-supervised deep learning have emerged as a workhorse for behavioral quantification [9–13]. This technology reduces high-dimensional videos of behaving animals to low-dimensional time series of their poses, defined in terms of a small number of user-selected keypoints per video frame. Three steps are required to accomplish this feat. Users first create a training dataset by manually labeling poses on a subset of video frames; typically hundreds or thousands of frames are labeled to obtain reliable pose estimates. A neural network is then trained to predict poses that match user labels. Finally, the network is run on a new video to predict a pose for each frame separately. This process of labeling-training-prediction can be iterated until performance is satisfactory. The resulting pose estimates are used extensively in downstream analyses including quantifying predefined behavioral features (e.g., gait features such as stride length, or social features such as distance between subjects), estimation of neural encoding and decoding models, classification of behaviors into discrete “syllables,” and closed-loop experiments [14–19].

Although the supervised paradigm is effective in many cases, a number of critical roadblocks remain. To start, the labeling process can be laborious, especially when labeling complicated skeletons on multiple views. Even with large labeled datasets, trained networks are often unreliable: they output “glitchy” predictions that require further manipulation before downstream analyses [20, 21], and struggle to generalize to animals and sessions that were not represented in their labeled training set. Even well-trained networks that achieve low pixel error on a small number of labeled test frames can still produce error frames that hinder downstream scientific tasks. Manually identifying these error frames is like finding a needle in a haystack [22]: errors persist for a few frames at a time whereas behavioral videos can be hours long. Automatic approaches – currently limited to filtering low-confidence predictions and temporal discontinuities – can easily miss scientifically critical errors.

To improve the robustness and usability of animal pose estimation, we present *Lightning Pose*, a solution at three levels: modeling, software, and a cloud-based application.

First, we leverage semi-supervised learning, which involves training networks on both labeled frames and unlabeled videos, and is known to improve generalization and data-efficiency [23]. On unlabeled videos, the networks are trained to minimize a number of unsupervised losses that encode our prior beliefs about moving bodies: poses should evolve smoothly in time, be physically plausible, and be localized consistently when seen from multiple views. In addition, we leverage unlabeled frames in a *Temporal Context Network* architecture, which instead of taking in a single frame at a time, processes each frame with its neighboring (unlabeled) frames. Our resulting models outperform their purely supervised counterparts across a range of metrics and datasets, providing more reliable predictions for downstream analyses.

We further improve our networks’ predictions using a general Bayesian post-processing approach, which we coin the *Ensemble Kalman Smoother*: we aggregate (“ensemble”) the predictions of multiple networks – which is known to improve their accuracy and robustness [24, 25] — and model those aggregated predictions with a spatially-constrained Kalman smoother that takes their collective uncertainty into account.

We implemented these tools in a deep learning software package that capitalizes on recent advances in the deep learning ecosystem. Open-source technologies allow us to outsource engineering-heavy tasks (such as GUI development, or training orchestration), which simplifies our package and allows users to focus on scientific modeling decisions. We name our package *Lightning Pose*, as it is based on the PyTorch Lightning deep learning library [26]. Unlike most existing packages, Lightning Pose is video-centric and built for manipulating large videos directly on the GPU, to support our semi-supervised training (and enable fast evaluation on new videos). Our modular design allows users to quickly prototype new training objectives and network architectures without affecting any aspects of training.

Finally, to make pose estimation tools accessible to the broader audience in life sciences, their adoption should not depend on programming skills or access to specialized hardware. Therefore, we developed a no-install cloud application that runs on the browser and allows users to perform the entire cycle of pose estimation: uploading raw videos to the cloud, annotating frames, training networks, and diagnosing the reliability of the results using our unsupervised loss terms.

## 2 Results

We first describe the dominant supervised approach to pose estimation and illustrate its drawbacks, especially when applied to new subjects and sessions. Next, we introduce our unsupervised losses and Temporal Context Network architecture. We illustrate that these unsupervised losses can be used to identify outlier predictions in unlabeled videos, and find that networks trained with these losses lead to more reliable tracking compared to purely supervised models. We then introduce the Ensemble Kalman Smoother post-processing approach and show that it further improves tracking performance. We proceed to apply our combined methods to the International Brain Lab datasets, and show that they improve pupil and paw tracking, thereby improving neural decoding. Finally, we showcase our software package and cloud-hosted application. Further details on our models, losses, and training protocol are provided in the Methods.

### 2.1 Supervised pose estimation and its limitations

The leading packages for animal pose estimation – DeepLabCut [9], SLEAP [10], DeepPoseKit [11], and others – differ in architectures and implementation but all perform supervised heatmap regression on a frame-by-frame basis (Fig. 1A). A standard model is composed of a “backbone” that extracts features for each frame (e.g., a ResNet-50 network) and a “head” that uses these features to predict body part location heatmaps. Networks are trained to match their outputs to manual labels.

**Figure 1.**
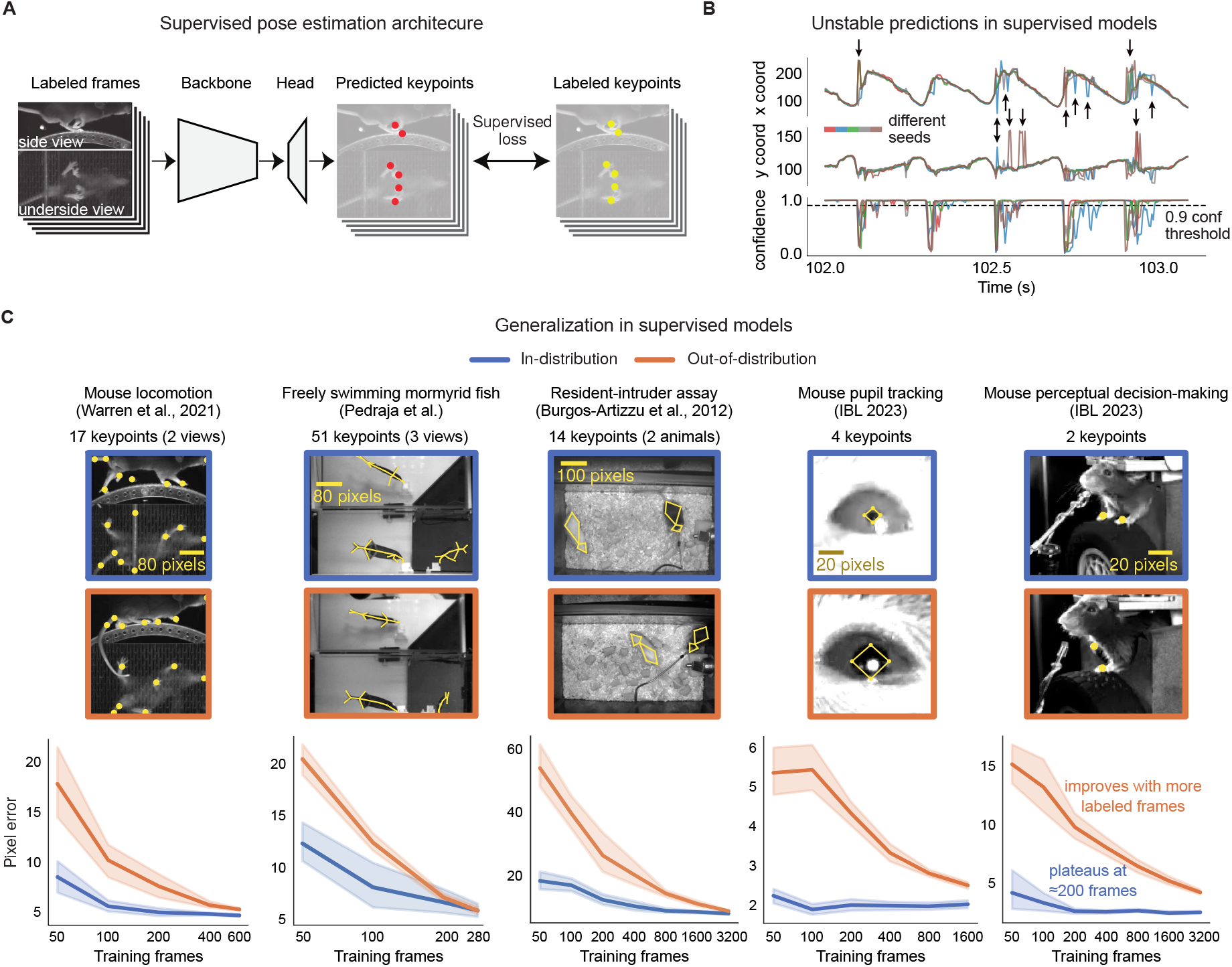
Fully-supervised pose estimation often outputs unstable predictions and requires many labels to generalize to new animals. **A**. Diagram of a typical pose estimation model trained with supervised learning, illustrated using the mirror-mouse dataset [16]. A dataset is created by labeling keypoints on a subset of video frames. A convolutional neural network, consisting of a “backbone” and a prediction “head,” takes in a batch of frames as inputs, and predicts a set of keypoints for each frame. It is trained to minimize the distance from the labeled keypoints. **B**. Predictions from five supervised DeepLabCut networks (trained with 631 labeled frames on the mirror-mouse dataset), for the left front paw position (top view) during one second of running behavior. *Top*: *x*-coordinate; *Middle*: *y*-coordinate; *Bottom*: confidence, applying a standard 0.9 threshold in a dashed line. The predictions demonstrate occasional discontinuities and disagreements across the five networks, only some of which are flagged by low confidence (Supplementary Video 1). **C**. To generalize robustly to unseen animals, many more labels are required. Top row shows five example datasets. Each blue image is an example taken from the in-distribution (InD) test set, which contains new images of animals that were seen in the training set. The orange images are test examples from unseen animals altogether, which we call the out-of-distribution (OOD) test set. Bottom row shows data-efficiency curves, measuring test-set pixel error as a function of the training set size. InD pixel error is in blue and OOD in orange. Line plots show the mean pixel error across all keypoints and frames *±* standard error over n=10 random subsets of InD training data.

Training supervised networks from a random initialization requires a large amount of labeled frames. Existing methods [9] circumvent this requirement by relying on *transfer learning*: *pre-training* a network on one task (e.g., image classification on ImageNet, with over one million labeled frames), and *fine-tuning* it on another task, in this case pose estimation, using far fewer labeled frames (*∼*100s-1000s). Typically, the backbone is fine-tuned and the head is trained from a random initialization.

After training, a fixed model is evaluated on a new video by predicting pose on each frame separately. Each predicted keypoint on each frame is accompanied by an estimate of the network’s confidence for that prediction; often low-confidence estimates are dropped in a post-processing step to reduce tracking errors.

Even when trained with many labeled frames, pose estimation network outputs may still be erroneous. We highlight this point using the “mirror-mouse” dataset, which features a head-fixed mouse running on a wheel and performing a sensory-guided locomotion task ([16]; see Methods). Using a camera and a bottom mirror, the mouse’s side and underside are observed simultaneously, recorded at 250 frames per second. 17 body parts are tracked, including all four paws in both views. We trained five DeepLabCut networks on 631 labeled frames (for each network, we used a different random seed to split the labeled frames into train and test sets).

Figure 1B shows the time series of the estimated left hind paw position during one second of a running behavior for each of the five networks (in colors). Each time series exhibits the expected periodic pattern (due to the running gait), but includes numerous “glitches,” some of which are undetected by the networks’ confidence. This collection of five networks – also known as a “deep ensemble” [24] – outputs highly variable predictions on many frames, especially in challenging moments of ambiguity or occlusion (Supplementary Video 1). We will later use this ensemble variance as a proxy for frame “difficulty.”

### 2.2 Supervised networks need more labeled data to generalize

It is standard to train a pose estimator using a representative sample of subjects, evaluate performance on held-out examples from that sample (“In Distribution” test set, henceforth InD), and then deploy the network for incoming data. The incoming data may include new subjects, seen from slightly different angles and lighting conditions (“Out of Distribution” test set, henceforth OOD). Differences between the InD and OOD test sets are termed “OOD shifts”; building models that are robust to such shifts is a contemporary frontier in machine learning research [27, 28].

We analyze five datasets: the “mirror-mouse” dataset introduced above [16], a freely swimming Mormyrid fish imaged with a single camera and two mirrors (for three views total; “mirror-fish,” Supplementary Fig. 1), a resident-intruder assay (“CRIM13;” two camera views are available but we consider the top view only; [29]), paw tracking in a head-fixed mouse (“IBL-paw;” three camera views are available but we only use the two side cameras; [30]), and a crop of the pupil area in IBL-paw (“IBL-pupil;” we use just one camera view). We split each labeled dataset into two cohorts of subjects, InD and OOD (see dataset and split details in Methods and Supplementary Table 1).

We train supervised heatmap regression networks that use a pretrained ResNet-50 backbone, similar to DeepLabCut (see Methods for architectural details) on InD data with an increasing number of labeled frames. Ten networks are trained per condition, each on a different random subset of InD data. We evaluate the networks’ performance on held-out InD and OOD labeled examples.

In Fig. 1C, we first replicate the observation that InD test-set error (blue curve) plateaus starting from *∼*200 labeled frames [18]. From looking at this curve in isolation, it could be inferred that additional manual annotation is unnecessary. However, the OOD error curve (orange) is both overall higher, and keeps steeply declining as more labels are added. To obtain an OOD error comparable to InD, many more labels will be needed. This larger label requirement is consistent with recent work showing that *∼*50k labeled frames are needed to robustly track ape poses [31], and that mouse face tracking networks need to be explicitly finetuned on labeled OOD data to achieve good performance [32]. For scarce labels, we find the gap between InD and OOD errors to be so large for some datasets that it renders prediction on new animals unusable for many downstream analyses.

To address these limitations, we propose the *Lightning Pose* framework, comprising two components: semisupervised learning and a Temporal Context Network architecture, which we describe next.

### 2.3 Semi-supervised learning via spatiotemporal constraints

Most animal pose estimation algorithms treat body parts as independent in time and space. Moreover, they do not utilize the vast amounts of available unlabeled videos for training the networks; instead, most video data are used just at prediction time. These two observations offer an opportunity for semi-supervised learning [23]. We thus train a network on both labeled frames (supervised) and large volumes of unlabeled videos (unsupervised). During training, the network is penalized whenever its pose predictions violate a set of spatiotemporal constraints on the unlabeled videos. We use “soft” constraints, i.e., the network is penalized only for severe constraint violations (with a controllable threshold parameter *E*). The unsupervised losses are applied only during training and not during video prediction. As a result, after training, a semisupervised model predicts a video as quickly as its fully-supervised counterpart.

Our semi-supervised pose estimation paradigm is depicted in Fig. 2A. The top row, shaded in gray, is simply the supervised pose estimation approach à la DeepLabCut. In each training iteration, the network additionally receives an unlabeled video clip (selected at random from a queue of videos), and outputs a time-series of pose predictions - one pose vector for each frame (bottom row). Those predictions are subjected to our unsupervised losses. We describe these unsupervised losses next.

**Figure 2.**
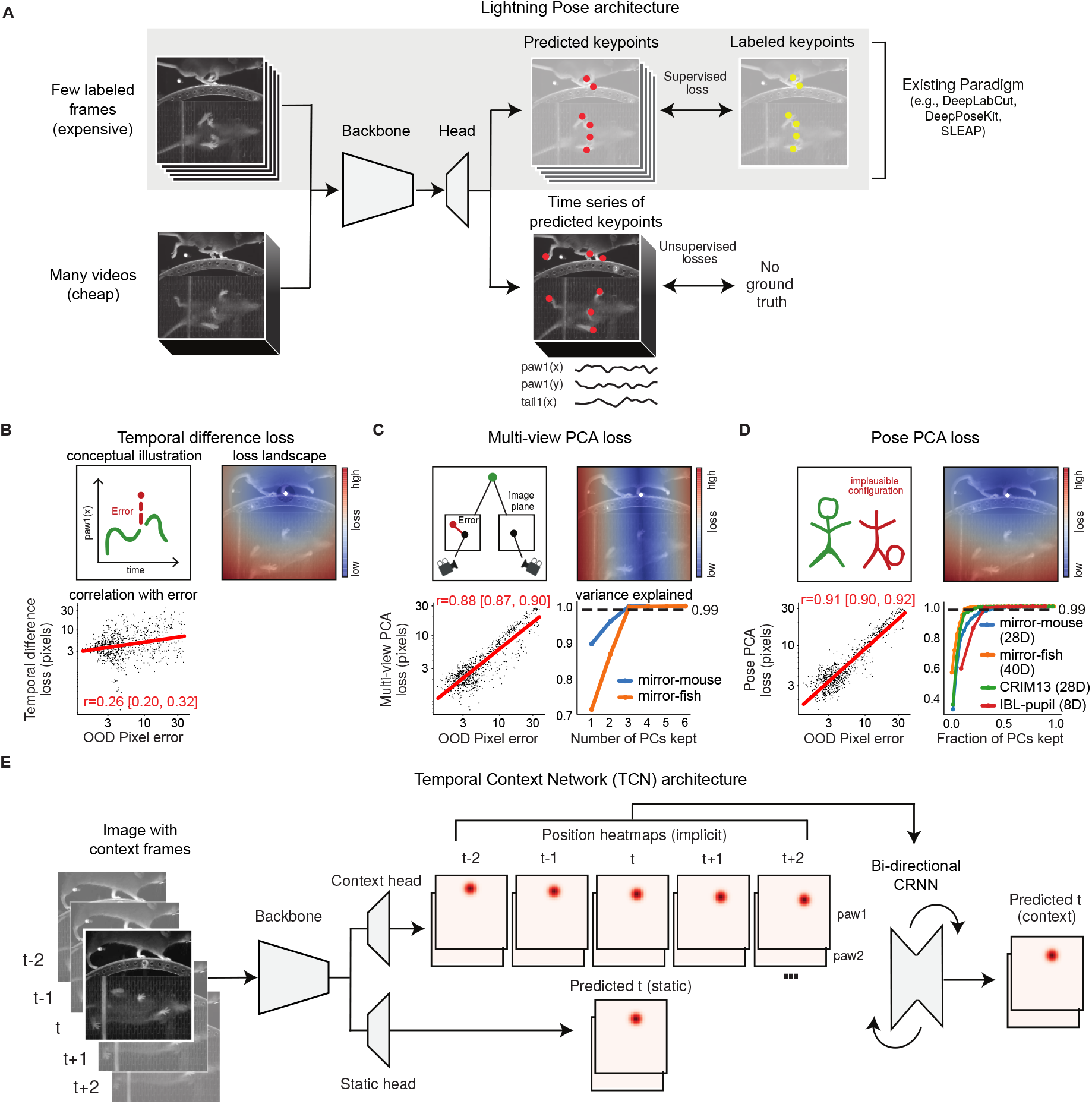
Lightning Pose exploits unlabeled data in pose estimation model training. **A**. Diagram of our semisupervised model that contains supervised (top row) and unsupervised (bottom row) components. **B**. Temporal difference loss penalizes jump discontinuities in predictions. Top left: illustration of a jump discontinuity. Top right: loss landscape for frame *t* given the prediction at *t −* 1 (white diamond), for the left front paw (top view). The loss increases further away from the previous prediction, and the dark blue circle corresponds to the maximum allowed jump, below which the loss is set to zero. Bottom left: correlation between temporal difference loss and pixel error on labeled test frames. **C**. Multi-view PCA loss constrains each multi-view prediction of the same body part to lie on a three-dimensional subspace found by Principal Component Analysis (PCA). Top left: illustration of a 3D keypoint detected on the imaging plane of two cameras. The left detection is inconsistent with the right. Top right: loss land-scape for the left front paw (top view; white diamond) given its predicted location on the bottom view. The blue band of low loss values is an “epipolar line” on which the top-view paw could be located. Bottom left: multi-view PCA loss is strongly correlated with pixel error. Bottom right: three PCs explain *>*99% of label variance on multi-view datasets. **D**. Pose PCA loss constrains predictions to lie on a low-dimensional subspace of plausible poses, found by PCA. Top left: illustration of a plausible and implausible poses. Top right: loss landscape for the left front paw (top view; white diamond) given all other keypoints, which is minimized around the paw’s actual position. Bottom left: Pose PCA loss is strongly correlated with pixel error. Bottom right: cumulative variance explained versus fraction of PCs kept. Across four datasets, *>* 99% of the variance in the pose vectors can be explained with *>*50% of the PCs. **E**. The Temporal Context Network processes each labeled frame with its adjacent unlabeled frames, using a bi-directional convolutional recurrent neural network. It forms two sets of location heatmap predictions, one using single-frame information and another using temporal context.

### 2.4 Temporal difference loss

The first spatiotemporal constraint we introduce is also one held by 4-month-old infants: objects should move continuously [33] and not jump too far between video frames. We define the temporal difference loss for each body part as the Euclidean distance between consecutive predictions in pixels. Similar losses have been used by several practitioners to detect outlier predictions post-hoc [16, 32], whereas our goal here, following [34], is to incorporate these penalties directly into network training to achieve more accurate network output. Figure 2B illustrates this penalty: the cartoon in the left panel indicates a jump discontinuity we would like to penalize. In the right panel we plot the loss landscape, evaluating the loss for every pixel in the image. The paw’s previous predicted position is depicted as a white diamond. Observe a ball of zero-loss values centered at the diamond with a radius of *ϵ* = 20 pixels which we set as the maximum allowed jump for this dataset; *ϵ* can be set depending on the frame rate, frame size, the camera’s distance from the subject, and how quickly or jerkily the subject moves. Outside the ball, the loss increases as we move farther away from the previous prediction.

If our losses are indeed viable proxies for pose prediction errors, they should be correlated with pixel errors in test frames for which we have ground-truth annotations. To test this, we trained a supervised model with 75 labeled frames, and computed the temporal difference loss on fully-labeled OOD frames. We anticipate a mild correlation with pixel error: prediction errors may persist across multiple frames and exhibit low temporal difference loss; in periods of fast motion, temporal difference loss may be high, yet keypoints may remain easily discernible. Indeed, in the bottom left panel of Fig. 2B, we see that the temporal difference loss is mildly correlated with pixel error on these frames (log-linear regression: Pearson *r* = 0.26, 95% CI = [0.20, 0.32]; each point is the mean across all keypoints for a given frame). As a comparison, confidence is a more reliable predictor of pixel error (Pearson *r* = *−*0.54, 95% CI = [*−*0.59, *−*0.49]).

### 2.5 Multi-view PCA loss

Our cameras see three-dimensional bodies from a two-dimensional perspective. It is increasingly common to record behavior using multiple synchronized cameras, train a network to estimate pose independently in each 2D view, and then use standard techniques post-hoc to fuse those 2D pose predictions into a 3D pose [20, 35]. This approach has two limitations. First, to reconstruct 3D poses, one needs to calibrate each camera, that is, to precisely infer where it is in the 3D world and carefully model its intrinsic parameters such as focal length and distortion. This typically involves filming a calibration board from all cameras after any camera adjustment; this adds experimental complexity and may be challenging or unreliable in some geometrically constrained experimental setups built for small model organisms. Second, localizing a body part in one view will constrain its allowed location in all other views [36], and we want to exploit this structure during training to obtain a stronger network; the post-hoc 3D reconstruction does not take advantage of this important structure to improve network training.

We use a “multi-view PCA” loss that constrains the predictions for unlabeled videos to be consistent across views [37, 38], while bypassing the need for complicated camera calibration. Each multi-view prediction (containing width-height coordinates for a single body part seen from multiple views) is compressed to three dimensions via simple principal components analysis (PCA; see Methods), and then this three-dimensional representation is linearly projected back into the original pixel coordinates (henceforth, “PCA reconstruction”). If the predictions are consistent across views and nonlinear camera distortion is negligible, no information should be lost when linearly compressing to three dimensions. We define the multi-view PCA loss as the pixel error between the original versus the PCA-reconstructed prediction, averaged across keypoints and views.

This simple linear approach will not be robust to substantial nonlinear distortions coming either from the lens or from a water medium. In both the mirror-mouse (two views) and mirror-fish (three views) datasets, distortions were minimized by placing the camera far from the subject (*∼* 1.1 and *∼* 1.7 meters respectively). Indeed, in both cases, three PCA dimensions explain *>* 99.9% of the multi-view ground truth label variance (Fig. 2C, bottom right).

Figure 2C (top left) provides a cartoon illustration of the idea we have just described: an inconsistent detection by the left camera will result in high multi-view loss. In the top right panel we compute the loss landscape for the left front paw on the top view, given its position in the bottom view. According to principles of multiple-view geometry, a point identified in one camera constrains the corresponding point in a second camera to a specific line, known as the “epipolar line” [36]. Indeed, the loss landscape exhibits a line of low loss values (blue) that intersects with the paw’s true location. Finally, as we did for the temporal difference loss, we compute the correlation between the multi-view loss and objective prediction errors for a test-set of labeled OOD frames. The multi-view loss is strongly correlated with pixel error (Pearson *r* = 0.88, 95% CI = [0.87, 0.90]), much more so than the temporal difference loss or confidence, motivating its use both as a post-hoc quality metric and as a penalty during training.

### 2.6 Pose PCA loss

Not all body configurations are feasible, and of those that are feasible, many are unlikely. Even diligent yoga practitioners will find their head next to their foot only on rare occasions (Fig. 2D, top left). In other words, in many pose estimation problems there are fewer degrees of freedom than there are body parts. The Pose PCA loss constrains the full predicted pose (over all keypoints) to lie on a low-dimensional subspace of feasible and likely body configurations. It is defined as the pixel error between an original pose prediction and its reconstruction after low-dimensional compression (see Methods).

Our loss is inspired by the success of low-dimensional models in capturing biological movement [39], ranging from worm locomotion [40] to human hand grasping [41]. We similarly find that across four of our datasets, 99% of the pose variance can be explained with far fewer dimensions than the number of pose coordinates (Fig. 2D, bottom right) – mirror-mouse: 14/28 components; mirror-fish: 8/40; CRIM13: 8/28; IBL-pupil 3/8 (IBL-paw only contains four dimensions). The effective pose dimensionality depends on the complexity of behavior, the keypoints selected for labeling, and the quality of the labeling. Sets of spatially-correlated keypoints will have a lower effective dimension (relative to the total number of keypoints); label errors tend to reduce these correlations and inflate the effective dimension.

Fig. 2D (top right) shows the Pose PCA loss landscape for the left hind paw location in the mirror-mouse dataset (true location shown as a white diamond) given the location of all the other body parts. As desired, the Pose PCA loss is lower around the paw’s true location and accommodates plausible neighbouring locations. Here too, the Pose PCA loss closely tracks ground truth pixel error on labeled OOD frames (Fig. 2D, bottom left; Pearson *r* = 0.91, 95% CI = [0.90, 0.92]).

The Pose PCA loss might erroneously penalize valid postures that are not represented in the labeled dataset. To test the prevalence of this issue, we took DeepLabCut models trained with abundant labels and computed the Pose PCA loss on held-out videos. We collected 100 frames with the largest Pose PCA loss per dataset. Manual labeling revealed that 85/100 (mirror-mouse; Supplementary Video 2), 87/100 (mirror-fish; Supplementary Video 3), and 100/100 (CRIM13; Supplementary Video 4) of the frames include true errors, indicating that in most cases, large Pose PCA losses correspond to pose estimation errors, rather than unseen rare poses.

### 2.7 Temporal Context Network

Some frames are more challenging to label than others, due to occlusions or ambiguities between similar body parts. In many cases, additional *temporal context* can help resolve ambiguities: e.g., if a keypoint is occluded briefly we can scroll backwards and forwards in the video to help “fill in the gaps.” However, this useful temporal context is not provided to standard frame-by-frame pose estimation architectures, which instead must make guesses about such challenging keypoint locations given information from just one frame at a time.

Therefore, we propose a Temporal Context network (TCN), illustrated in Fig. 2E, which uses a 2*J* + 1 frame sequence to predict the location heatmaps for the middle (i.e., *J* + 1) frame. As in the standard architecture, the TCN starts by pushing each image through a neural network backbone that computes useful features from each frame. Then, instead of predicting the pose directly from each of these individual per-frame feature vectors, we combine this information across frames using a bi-directional convolutional recurrent neural network (CRNN), and then use the output of the CRNN to form predictions for the middle frame.

The CRNN is lightweight compared to the backbone, and we only apply the backbone once per frame; therefore the TCN runtime scales linearly with the number of total context frames. We have found that a context window of 5 frames (i.e., *J* = 2) provides an effective balance between speed and accuracy and have used this value throughout the paper. In practice, the output of the TCN and single-frame architectures usually match on visible keypoints; rather, the TCN helps with more rare occlusions and ambiguities, where predictions from a single-frame architecture might jump to a different region of the image.

### 2.8 Spatiotemporal losses enhance outlier detection

Before training networks with these spatiotemporal losses, we first assess whether violations of these losses correspond to meaningful errors in video predictions, going beyond correlations with pixel errors on relatively small labeled test sets (Fig. 2B,C,D). Practitioners often detect outliers using a combination of low confidence and large temporal difference loss [16, 20, 32, 42]. Here we show that the standard approach can be complemented by multi-view and Pose PCA, which capture additional unique outliers.

We start with an example video snippet from the mirror-mouse dataset, focusing on the left hind paw on the bottom view (Fig. 3A,B). We analyze the predictions from a DeepLabCut model (trained as in Fig. 1B). Fig. 3A shows that the *x* coordinate’s discontinuity in frames 290-294, for example, is a result of the network switching back and forth between the similar-looking front and hind paws. These common “paw-switching” errors are mostly missed by network confidence, which remains almost entirely above the *>*0.9 threshold. The temporal difference loss does not detect these errors, due to two main issues: first, this loss spikes not only when the network jumps to a wrong location (frame 291), but also when it jumps back to the correct location (frame 292). Second, the temporal difference loss misses frames when the network lingers at the wrong location (frame 294). On the other hand, the multi-view PCA loss trace correctly utilizes the top-view prediction for this keypoint (white circle) to flag the error frames as inconsistent across views.

**Figure 3.**
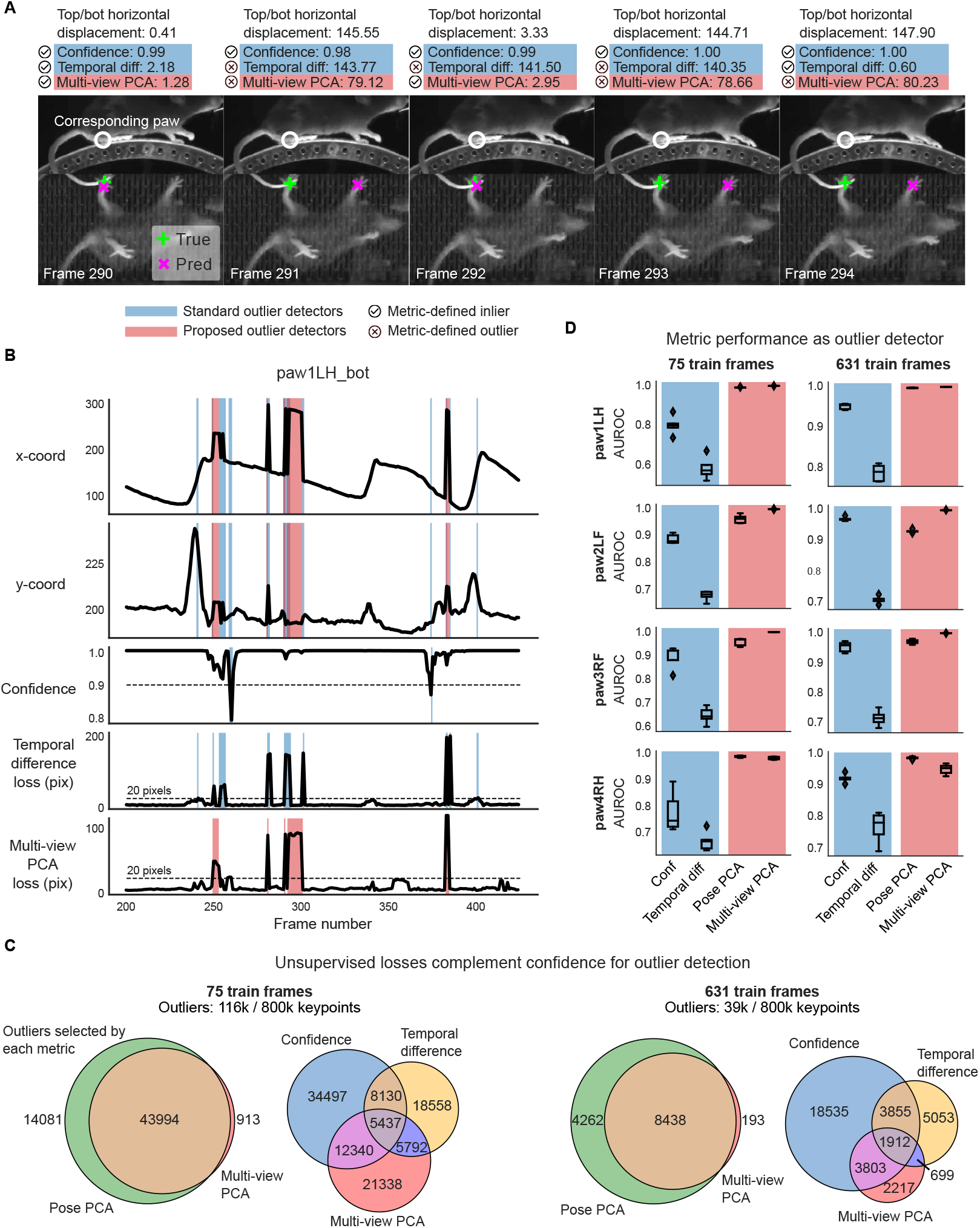
Unsupervised losses complement model confidence for outlier detection. **A**. Example frame sequence from the mirror-mouse dataset. Predictions from a DeepLabCut model (trained on 631 frames) are overlaid (magenta *×*), along with the ground truth (green +). Open white circles denote the location of the same body part (left hind paw) in the other (top) view; given the geometry of this setup, a large horizontal displacement between the top and bottom predictions indicates an error. Each frame is accompanied with “standard outlier detectors,” including confidence, temporal difference loss (shaded in blue), and “proposed outlier detectors,” including multi-view PCA loss (shaded in red; Pose PCA excluded for simplicity). 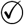 indicates an inlier as defined by each metric, and ⊗ indicates an outlier. Confidence is high for all frames shown, and the temporal difference loss misses error frame 294 which does not contain an immediate jump, and flags frame 292 which demonstrates a jump to the correct location. Multi-view PCA captures these correctly. **B**. Example traces from the same video. Blue background denotes times where standard outlier detection methods flag frames: confidence falls below a threshold (0.9) and/or the temporal difference loss exceeds a threshold (20 pixels). Red background indicates times where the multi-view PCA error exceeds a threshold (20 pixels). Purple background indicates both conditions are met. **C**. The total number of keypoints flagged as outliers by each metric, and their overlap. **D**. Area under the receiver operating characteristic curve (AUROC) for each paw, for DeepLabCut models trained with 75 and 631 labeled frames (left and right columns, respectively). AUROC=1 indicates the metric perfectly identifies all nominal outliers in the video data; 0.5 indicates random guessing. AUROC values are computed across all frames from 20 test videos; boxplot variability is over n= 5 random subsets of training data. Boxes use 25th/50th/75th percentiles for min/center/max; whiskers extend to 1.5 * IQR (inter-quartile range).

We proceed to generalize the example and quantify the overlaps and unique contributions of the different outlier detection methods on 20 unlabeled videos. We investigate two data regimes: “scarce labels” (75), which mimics prototyping a new tracking pipeline, and “abundant labels” (631 for the mirror-mouse dataset), i.e., a “production” setting with a fully trained network.

First, as we move from the scarce to the abundant labels regime, we find a 66% reduction in the outlier rate – the union of keypoints flagged by confidence, temporal difference, and multi-view PCA losses – going from 116k/800k to 39k/800k keypoints. This indicates that the networks become better and more confident. The Venn diagrams in Fig. 3C show that multi-view PCA captures a meaningful number of unique outliers which are missed by confidence and the temporal difference loss. (The Pose PCA includes both views and thus is largely overlapping with multi-view PCA.)

The overlap analysis above does not indicate which outliers are true versus false positives. To analyze this at a large scale, we restrict ourselves to a meaningful subset of the “true outliers” that can be detected automatically, namely predictions that are impossible given the mirrored geometry. We define this subset of outliers as frames for which the horizontal displacement between the top and bottom view predictions for a paw exceeds 20 pixels, similar to [16]; the networks output 72k/800k such errors with scarce labels, and 16k/800k with abundant labels. These spatial outliers should violate the PCA losses, but it is unknown whether they are associated with low confidence and large temporal differences. Instead of setting custom thresholds on our metrics as in Fig. 3B, we now estimate each metric’s sensitivity via a “Receiver Operating Characteristic” (ROC) curve, which plots the true positive rate against the false positive rate (both between 0 and 1), across *all* possible thresholds. The area under the ROC curve (AUROC) is a single measure summarizing the performance of each outlier detector: AUROC equals 1 for a perfect outlier detector, 0.5 for random guessing, and values below 0.5 indicate systematic errors. All metrics are above chance in detecting “true outliers” (Fig. 3D); for this class of spatial errors, the PCA losses are more sensitive outlier detectors than network confidence, and certainly more than the temporal difference loss (due to the pathologies described above).

To summarize, the PCA losses identify additional outliers that would have been otherwise missed by standard confidence and temporal difference thresholding (see Extended Data Fig. 1 and Extended Data Fig. 2 for similar results on mirror-fish and CRIM13 datasets). It is therefore advantageous to include PCA losses in standard outlier detection pipelines.

### 2.9 Both unsupervised losses and TCN boost tracking performance

Above we established that spatiotemporal constraint violations help identify network prediction errors. Next we quantify whether networks trained to avoid these constraint violations achieve more accurate and reliable tracking performance. As a “baseline” model for comparison, we implemented a supervised heatmap regression network that is identical to our more sophisticated model variants, but without semi-supervised learning or the TCN architecture. The baseline model matches DeepLabCut in performance across all datasets, though it is not intended to exactly match it in implementation (see Methods). The baseline model is useful because it eliminates implementation-level artifacts from model comparison. We quantify the networks’ performance both on an out-of-distribution labeled test set as well as on many unlabeled video frames. In this section, we compare the networks’ raw predictions, without any post-processing, to focally assess the implications of our architecture and unsupervised losses.

In Fig. 4A and Supplementary Video 5, we examine the mouse’s right hind paw position (top view) during two seconds of running. We compare the raw video predictions from our full semi-supervised model (in blue), including all supervised and unsupervised losses and a TCN architecture (henceforth, SS-TCN), to the predictions generated by our supervised baseline model, both trained on 75 labeled frames. The SS-TCN predictions are smoother (top two panels) and more confident (bottom panel), exhibiting a clearer periodic pattern expected for running on a stationary wheel. While some of the baseline model’s discontinuities are flagged by low confidence, some reflect a confident confusion between similar body parts, echoing Fig. 3A. One such confident confusion is highlighted in gray shading, and further scrutinized in Fig. 4B, showing that the baseline model (red) mistakenly switches to the left hind paw for two frames. The SS-TCN model avoids paw switching first because each frame is processed with its context frames, and second, because switching would have been heavily penalized by both the temporal difference loss and multi-view PCA loss.

**Figure 4.**
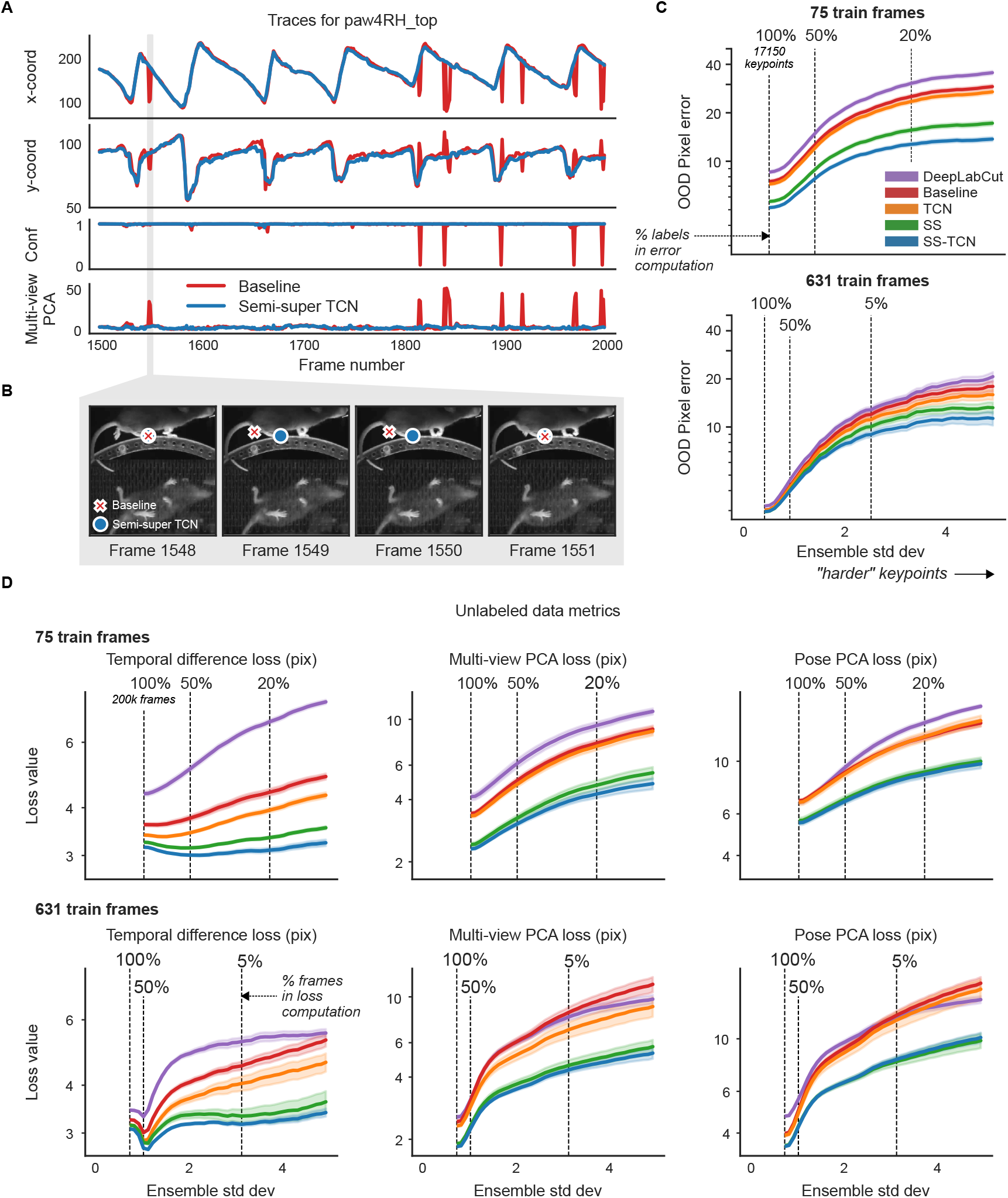
Unlabeled frames improve pose estimation (raw network predictions). **A**. Example traces from the baseline model and the semi-supervised TCN model (trained with 75 labeled frames) for a single keypoint (right hind paw; top view) on a held-out video (Supplementary Video 5). The semi-supervised TCN model is able to resolve the visible glitches in the trace, only some of which are flagged by the baseline model’s low confidence. One erroneous paw switch missed by confidence – but captured by multi-view PCA loss – is shaded in gray. **B**. A sequence of frames (1548-1551) corresponding to the gray shaded region in panel A in which a paw switch occurs. The estimates from both models are initially correct, then at Frame 1549 the baseline model prediction jumps to the incorrect paw, and stays there until it jumps back at Frame 1551. **C**. We compute the standard deviation of each keypoint prediction in each frame in the OOD labeled data across all model types and seeds (five random shuffles of training data). We then take the mean pixel error over all keypoints with a standard deviation larger than a threshold value, for each model type. Smaller standard deviation thresholds include more of the data (n=17150 keypoints total, indicated by the “100%” vertical line; (253 frames) *×* (5 seeds) *×* (14 keypoints) - missing labels), while larger standard deviation thresholds highlight more “difficult” keypoints. Error bands represent standard error of the mean over all included keypoints and frames for a given standard deviation threshold. **D**. Individual unsupervised loss terms are plotted as a function of ensemble standard deviation for the scarce (top) and abundant (bottom) label regimes. Error bands as in panel C, except we first compute the average loss over all keypoints in the frame (200k frames total; (40k frames) *×* (5 seeds)).

We perform an ablation study to isolate the contributions of our semi-supervised losses, our TCN architecture, and their combination. For each model type, we trained five networks with different random subsets of InD data. As noted by [22, 43], simple pixel error is an incomplete summary of network performance, since error averages may be dominated by a majority of “easy” keypoints, obscuring differences that may only be visible on the minority of “difficult” keypoints. We found it informative to quantify the pixel error as a function of keypoint difficulty, where we operationally define “difficulty” as the variance in the predictions of a deep ensemble of (five) networks (averaged across all model types). When this variance is large, at least one network in the ensemble must be in error; indeed, qualitatively, Fig. 1 and Supplementary Video 1 show that ensemble variance tends to increase on occlusion frames.

As expected, for both scarce and abundant label regimes (Fig. 4C), OOD pixel error increases as a function of ensemble standard deviation. With scarce labels, models struggle to resolve even “easy” keypoints, and SS-TCN outperforms baseline and DeepLabCut models across all levels of difficulty. The TCN architecture alone only mildly contributes to the improvements compared to semi-supervised learning in this dataset. By training semi-supervised models with a single loss at a time, we identify that multi-view and Pose PCA losses underlie most improvements (Extended Data Fig. 3). With abundant labels, all models accurately resolve “easy” keypoints, and the trends observed in the scarce labels regime become pronounced only for more “difficult” keypoints.

The above analysis was performed on a small set of 253 labeled OOD test frames. If we assess performance on a much larger unlabeled dataset of 20 OOD videos, and compute each of our losses for every predicted keypoint on every video frame, we observe similar trends (Fig. 4D): the SS-TCN model improves sample-efficiency with scarce labels, and reduces rare errors with abundant labels. (Recall that the semi-supervised models are explicitly trained to minimize these losses, and so these results on OOD data are consistent with expectations.)

We find similar patterns for the mirror-fish (Extended Data Fig. 4, Supplementary Video 6) and CRIM13 (Extended Data Fig. 5, Supplementary Video 7) datasets.

### 2.10 The Ensemble Kalman Smooter (EKS) enhances accuracy post-hoc

While our semi-supervised networks improve upon their fully-supervised counterparts, they still make mistakes. Recall that the spatiotemporal constraints are enforced during training but not at prediction time; therefore, we now present a post-processing algorithm which uses these constraints to refine the predictions. Successful post-processing requires identifying which predictions need fixing, that is, properly quantifying *uncertainty* for each keypoint on each frame. As emphasized above, low network confidence captures some, but not all, errors; conversely, constraint violations indicate the presence of additional errors within a set of keypoints but do not identify which specific keypoint within a given constraint violation is in fact an error.

Fig. 4C demonstrates that the *ensemble variance* – which varies for each keypoint on every frame – is an additional useful signal of model uncertainty [44, 45]. Hence, we developed a post-processing framework that integrates this ensemble variance uncertainty signal with our spatiotemporal constraints, via a probabilistic “state-space” model approach (Fig. 5A,B). Our model posits a latent “state” that evolves smoothly in time, and is projected onto the keypoint positions to enforce our spatial constraints. For example, we enforce multi-view constraints by projecting the three-dimensional true position of the body part (the “latent state”) through two-dimensional linear projections to obtain the keypoints in each camera view. This probabilistic state-space model corresponds to a Kalman filter-smoother model [46] and so we name the resulting post-processing approach the “Ensemble Kalman Smoother” (EKS) (see Methods).

**Figure 5.**
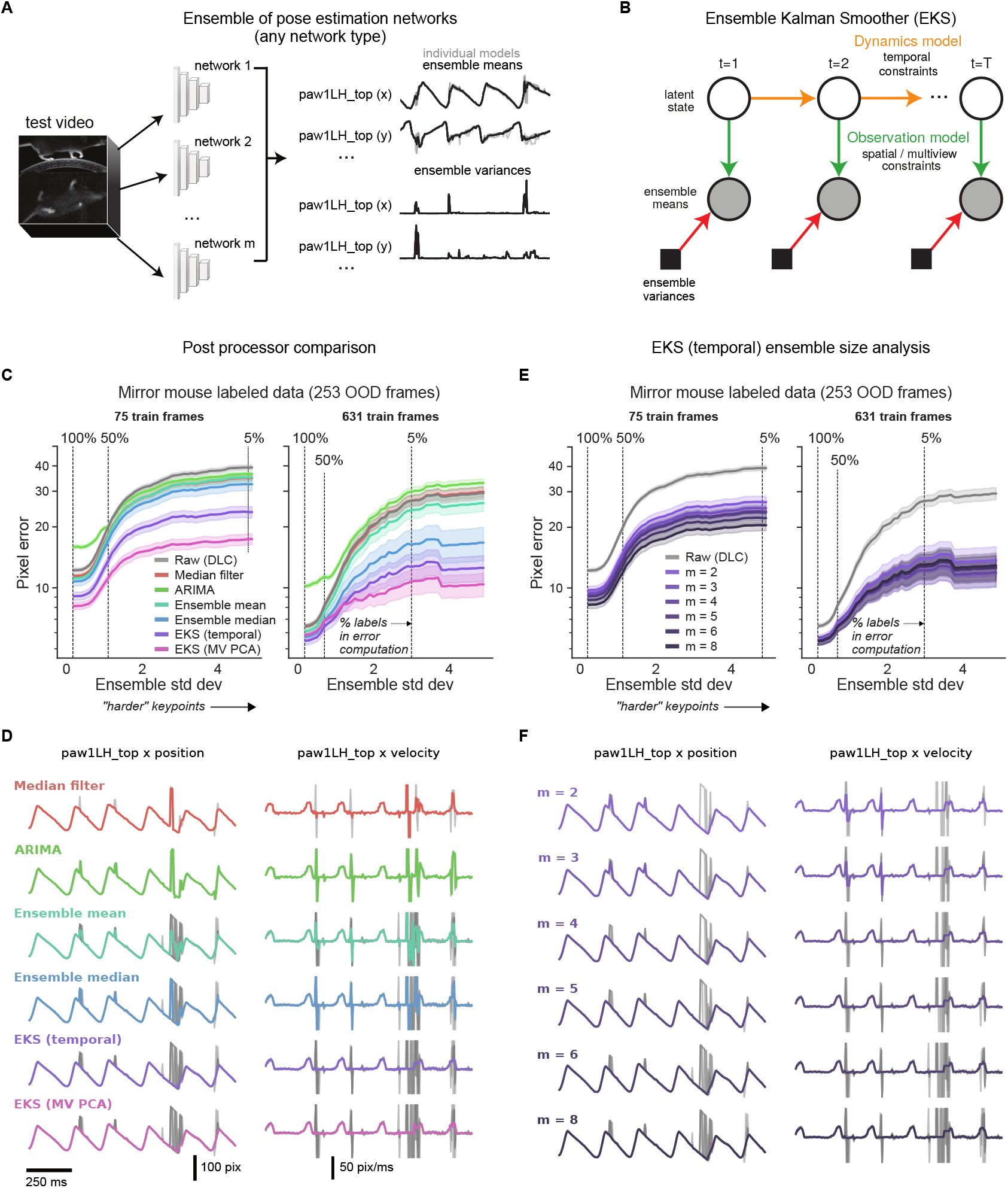
The Ensemble Kalman Smoother (EKS) post-processor. Results are based on DeepLabCut models trained with different subsets of InD data and different random initializations of the head. **A**. *Deep ensembling* combines the predictions of multiple networks. The *ensemble mean* is potentially more accurate than single model predictions, and the *ensemble variance* can be a useful measure of uncertainty that is complementary to single model confidence values. **B**. EKS leverages the spatiotemporal constraints of the unsupervised losses as well as uncertainty measures from the ensemble variance in a probabilistic state-space model. Ensemble means of the keypoints are modeled with a latent linear dynamical system; temporal smoothness constraints are enforced through linear dynamics (yellow arrows) and spatial constraints (Pose or multi-view PCA) are enforced through a fixed observation model that maps the latent state to the observations (green arrows). Instead of learning the observation noise, we use the time-varying ensemble variance (red arrows). EKS uses a Bayesian approach to weight the relative contributions from the prior and observations. **C**. Post-processor comparison on OOD frames from the mirror-mouse dataset. We plot pixel error as a function of ensemble standard deviation (as in Fig. 4). The median filter and ARIMA models act on the outputs of single networks; the ensemble means, ensemble medians, and EKS variants act on an ensemble of five networks. EKS (temporal) only utilizes temporal smoothness, and is applied one keypoint at a time. EKS (MV PCA) utilizes multiview information as well as temporal smoothness, and is applied one body part at a time (tracked by one keypoint in each of two views). Error bands as in Fig. 4 (n=17150 keypoints at 100% line). **D**. Trace comparisons for different methods (75 train frames). Gray lines show the raw traces used as input to the method colored lines show the post-processed trace. **E**. Pixel error comparison for the EKS (temporal) post-processor as a function of ensemble members (m). Error bands as in panel C. **F**. Trace comparisons for varying numbers of ensemble members (75 train frames).

The EKS model output represents a Bayesian compromise between the spatiotemporal constraints (prior) and the information provided by the ensemble observations (likelihood). Concretely, if a keypoint’s uncertainty is low (i.e., all ensemble members agree) then this observation will be upweighted relative to the spatiotemporal prior and will only be lightly smoothed. Conversely, when a keypoint’s uncertainty is high, the spatiotemporal priors and other more-confident keypoints’ predictions will be used to interpolate over these uncertain observations. Unlike previous approaches [16, 20, 22, 32, 42], EKS requires no manual selection of confidence thresholds or (suboptimal) temporal linear interpolation separately for each dropped keypoint. Moreover, the EKS post-processing approach is agnostic to the type of networks used to generate the ensemble predictions.

We benchmark EKS on DeepLabCut models fit to the mirror-mouse dataset. EKS compares favorably to other standard post-processors, including median filters and ARIMA models (which are fit on the outputs of single networks), and the ensemble mean and median (computed using an ensemble of multiple networks; Fig. 5C,D). EKS provides substantial improvements in OOD pixel errors with as few as *m* = 2 networks; we find *m* = 5 networks is a reasonable choice given the computation-accuracy tradeoff (Fig. 5E,F), and use this ensemble size throughout.

When applied to Lightning Pose semi-supervised TCN models, EKS provides additional improvements across multiple datasets, particularly on “difficult” keypoints where the ensemble variance is higher (Extended Data Fig. 6). EKS achieves smooth and accurate tracking even when the models make errors due to occlusion and paw confusion (Extended Data Fig. 6; Supplementary Video 8-12; Supplementary Figs. 2-4).

### 2.11 Improved tracking on International Brain Laboratory datasets

Next we turn to an analysis of large-scale public datasets from the International Brain Laboratory (IBL) [30]. In each experimental session, a mouse was observed by three cameras while performing a visually-guided decision-making task. The mouse signaled its decisions by manually moving a rotary wheel left or right. We analyze two of IBL’s video datasets. “IBL-pupil” contains zoomed-in videos of the pupil, where we track the top, bottom, left, and right edges of the pupil. In “IBL-paw” we track the left and right paws.

Despite efforts at standardization, the data exhibit considerable visual variability between sessions and labs, which presents serious challenges to existing pose estimation methods. Specifically, in IBL’s preliminary data release we used DeepLabCut, followed by custom post-processing. As detailed in [30], this approach fails in a majority of pupil recordings: the signal-to-noise ratio of the estimated pupil diameter is too low for reliable downstream use, largely due to occlusions caused by whisking and infrared light reflections. Paw tracking tends to be more accurate, but is contaminated by discontinuities especially when a paw is retracted behind the torso. In this section, we report the results for IBL-pupil. The IBL-paw results appear in Extended Data Fig. 7 and the Supplementary Information.

In Fig. 6, we evaluate three pose estimators: DeepLabCut with custom post-processing (DLC; left column of example session), Lightning Pose’s semi-supervised TCN model with the same post-processing (LP; middle column, using temporal difference and Pose PCA losses), and the pupil-specific EKS variant applied to an ensemble of *m* = 5 LP models (LP+EKS; right column). The pupil-specific EKS uses a threedimensional latent state: pupil centroid (width and height coordinates) and a diameter. (It is straightforward to construct an EKS version with separate horizontal and vertical diameters, but we found this extension to be unnecessary for this dataset.) The latent state is then projected linearly onto the eight-dimensional tracked pixel coordinates (width and height for top, bottom, left, and right edges; see Methods).

**Figure 6.**
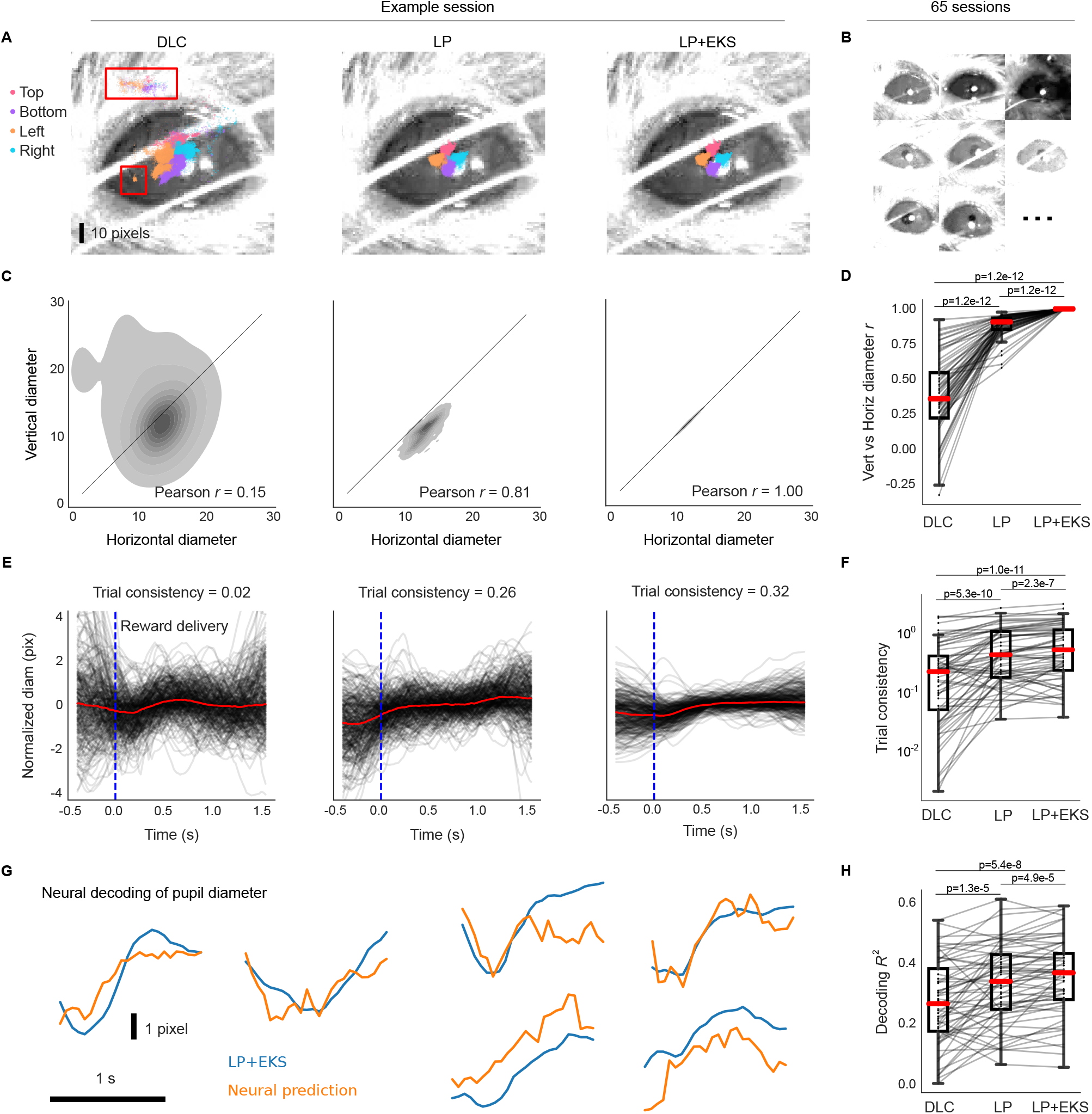
Lightning Pose models and EKS improve pose estimation on IBL pupil data. **A**. Sample frame overlaid with a subset of pupil markers estimated from DeepLabCut (DLC; *left*), Lightning Pose using a semi-supervised TCN model (LP; *center*), and a 5-member ensemble using semi-supervised TCN models (LP+EKS; *right*). **B**. Example frames from a subset of 65 IBL sessions, illustrating the diversity of imaging conditions in the dataset. **C**. Empirical distribution of vertical diameter measured from top and bottom markers scattered against horizontal pupil diameter measured from left and right markers. These estimates should ideally be equal, i.e., the distribution should lie near the diagonal line. Column arrangement as in panel A. The LP+EKS estimate imposes a low-dimensional model that enforces perfectly correlated vertical and horizontal diameters by construction. **D**. Vertical vs horizontal diameter correlation is computed across n=65 sessions for each model. The LP+EKS model has a correlation of 1.0 by construction. **E**. Pupil diameter is plotted for correct trials aligned to feedback onset; each trial is mean-subtracted. DeepLabCut and LP diameters are smoothed using IBL default post-processing (Methods), compared to LP+EKS outputs. We compute a *trial consistency* metric (the variance explained by the mean over trials; see text) as indicated in the panel titles. See Supplementary Video 13. **F**. The trial consistency metric computed across n=65 sessions. **G**. Example traces of LP+EKS pupil diameters (blue) and predictions from neural activity (orange) for several trials using cross-validated, regularized linear regression (Methods). **H**. Neural decoding performance across n=65 sessions. Panels D, F, and H use a one-sided Wilcoxon signed-rank test; boxes use 25th/50th/75th percentiles for min/center/max, and whiskers extend to 1.5 * IQR. See Supplementary Table 2 and main text for further quantification of boxes.

To directly compare our methods to the publicly released IBL DeepLabCut traces, we train on all available data and evaluate on held-out unlabeled videos. We define several pupil-specific metrics to quantify the accuracy of the different models and their utility for downstream analyses.

The first metric compares the “vertical” and “horizontal” diameters, i.e., top(*y*) - bottom(*y*) and right(*x*) - left(*x*), respectively. The vertical and horizontal diameters should be equal (or at least highly correlated) and, therefore, low correlations between these two values signal poor tracking. We compute this correlation in an example session in Fig. 6C, and over 65 sessions in Fig. 6D. The LP model (Pearson’s *r*=0.88*±*0.01, mean*±*sem) improves over the DeepLabCut model (*r*=0.36*±*0.03). (Since the pupil-specific EKS uses a single value for both vertical and horizontal diameters, it enforces a correlation of 1.0 by construction.)

Scientifically, we are interested in how behaviorally-relevant events (such as reward onset) impact pupil dynamics, as well as the correlation between pupil dynamics and neural activity. We expect that noise in our estimates of pupil diameter would reduce the apparent consistency of pupil dynamics across trials and also reduce any correlations between pupil diameter and neural activity; this is exactly what we observe (Fig. 6E-H). In Fig. 6E, we align diameter estimates across multiple trials to the time of reward delivered at the end of each successful trial. We define a second quality metric – trial consistency – by taking the variance of the mean pupil diameter trace and dividing by the variance of the mean-subtracted traces across all trials. This metric is zero if there are no reproducible dynamics across trials; it is infinity if the pupil dynamics are identical and non-constant across trials (constant outputs will result in an undefined metric since both numerator and denominator are zero). Although we expect some amount of real trial-to-trial variability in pupil dynamics, any noise introduced during pose estimation will decrease this metric. The LP and LP+EKS estimates show greater trial-to-trial consistency compared to the DeepLabCut estimates, both within a single session (Fig. 6E) and across multiple sessions (Fig. 6F; DeepLabCut 0.35*±*0.06; LP 0.62*±*0.07; LP+EKS 0.74*±*0.08). Supplementary Video 13 shows pupil diameter traces in multiple trials, and demonstrates that the increased trial-to-trial consistency does not compromise the model’s ability to track the pupil well within individual trials.

After establishing the improved reliability of the pupil diameter signal, we now examine whether it is more correlated with neural data. This analysis serves to verify that the LP+EKS approach is not merely suppressing pupil diameter fluctuations, but rather better capturing pupil signals that can be predicted from an independent measurement of neural activity. We compute the accuracy of a ridge regression model from neural data to pupil diameter (*R*^2^ on held-out trials; see Methods). Fig. 6H shows that across sessions, LP and LP+EKS enhance decoding accuracy compared to DeepLabCut (DLC *R*^2^=0.27*±*0.02; LP 0.33*±*0.02; LP+EKS 0.35*±*0.02).

### 2.12 The Lightning Pose software package and a cloud application

We open-source a flexible software package and an easy-to-use cloud application. Both are presented in detail in the Methods, and we provide a brief overview here.

Extended Data Fig. 8A illustrates the design principles of the Lightning Pose package, which we built to be 1) *simple to use and easy to maintain*: we aim to minimize “boilerplate” code (such as basic GUI development or training loggers) by outsourcing to industry-grade packages; 2) *video-centric*: the networks operate on video clips, rather than on a single image at a time; 3) *modular and extensible*: our goal is to facilitate prototyping of new losses and models; 4) *scalable*: we support efficient semi-supervised training and evaluation; 5) *interactive*: we offer a variety of tracking performance metrics and visualizations during and after training, enabling easy model comparison and outlier detection.

The scientific adoption of deep learning packages like ours presents an infrastructure challenge. Labs need access to GPU-accelerated hardware with a set of pre-installed drivers and packages. Often, developers spend more time in hardware setup and installation than configuring, training, and evaluating pose tracking models (see e.g. [25] for further discussion). We therefore developed a cloud application that supports the full life cycle of animal pose estimation (Extended Data Fig. 8B), and is suitable for users with minimal coding expertise and only requires internet access.

## 3 Discussion

We presented Lightning Pose, a semi-supervised deep learning system for animal pose estimation. Lightning Pose uses a set of spatiotemporal constraints on postural dynamics to improve network reliability and efficiency. We further refine the pose estimates post-hoc, with the Ensemble Kalman Smoother (EKS) that uses reliable predictions and spatiotemporal constraints to interpolate over unreliable ones.

Our work builds on previous semi-supervised animal pose estimation algorithms that use spatiotemporal losses on unlabeled videos [34, 37]. Semi-supervised learning is not the only technique that enables improvements over standard supervised learning protocols. First, it has been suggested that supervised pose estimation networks can be improved by pretraining them on large labeled datasets for image classification

[9] or pose estimation [47], to an extent that might eliminate dataset-specific training [48]. Other work avoids pretraining altogether by using lighter architectures [10]. These ideas are complementary to ours: any robust backbone obtained through these procedures could be easily integrated into Lightning Pose, and further refined via semi-supervised learning.

Human pose estimation, like animal pose estimation, is most commonly approached using supervised heatmap regression on a frame-by-frame basis [49]. Unlike the animal setting, human models are trained on much larger labeled datasets containing either annotated images [50] or 3D motion capture [51]. Moreover, human models track a standardized set of keypoints, and some operate on a standard skinned human body model [52]. In contrast, most animal pose estimation must contend with relatively scarce labels, lower quality videos, and bespoke sets of labels to track, varying by species and lab. Though human pose estimation models can impressively track crowds of moving humans, doing downstream science using the keypoints still presents several challenges [49] similar to those discussed in the Results. Lightning Pose can be applied to single-human pose estimation, by fine-tuning an off-the-shelf human pose estimation backbone to specific experimental setups (such as patients in a clinic), while enforcing our spatiotemporal constraints (or new ones). Future work could also apply EKS to the outputs of off-the-shelf human trackers.

Roughly speaking, two camps coexist in multi-view animal pose estimation [53]: those who use 3D information during training [12, 37, 54, 55] and those who train standard 2D networks and perform 3D reconstruction post-hoc [20, 56]. Either approach involves camera calibration, whose limitations we discussed above. Lightning Pose can be seen as an intermediate approach: we train with 3D constraints without an explicit camera calibration step (however, our current approach assumes the cameras have no distortion). At the same time, Lightning Pose does not provide an exact 3D reconstruction of the animal, but rather a scaled, rotated and shifted version thereof. Our improved predictions could be readily used as inputs to existing 3D reconstruction pipelines. Some authors employed 3D convolutional networks that operate on 3D location voxels instead of 2D heatmaps [12]. These architectures have been recently trained in a semi-supervised fashion with temporal constraints [43] akin to to [34] and the current work. We note that, although 3D voxels are more computationally expensive than 2D heatmaps, they could be incorporated as strong backbones for Lightning Pose.

A number of additional important directions remain for future work. One involves implementing richer models of moving bodies as losses. Multiple approaches have been recently proposed for modeling pose trajectories post-hoc. These include probabilistic body models [21, 57, 58], mechanical models [59], switching linear dynamical systems [19, 34, 60], and autoencoders [20]. These models could be made even more effective by being integrated into network training in a so-called end-to-end manner. Any model of pose dynamics, as long as it is differentiable, could be incorporated as an unsupervised loss.

Another important direction is to improve the efficiency of the EKS method. The advantages of ensembling come at a cost: we need to train, store, and run inference with multiple networks. (Post-processing the networks’ output with EKS is relatively computationally cheap.) One natural approach would be *knowledge distillation* [61]: train a single network to emulate the full EKS output.

Finally, while the methods proposed here can currently track multiple distinguishable animals (e.g., a black mouse and a white mouse), they do not apply directly to multi-animal tracking problems involving multiple similar animals [18, 62], since to compute our unsupervised losses we need to be able to know which keypoint belongs to which animal. Thus adapting our approaches to the general multi-animal setting remains an important open avenue for future work.

## Acknowledgments

We thank Peter Dayan and Nick Steinmetz for serving on our IBL paper board, as well as two anonymous reviewers whose detailed comments considerably strengthened our manuscript. We are grateful to Natalie Biderman for productive discussions and help with visualization. We thank Matteo Carandini and Jacob Portes for helpful comments; Taiga Abe, Kelly Buchanan, and Geoff Pleiss for helpful discussions on ensembling; and Haotian Xiang for conversations on active learning and outlier detection. Thanks to William Falcon, Luca Antiga, Thomas Chaton and Adrian Wälchi (Lightning AI) for their technical support and advice on implementing our package and the cloud application. This work was supported by the following grants: Gatsby Charitable Foundation GAT3708 (DB, MRW, CH, NRG, AV, JPC, LP), German National Academy of Sciences Leopoldina (AEU), Irma T Hirschl Trust (NBS), Netherlands Organisation for Scientific Research (VI.Veni.212.184) (AEU), NSF IOS-2115007 (NBS), NIH K99NS128075 (JPN), NIH NS075023 (NBS), NIH U19NS123716 (MRW), NSF 1707398 (DB, MRW, CH, NRG, AV, JPC, LP), Simons Foundation (MRW, MS, JMH, AK, GTM, JPN, APV, KZS), Wellcome

Trust 216324 (MS, MS, JMH, AK, GTM, JPN, APV, KZS). The funders had no role in study design, data collection and analysis, decision to publish or preparation of the manuscript.

## Author Contributions Statement

Conceptualization: DB, MRW, LP; Software Package — core development: DB, MRW, NRG; Software Package — contribution: CH, AV; Cloud Application — development: MRW, DB, RL, AV; First draft — writing: DB, MRW, LP; First draft — editing: DB, MRW, CH, LP; Data collection: DB, MS, JMH, AK, GTM, JPN, APV, KZS, AEU, RW, DN, FP; Funding — JPC, NS, LP; Semi-supervised learning algorithms: DB, MRW, NRG, LP; Deep ensembling: DB, MRW, CH, LP; Ensemble Kalman Smoothing: CH, LP; Temporal Context Network: CH, DB, MRW, LP; Diagnostic tools and visualization: DB, MRW, AV; Neural network experiments and analysis: DB, MRW.

## 4 Methods

All datasets used for the experiment were collected in compliance with the relevant ethical regulations. See the following published papers for each dataset: mirror-mouse: [1], CRIM13: [2], and IBL datasets: [3]. All mirror-fish experiments adhere to the American Physiological Society’s Guiding Principles in the Care and Use of Animals and were approved by the Institutional Animal Care and Use Committee of Columbia University, protocol number AABN0557.

### 4.1 Datasets

We consider diverse datasets collected via different experimental paradigms for mice and fish. For each dataset, we collected a large number of videos including different animals and experimental sessions, and labeled a subset of frames from each video. We then split this data into two non-overlapping subsets (i.e., a given animal and/or session would appear only in one subset). The first subset is the “in-distribution” (InD) data that we use for model training. The second subset is the “out-of-distribution” (OOD) data that we use for model evaluation. This setup mimics the common scenario in which a network is thoroughly trained on one cohort of subjects, and is then used to predict new subjects. Supplementary Table 1 details the number of frames for each subset per dataset, as well as the number of unique animals and videos those frames came from.

### Mirror-mouse

Head-fixed mice ran on a circular treadmill while avoiding a moving obstacle [1]. The treadmill had a transparent floor and a mirror mounted inside at 45^*°*^, allowing a single camera to capture two roughly orthogonal views (side view and bottom view via the mirror) at 250 Hz. The camera was positioned at a large distance from the subject (*∼* 1.1 meters) to minimize perspective distortion. Frames are (406*×*396) and reshaped during training to (256*×*256). 17 keypoints were labeled across the two views including seven keypoints on the mouse’s body per view, plus three keypoints on the moving obstacle.

### Mirror-fish

19 wild-caught (age unknown) adult male and female Mormyrid fish (15-22 cm in length) of the species *Gnathonemus petersii* were used in the experiment. Fish were housed in 60-gallon tanks in groups of 5-20. Water conductivity was maintained between 60-100 microsiemens both in the fish’s home tanks and during experiments.

The fish swam freely in and out of an experimental tank, capturing worms from a well. The tank had a side mirror and a top mirror, both at 45^*°*^, providing three different views seen from a single camera at 300 Hz (Supplementary Fig. 1). Here too, the camera was placed *∼* 1.7 meters away from the center of the fish tank to reduce distortions. Frames are (384 *×* 512) and reshaped during training to (256 *×* 384).

17 body parts were tracked across all three views for a total of 51 keypoints. We pre-processed the labeled dataset as follows. First, we identified labeling errors by flagging large values of the multi-view PCA loss. We then fixed the wrong labels manually. Next, in the InD data only, we used a probabilistic variant of multi-view PCA (PPCA) to infer keypoints that were occluded in one out of the three views, effectively similar to the triangulation-reprojection protocols used for multi-view tracking by [4, 5]. This resulted in a 30% increase in the number of keypoints usable for training, with more occluded keypoints included in the augmented label set.

### CRIM13

The Caltech Resident-Intruder Mouse dataset (CRIM13) [2] consists of two mice interacting in an enclosed arena, captured by top and side view cameras at 30 Hz. We only use the top view. Frames are (480 *×* 640) and reshaped during training to (256 *×* 256). Seven keypoints were labeled on each mouse for a total of 14 keypoints.

Unlike the other datasets, the InD/OOD splits do not contain completely non-overlapping sets of animals, as we used the train/test split provided in the dataset. The (4) resident mice are present in both InD and OOD splits; however, the intruder mouse is different for each session. Each keypoint in the CRIM13 dataset is labeled by five different annotators. To create the final set of labels for network training, we took the median across all labels for each keypoint.

### IBL-paw

This dataset [3] comes from the International Brain Lab and consists of head-fixed mice performing a decision-making task [6, 7]. Two cameras – “left” (60 Hz) and “right” (150 Hz) – capture roughly orthogonal side views of the mouse’s face and upper trunk during each session. The original dataset does not contain synchronized labeled frames for both cameras, preventing the direct use of multi-view PCA losses during training. Instead, we treat the frames as coming from a single camera by flipping the right camera video. Frames were initially downsampled to (102 *×* 128) for labeling and video storage; frames were reshaped during training to (128 *×* 128). We tracked two keypoints per view, one for each paw. More information on the IBL video processing pipeline can be found in [8]. For the large scale analysis in Extended Data Fig. 7 we selected 44 additional test sessions that were not represented in the InD or OOD sessions listed in Supplementary Table 1; these could be considered additional OOD data.

### IBL-pupil

The pupil dataset also comes from the International Brain Lab. Frames from the right camera were spatially upsampled and flipped to match the left camera. Then, a 100 *×* 100 pixel ROI was cropped around the pupil. The frames were reshaped in training to (128 *×* 128). Four keypoints were tracked on the top, bottom, left and right edges of the pupil, forming a diamond shape. For the large scale analysis in Fig. 6 we selected left videos from 65 additional sessions that were not represented in the InD or OOD sessions listed in Supplementary Table 1.

### 4.2 Problem formulation

Let *K* denote the number of keypoints to be tracked, and *N* the number of labeled frames. After manual labeling, we are given a dataset:

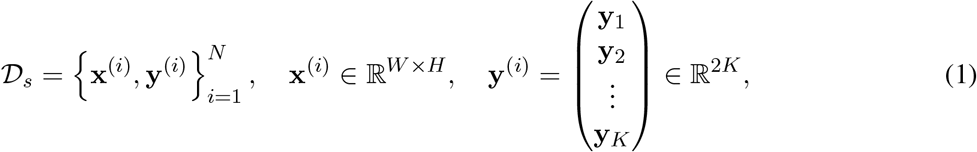

where **x**^(*i*)^ is the *i*-th image and **y**^(*i*)^ its associated label vector, stacking the annotated width-height pixel coordinates for each of the *K* tracked keypoints.

It is standard practice to represent each annotated keypoint **y**_*k*_, *k* = 1, … *K* as a heatmap 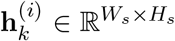 with width *W*_*s*_ and height *H*_*s*_, thus converting **y**^(*i*)^ to a set of *K* heatmaps 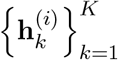. This is done by defining a bivariate Gaussian centered at each annotated keypoint with variance *σ*^2^ (a controllable parameter), and evaluating it at 2D grid points (for more details, see [9]). If 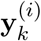 lacks an annotation (e.g. if it is occluded), we do not form a heatmap for it.

We normalize the heatmaps 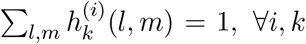, which allows us to both evenly scale the outputs during training and use losses that operate on heatmaps as valid probability mass functions. Then, the dataset for training supervised networks is just frames and heatmaps 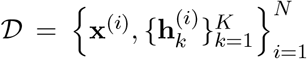. To accelerate training, the heatmaps are made 4 or 8 times smaller than the original frames.

### 4.3 Model architectures

#### 4.3.1 Baseline

Our baseline model performs heatmap regression on a frame-by-frame basis, akin to DeepLabCut [10], SLEAP [9], DeepPoseKit [11], and others. It has roughly the same architecture: a “backbone” network that extract a feature vector per frame, and a “head” that transforms these into *K* predicted heatmaps. In the results reported here, we use a ResNet-50 backbone network pretrained on the AnimalPose10K dataset ([12]; 10,015 annotated frames from 54 different animal species). For the mirror-fish dataset, we rely on ImageNet pretraining (except for the sample efficiency experiments in Fig. 1). However, our package, like others, is largely agnostic to backbone choices. Let *B* denote batch size, *C* = 3 the RGB color channels, and *r* an “upscaling factor” by which we increase the size of our representations. The head includes a fixed PixelShuffle(2) layer that reshapes the features tensor output by the backbone from (*B, C × r*^2^, *H, W*) to (*B, C, H × r, W × r*) and a series of identical ConvTranspose2D layers that further double it in size (kernel size (3 *×* 3), stride (2 *×* 2), input padding (1 *×* 1), and output padding (1 *×* 1)) [13]. The number of ConvTranspose2D layers is determined by the desired shape of the output heatmaps, and most commonly two such layers are used. Each heatmap is normalized with a 2D spatial softmax with a temperature parameter *τ* = 1. The supervised loss is a divergence between predicted heatmaps and labeled heatmaps. Here, we use squared error for each batch element *b* and keypoint *k*: 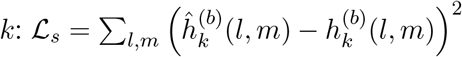.

Once heatmaps have been predicted for each keypoint, we must transform these 2D arrays into estimates of the width-height coordinates in the original image space. We first upsample each heatmap 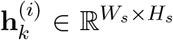 to 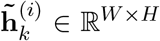 using bicubic interpolation. We then compute a subpixel maximum akin to DeepPoseKit [11]. A 2D spatial softmax renormalizes the heatmap to sum to 1, and we apply a high temperature parameter (*τ* = 1000) to suppress non-global maxima. A 2D spatial expectation then produces a subpixel estimate of the location of the heatmap’s maximum value. These two operations – spatial softmax followed by spatial expectation – are together known as a soft argmax [14]. Importantly, this soft argmax operation is differentiable (unlike the location refinement strategy employed in [10]), and allows the estimated coordinates to be used in downstream losses. To compute the confidence value associated with the pixel coordinates, we sum the values of the normalized heatmap within a configurable radius of the soft argmax.

#### 4.3.2 Temporal Context Network

Many detection ambiguities and occlusions in a given frame can be resolved by considering some video frames before and after it. The Temporal Context Network (TCN) uses a sequence of 2*J* + 1 frames to predict the labeled heatmaps for the middle frame,

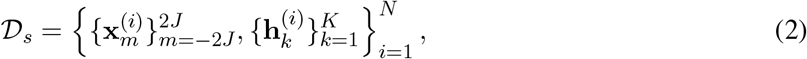

Where 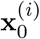 is the labeled frame and, for example, 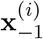 is the preceding (unlabeled) frame in the video.

During training, batches of 2*J* +1 frame sequences are passed through the backbone to obtain 2*J* +1 feature vectors. The TCN has two upsampling heads, one “static” and one “context-aware,” each identical to the baseline model’s head. The static head takes the features of only the central frame and predicts location heatmaps for that frame. The context-aware head generates predicted location heatmaps for each of the 2*J* + 1 frames (note, these are the same shape as the location heatmaps, but we do not explicitly enforce them to match labeled heatmaps). Those heatmaps are passed as inputs to a bi-directional convolutional recurrent neural network whose output is the context-aware predicted heatmap for the middle frame. We then apply our supervised loss to both predicted heatmaps, forcing the network to learn the standard static mapping from an image to heatmaps, while independently learning to take advantage of temporal context when needed. (Recall Fig. 2E, which provides an overview of this architecture.)

The network described above outputs two predicted heatmaps per keypoint, one from each head, and applies the computations described above to obtain two sets of keypoint predictions with confidences. For each keypoint, the more confident prediction of the two is selected for downstream analysis.

### 4.4 Semi-supervised learning

We perform semi-supervised learning by jointly training on labeled dataset *𝒟*_*s*_ (constructed as described above) and an unlabeled dataset *𝒟*_*u*_:

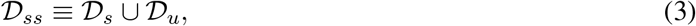

where *𝒟*_*u*_ is constructed as follows.

Assume we have access to one or more unlabeled videos; we splice these into a set of *U* disjoint *T* -frame clips (discarding the very last clip if it has fewer than *T* frames),

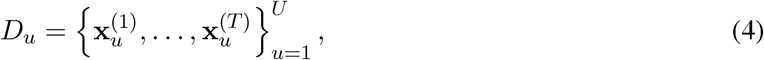

where, typically, *T* = 32*/*64*/*96*/*128*/*256 with with smaller frame sizes freeing up memory for longer sequences.

Now, assume we selected a mechanism (baseline model or TCN) for predicting keypoint heatmaps for a given frame. At each training step, in addition to a batch of labeled frames drawn from *𝒟*_*s*_, we present the network with a short unlabeled video clip randomly drawn from *𝒟*_*u*_. The network outputs a time-series of keypoint predictions (one pose for each of the *T* frames in the clip), which is then subjected to one or more of our unsupervised losses.

All unsupervised losses are expressed as pixel distance between a keypoint prediction and the constraint. Since our constraints are merely useful approximate models of reality, we do not require the network to perfectly satisfy them. We are particularly interested in preventing, and having the network learn from, severe violations of these constraints. Therefore, we enforce our losses only when they exceed a tolerance threshold *ϵ*, rendering them *ϵ*-insensitive:

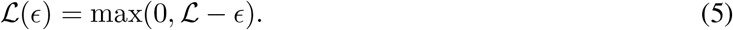

The *ϵ* threshold could be chosen using prior knowledge, or estimated empirically from the labeled data, as we will demonstrate below. *ℒ*(*ϵ*) is computed separately for each keypoint on each frame, and averaged to obtain a scalar loss to be minimized. Multiple losses can be jointly minimized via a weighted sum, with weights determined by a parallel hyperparameter search, which is supported in Lightning Pose with no code changes.

#### 4.4.1 Temporal difference loss

Keypoints should not jump too far between consecutive frames. We measure the jump in pixels and ignore jumps smaller than *ϵ*, the maximum jump allowed by user,

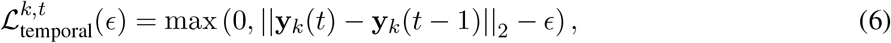

where *ϵ* could be determined based on image size, frame rate, and rough viewing distance from the subject. We compute this loss for a pair of successive predictions only when both have confidence greater than a configurable threshold (e.g. 0.9) to avoid artificially enforcing smoothness in stretches where the keypoint is unseen. We average the loss across keypoints and unlabeled frames:

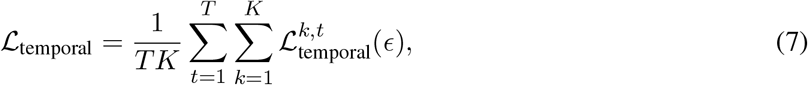

and minimize *ℒ*_temporal_ during training. Lightning Pose also offers the option to apply the temporal difference loss on predicted heatmaps instead of the keypoints. We have found both methods comparable and focus on the latter for clarity.

#### 4.4.2 Multi-view PCA loss

### Background

Let 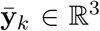 be an unknown 3D keypoint of interest. Assume that we have *V* cameras and that each *v* = 1, …, *V* camera sees a single 2D perspective projection of 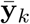 denoted as **y**_*k*_(*v*) *∈ ℛ*^2^, in pixel coordinates. (Note that is standard to express 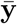 and **y**(*v*) in “homogeneous coordinates”, i.e., appending another element to each vector, yet we omit this for simplicity and for a clearer connection with our PCA approach.) Thus, we have a 2*V* -dimensional measurement (**y**_*k*_(1)^*T*^ *· · ·* **y**_*k*_(*V*)^*T*^) of our 3D keypoint 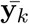.

### The multi-view geometry approach

It is standard to model each view as a pinhole camera [15]: such a camera has intrinsic parameters (focal length and distortion) and extrinsic parameters (its 3D location and orientation, a.k.a “camera pose”), that together specify where a 3D keypoint will land on its imaging plane, i.e., the transformation from 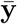 to **y**(*v*). This transformation involves a linear projection (scaling, rotation, translation) followed by a nonlinear distortion. While one might know a camera’s focal length and distortion, in general, both the intrinsic and extrinsic parameters are not exactly known and have to be estimated. A standard way to estimate these involves “calibrating” the camera; filming objects with groundtruth 3D coordinates, and measuring their 2D pixel coordinates on the camera’s imaging plane. Physical checkerboards are typically used for this purpose. They have known patterns that can be presented to the camera and detected using traditional computer vision techniques. Now with a sufficient set of 3D inputs and 2D outputs, the intrinsic and extrinsic parameters can be estimated via (nonlinear) optimization.

### Multi-view PCA on the labels (our approach)

We take a simpler approach which does not require camera calibration or, in the mirrored datasets considered in this paper, explicit information about the location of the mirrors. Our first insight is that the multi-view (2*V* -dimensional) labeled keypoints could be used as key-point correspondences to learn the geometric relationship between the views. We approximate the pinhole camera as a linear projection (with zero distortion), and estimate the parameters of this linear projection by fitting PCA on the labels (details below), and keeping the first three PCs, since all we are measuring from our different cameras is a single 3D object. Fig. 2C (bottom right) confirms that our PCA model can explain *>* 99% of the variance with the first three PCs in several multi-view experimental setups, indicating that our linear approximation is suitable at least for the mirror-mouse and mirror-fish datasets, in which the camera is relatively far from the subject. We do anticipate cases where our linear approximation will not be sufficiently accurate (e.g., strongly distorted lenses, or highly zoomed in); the more general epipolar geometry approach of [16, 17] could be applicable here. Note that our three-dimensional PCA coordinates do not exactly match the 3D width-height-depth physical coordinates of the keypoints in space; instead, these two sets of three-dimensional coordinates are related via an affine transformation.

### Before training: fitting multi-view Principal Component Analysis (PCA) on the labels

Our goal is to estimate a projection from 2*V* dimensions (width-height pixel coordinates for *V* views) to three dimensions, which we could use to relate the different views to each other. Given the indices of matching keypoints across views, we form a tall and thin design matrix by vertically stacking all the 2*V* -dimensional multi-view labeled keypoints. We denote this matrix as **Y**_*MV*_ *∈* ℛ^*NK×*2*V*^,

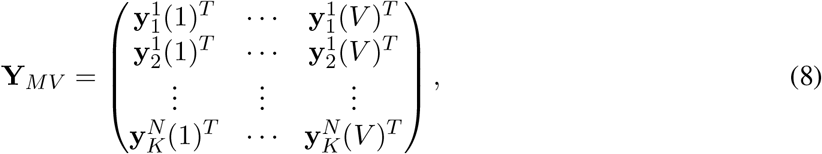

Where 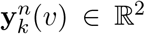 represents the width-height coordinates on frame *n* for keypoint *k* in camera *v*. To reiterate, each row contains the labeled coordinates for a single body part seen from *V* views. The rows of this matrix contain examples from all available labeled keypoints, which are all used for learning the 3D projection. We exclude rows in which a body part is missing from one or more views. The number of examples used to estimate PCA is, as desired, always much larger than the label dimension (*NK >>* 2*V*). We perform PCA on **Y**_*MV*_ and keep the first three PCs, which we denote as **P** = (**P**_1_ **P**_2_ **P**_3_)*∈* ℝ^2*V ×*3^ and the data mean ***µ*** *∈* ℝ^2*V*^ . The three PCs form three orthogonal axes in 2*V* dimensions, and projecting the 2*V* -dimensional labels on them will provide width-height-depth-like coordinates. These 3D coordinates are related to the “real-world” 3D coordinates (relative to some arbitrary “origin” point) by an affine transformation (they need to be rotated, stretched and translated), but critically, we do not need these “real-world” coordinates to apply the multi-view constraints during network training, as described below.

### During training: Penalizing the unlabeled data for PCA reconstruction errors

Let 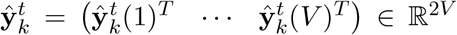 be the network’s prediction for the *k*-th body part on the *t*-th unlabeled video frame, on all *V* views (as before, this requires specifying the indices of corresponding keypoints across views). The prediction’s multi-view PCA reconstruction is given by projecting it down to 3 dimensions and then back up to 2*V* dimensions:

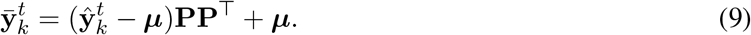

When the prediction 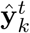 is consistent across views, i.e., on the 3D hyperplane specified by **P**, we will get 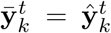, a perfect reconstruction. The loss is defined as the average pixel distance between each 2D predicted keypoint 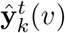 and its multi-view PCA reconstruction 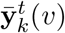:

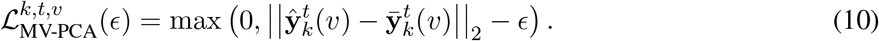

The loss encourages the predictions to stay within the fixed 3D hyperplane estimated by PCA, and thus be consistent across views. In training, we minimize its average across views, body parts, and frames

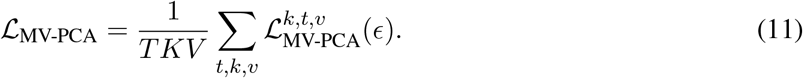

We choose *ϵ* by computing the PCA reconstruction errors (in pixels) for each of the labeled keypoints, and taking the maximum. This represents the maximal multi-view inconsistency observed in the labeled data.

We note that the multi-view PCA loss does not require any modifications to network architecture. Each view is processed independently by the network. As mentioned above, all that is required is specification of which keypoints from which views correspond to the same body part. The mirrored datasets considered in this paper are handled similarly: the single frame containing all available views is processed by the network, and different keypoints are linked to the same body part via an entry in the model configuration file.

#### 4.4.3 Pose PCA loss

There are certain things that bodies cannot do. We might track 2*K* pose coordinates but it does not mean that they can all move independently and freely. Indeed, there is a long history of using low-dimensional models to describe animal movement [18–20]. Here, we extend the PCA approach to full pose vectors, and constrain the 2*K*-dimensional poses to lie on a low-dimensional hyperplane of plausible poses, which we estimate from the labels.

### Before training: fitting Pose PCA on the labels

This approach is identical to multi-view PCA, with the following exceptions. First, our observations are full pose vectors and not single keypoints seen from multiple views. The design matrix of labels is therefore shorter and wider **Y**_P-PCA_ *∈* ℝ^*N×*2*K*^; it has as many rows as labeled frames, and each row contains the entire pose vector. Rows (poses) with missing bodyparts are discarded from this matrix. The number of examples available for PCA estimation is now simply the number of non-discarded labeled frames, *N*_train_, which is not allowed to be smaller than the number of pose coordinates, i.e., *N*_train_ *≥* 2*K*. A second exception is that instead of keeping three PCs, we keep as many PCs needed to explain 99% of the pose variance, denoted as *R <<* 2*K*. We collect the kept PCs as columns of a (2*K × R*) matrix **P** = (**P**_1_ *· · ·* **P**_*R*_) . Each of the PCs represents an axis of plausible whole-body movement, akin to previous approaches [19, 21]. Figure 2D shows that the number of kept PCs is usually less than half of the observation dimensions. We now keep **P** and ***µ*** *∈* ℝ^2*K*^ to be used in training. For multi-view setups, it is possible to form an even wider (*N ×* 2*KV*) design matrix, appending all *V* views, to jointly enforce the multi-view PCA loss. We have done so in the mirror-mouse and mirror-fish datasets.

### During training: penalizing for implausible poses

As in Eq. 9, we project the full predicted poses down to the low-dimensional hyperplane, then back up to 2*K* dimensions, to form their Pose PCA reconstructions. Then, for each 2D keypoint on each unlabeled video frame, we define the loss as the pixel error between the raw prediction 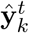 and its reconstruction 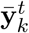:

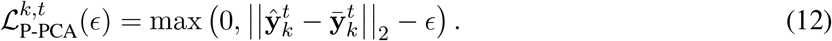

This loss tells us how many pixels are needed to move the predicted keypoint onto the hyperplane of plausible poses. During training, we minimize the average loss across keypoints and frames,

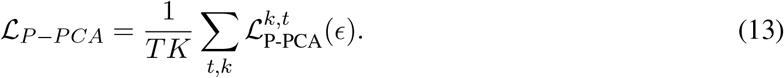

Here too, *ϵ* is chosen by reconstructing the labeled pose vectors, computing the pixel error between each 2D labeled keypoint and its PCA reconstruction, and taking the maximum value.

### 4.5 Training

Batch sizes are determined based on image size and GPU memory constraints (see Supplementary Table 3 for the batch sizes of the experiments reported in this paper). In general, denote a labeled batch size of *B* frames, a context window of 2*J* + 1 frames, and a short unlabeled clip of *T* frames (typically tens to hundreds) randomly drawn from a much longer video. The batch sizes will be *B* for a supervised model, *B* + *T* for a semi-supervised model, (2*J* + 1)*B* for a TCN model, and (2*J* + 1)*B* + *T* for a semi-supervised TCN model. In our TCN experiments we use *J* = 2. To efficiently use unlabeled clips for TCN models, we push the full clip through the backbone once, then discard predictions from the first and last *J* frames, which do not have sufficient context. To make our experiments controlled and reproducible across GPU types, we explicitly chose small labeled batch sizes, such that each of our model variants trains with an equal number of labeled frames per batch (the semi-supervised and TCN models see many more unlabeled frames per batch, which can become memory-prohibitive).

We use an Adam optimizer [22] with an initial learning rate of 0.001, halving it at epochs 150, 200, and 250. In the experiments reported here, the ResNet50 backbone was kept frozen for the first 20 epochs. We trained our models for a minimum number of 300 training epochs and a maximum number of 750 epochs. During training we split the InD data into training (80%), validation (10%), and test (10%) sets. We performed early stopping by checking the heatmap loss on validation data every five epochs and exiting training if it does not improve for three consecutive checks.

During training we apply standard image augmentations to labeled frames including geometric transforms (e.g. rotations and crops), color space manipulations (e.g. histogram equalization) and kernel filters (e.g. motion blur), following [10]. A different random combination of augmentations is used for each frame in a batch. For the TCN architecture, the same augmentation combination is used for a labeled frame and its associated context frames. For the semi-supervised models, we apply augmentations to unlabeled video frames using DALI. A single random combination of augmentations is used for all video frames in a batch. Because the PCA losses are sensitive to geometric transforms, once the width-height coordinates have been inferred using the soft argmax described above we apply the inverse geometric transform before computing unsupervised losses.

While our package includes well-tested default hyperparameters for the unsupervised losses described in this paper, users implementing a new “bespoke” loss are advised to perform hyperparameter searches for this loss’s weight, which of course multiplies the amount of compute by the number of tested weights. However, hyperparameter searches can be run in parallel, and our Hydra scripts enable users to launch and log these jobs without additional custom scripts.

### 4.6 Diagnostics and model selection

#### 4.6.1 Constraint violations as diagnostic metrics

After training, we evaluate the network on the the labeled frames and on unlabeled videos. We then compute our individual loss terms (defined in Eq. 6, 10, 12) for each predicted keypoint, on each frame, and on each view for a multi-view setup, and use them as diagnostic metrics. For labeled frames, we compute the Euclidean pixel error. All metrics are measured as pixel distances on the full-sized image.

#### 4.6.2 Model selection based on pixel errors and constraint violations

Our loss factory requires users to select among different applicable losses, and for each loss, determine its weight (note that we offer robust default values in our package). We start by fitting a baseline model to the data (typically with three random seeds). Then, for each of the applicable losses, we search over 4 *−* 8 possible weights (between values of 3.0 and 7.0). We then compare the diagnostic metrics specified above on a held-out validation set (ignoring errors below a tolerance threshold). We pick the weight that exhibits the minimal loss across the majority of our diagnostics. Supplementary Table 4 displays the optimal weight chosen for each loss in each dataset using non-TCN models. We used the same weights for the TCN networks.

### 4.7 Sample efficiency experiments

The sample efficiency experiments in Fig. 1C demonstrate model performance on InD and OOD data as a function of training frames. For a given network trained with *N* frames, we actually need to select *N* ^*∗*^ = `ceiling`(1.25*N*) frames to account for additional validation frames used for early stopping, as well as InD test frames (the train/val/test split was 80%/10%/10%). To mimic a realistic labeling scenario, we randomly selected a video from all the InD data. If the number of frames in this first video (call this *M*_1_) was greater than or equal to *N* ^*∗*^, then we stopped here. If *M*_1_<*N* ^*∗*^ we continued to randomly select a video and add all labeled frames from that video to the labeled data pool. Once 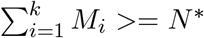, we randomly selected 10% of the frames in the pool for validation, 10% for testing, and of the remaining 80% we chose exactly *N* frames for training. Training was performed with supervised Lightning Pose models as described above. After training we computed InD pixel error on the 10% of test frames, and OOD pixel error on held-out videos that were never considered for the labeled data pool. We repeated this procedure 10 times for each value of *N* .

### 4.8 Ablation study showing the effects of individual losses

The goal of this analysis is to quantify the relative contribution of the individual unsupervised losses in the mirror-mouse, mirror-fish, and CRIM13 datasets. We focus on the scarce label regime (75 train frames), where the semi-supervised improvements are most pronounced. We train semi-supervised models with either temporal, multi-view PCA, or Pose PCA losses, and compare these to a supervised baseline and a semi-supervised model that combines all loss types. For each condition, we train three networks with different random seeds controlling the data presentation order. To simplify this analysis, we analyze pixel-error averages. The results indicate that across datasets, most pixel error savings were driven by the multiview and Pose PCA losses (Extended Data Fig. 3). A combination of all losses always performs the best.

### 4.9 DeepLabCut Training

For DeepLabCut experiments (version 2.2.3), we use their default parameters: an ImageNet-pretrained backbone, training for 50k “iterations” (batches) independent of the labeled dataset size, using the Adam optimizer [22] with a learning rate schedule that starts from 1e-4 and is reduced to 5e-5 at iteration 7500 then to 1e-5 at iteration 12000. We select the training frames to exactly match those used for the Lightning Pose models in all analyses with the mirror-mouse, mirror-fish, and CRIM13 datasets. For the IBL datasets, we use the same number of training frames but do not try to match them exactly. For differences between the baseline and DeepLabCut models, see the Supplementary Information.

### 4.10 Ensembling

To perform ensembling, we need a collection of models that output a diverse set of predictions. This can be achieved through various means. For the EKS analyses in Extended Data Fig. 6 we chose to study a single split of the data, and achieved diversity by randomly initializing the head of each model, as well the order in which the data was sent to the model during training. Despite these seemingly minor differences, the ensemble of models produced a variety of outputs (Extended Data Fig. 6B,D,F). For the other figures and videos related to ensembling (Figs. 5, 6, Extended Data Fig. 7; Supplementary Video 8-14; Supplementary Figs. 2-4) we achieved diversity by training each model with a different subset of training data (in line with the analyses performed in, e.g., Fig. 4).

### 4.11 Post-processor comparison

For the post-processor comparisons in Fig. 5 we used the following baselines:

### Median filter

We used the `medfilt` function from the SciPy package [23] using the default settings from the DeepLabCut package (`kernel_size=5`).

### ARIMA

We used a Seasonal Autoregressive Integrated Moving-Average with eXogenous regressors (SARIMAX) model using the default settings from the DeepLabCut package (`pcutoff=0.001`, `alpha=0.01`, `ARdegree=3`, `MAdegree=1`).

### Ensemble mean/median

We computed the mean/median over the ensemble members, independently for the x and y coordinates. We did not threshold by confidence.

### 4.12 Ensemble Kalman Smoother

The Ensemble Kalman Smoother (EKS) begins with the output of the ensemble of pose-estimation networks, an *m ×* 2*KV × T* tensor, for *m* ensemble members (here, *m ≈* 5), *K* keypoints, *V* views, and *T* video frames. EKS performs probabilistic inference to denoise the ensemble predictions to obtain more accurate and robust pose estimates. To be more specific, we compute the mean and variance for each keypoint across the ensemble to obtain the 2*KV × T* ensemble mean *M* and variance *C* matrices.

We first define the general state-space model, then discuss its useful special cases in the following sections. We specify a latent *state* variable *q*_*t*_, a linear Gaussian Markov *dynamics* model for this state variable of the form

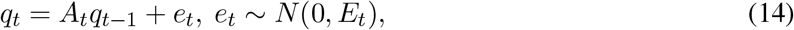

and a linear Gaussian *observation* model describing the relationship between the latent state variable *q*_*t*_ and the observed data *O*_*t*_,

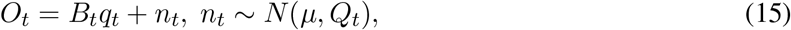

for some appropriate (potentially time-varying) system parameters *A*_*t*_, *B*_*t*_, *E*_*t*_, *Q*_*t*_, *µ*.

#### 4.12.1 Single-keypoint, single-camera case

This is the simplest case to consider: imagine that we want to denoise each keypoint individually, and we only have observations from a single camera. Here the latent state *q*_*t*_ is the true two-dimensional position of the keypoint on the camera. Now our model is

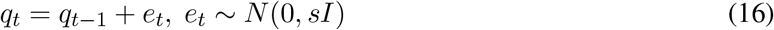

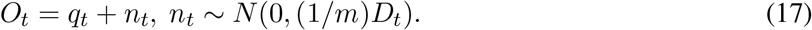

Comparing these equations to the general dynamics and observations equations above, we see that *A*_*t*_ = *B*_*t*_ = *I* here.

In the observation equation, *O*_*t*_ is the 2 *×* 1 keypoint vector, and *D*_*t*_ is a 2 *×* 2 diagonal matrix specifying the ensemble confidence about each observation. We use the *t*-th column of the ensemble mean *M* to fill in the observation *O*_*t*_, and the covariance from the *t*-th frame of the ensemble covariance *C* to fill in the observation variance *D*_*t*_ (note that larger values of *D*_*t*_ correspond to lower confidence in the corresponding observation *O*_*t*_). The factor of (1*/m*) in the observation variance follows from the fact that *O*_*t*_ is defined as a sample mean over *m* ensemble members.

Finally, *s* is an adjustable smoothing parameter: larger *s* leads to less smoothing. This smoothness parameter could be selected by maximum likelihood (e.g., using the standard expectation-maximization algorithm for the Kalman model) but can be set manually for simplicity.

Now, given the specified dynamics and observation model, we can run the standard Kalman forward-backward smoother to obtain the posterior mean state *Q* given the observations *O* (i.e., all the states *q*_*t*_ given all the observations *O*_*t*_). The smoother will “upweight” high-confidence observations *O*_*t*_ (i.e., small *D*_*t*_), and “downweight” low-confidence observations (large *D*_*t*_), e.g., from occlusion frames.

Note that this Kalman approach is the Bayesian optimal estimator under the assumption that the model in Eqs. 16-17 is accurate. In reality, this model holds only approximately: in general, neither the observation noise nor the state dynamics are exactly Gaussian. Therefore the Ensemble Kalman Smoother should be interpreted as an approximation to the optimal Bayesian estimator here. Generalizations (to handle multimodal observation densities, or switching or stochastic volatility dynamics models) are left for future work.

#### 4.12.2 Single-keypoint, multi-camera, synchronized cameras case

Given multiple cameras, we can estimate the true three-dimensional position of each keypoint. So letting the state vector *q*_*t*_ be the three-dimensional vector *q*_*t*_ = (*x*_*t*_, *y*_*t*_, *z*_*t*_), we have the model

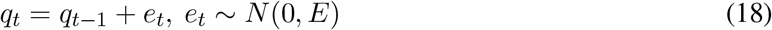

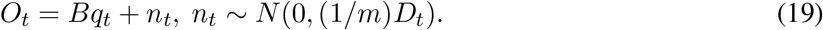

*B* is 2*V ×* 3 where *V* is the number of camera views; this maps the three-dimensional state vector *q*_*t*_ onto the *V* camera coordinates (assuming linear observations here; this can be generalized but was not necessary for the data analyzed here). *O*_*t*_ is 2*V ×* 1 and *D*_*t*_ is block-diagonal with 2 *×* 2 blocks. As above, observations *O*_*t*_ with high *D*_*t*_ (low confidence) will be downweighted by the resulting Ensemble Kalman Smoother: in practice, this means that cameras with an unobstructed view on a given frame (small *D*_*t*_) can help to correct frames that are occluded in other camera views (resulting in larger ensemble variance *D*_*t*_). We remark that in poorly trained models, the opposite can also (on rarer occasions) be true: the ensemble in one camera view can make “confident mistakes” on some frames, in which all ensemble members output the same wrong estimate (with corresponding small *D*_*t*_, i.e., high ensemble confidence) and induce errors in the other camera views after running the Ensemble Kalman Smoother. These errors can be detected as deviations between the Kalman smoother output and the original ensemble outputs; the training label set can then be augmented to correct these confident mistakes, followed by network ensemble retraining.

We initialize our estimates by restricting to confident frames and computing PCA to estimate *B*; then we take temporal differences of the resulting PCA projections and compute their covariance to initialize *E*.

Finally, note that this simple Kalman model does not output the true 3d location here, because the model is non-identifiable; instead we learn *q*_*t*_ up to a fixed invertible affine transformation.

#### 4.12.3 Pupil EKS

For the IBL-pupil dataset, we track *K* = 4 keypoints arranged in a diamond shape around the perimeter of the pupil. Therefore, at each frame we have 2*K* = 8 observations which are are constrained to lie in a three-dimensional subspace defined by the pupil center (denoted as (*x*_*t*_, *y*_*t*_)) and diameter *d*_*t*_. Given the state variable *q*_*t*_ = (*d*_*t*_, *x*_*t*_, *y*_*t*_), we can (linearly) predict the location of each of the 4 diamond corners.

In addition, we have strong prior information about the dynamics of the state variable: we know that the diameter *d*_*t*_ is a smooth function of time *t*, while the pupil center (*x*_*t*_, *y*_*t*_) can change more abruptly, due to saccades and rapid face movements that move the eye as well.

Together, these assumptions lead to the model

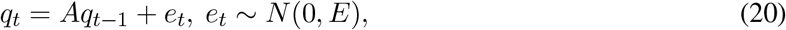

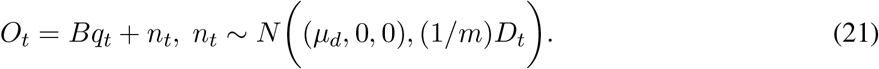

In the observation equation above, *µ*_*d*_ denotes the mean diameter, *O*_*t*_ is the 8 *×* 1 keypoint vector, *B* is a fixed 8 *×* 3 matrix that translates the state variable *q*_*t*_ into the keypoints, and *D*_*t*_ is a diagonal matrix whose diagonal entries include the ensemble confidence about each observation.

In the dynamics model above, *A* and *E* are both diagonal. This means that we model the priors for *d*_*t*_, *x*_*t*_, and *y*_*t*_ using independent autoregressive (AR(1)) processes. (The posteriors for these variables will not be independent, due to the non-separable structure of the observation model in Eq. 21.) We want to choose the diagonal values *diag*(*A*) and *diag*(*E*) so that these processes have the desired variance and time constant. The variance in a stationary AR(1) model with noise variance *e* and autoregressive parameter *a* is *e/*(1 *− a*^2^). We can crudely estimate the marginal mean and variance of *x*_*t*_, *y*_*t*_, and *d*_*t*_ from the ensembled mean *M*, and match the AR(1) marginal mean and variance accordingly. This leaves us with just two autoregressive parameters to choose: *A*(1, 1) and *A*(2, 2) (with *A*(3, 3) set equal to *A*(2, 2)). The time constant corresponding to *A*(1, 1) should be meaningfully larger than the time constant corresponding to *A*(2, 2), since as noted above the diameter *d*_*t*_ varies much more smoothly than the center (*x*_*t*_, *y*_*t*_).

#### 4.12.4 Single-keypoint, multi-camera, asynchronous cameras case

In some datasets (e.g. the IBL-paw dataset) frames from different cameras may be acquired asynchronously, perhaps with different frame rates. The Kalman model can be easily adapted to handle this case. Define the sampling times and camera ID for the *i*-th frame as: *{t*_*i*_, *v*_*i*_*}*, where *t*_*i*_ denotes the time the frame was acquired, and *v*_*i*_ denotes the camera that took the *i*-th frame. Again the state vector *q*_*t*_ is the true threedimensional location of the keypoint, *q*_*t*_ = (*x*_*t*_, *y*_*t*_, *z*_*t*_). We have the model

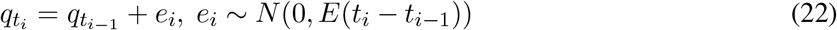

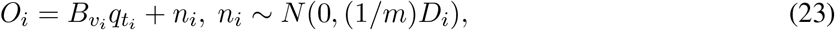

where now 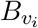 is 2 *×* 3; this tells us how the latent 3d coordinates are mapped into the *v*_*i*_’th camera. *O*_*i*_ is a 2 *×* 1 vector, and *D*_*i*_ is a 2 *×* 2 matrix. Here the Kalman Smoother is run only at frame acquisition times *{t*_*i*_*}*, but if desired we can perform predictions / interpolation at any desired time *t*.

#### 4.12.5 Pose PCA case

Let *q*_*t*_ represent the “compressed pose,” the *R ×* 1 vector obtained by projecting the true pose into the *R*-dimensional Pose PCA subspace. Here we have the model

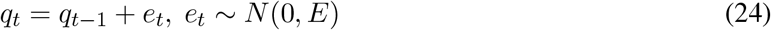

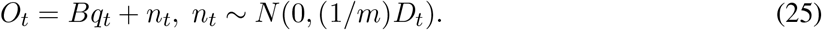

*B* is 2*K × R*; this maps the *R*-dimensional state vector *q*_*t*_ onto the 2*K* camera coordinates. *O*_*t*_ is 2*K ×* 1 and *D*_*t*_ is block-diagonal with 2 *×* 2 blocks. As in the synchronous multi-camera setting, we initialize our estimates by restricting to confident frames and computing PCA to estimate *B*; then we take temporal differences of the resulting PCA projections and compute their covariance to initialize *E*.

The output of this smoother is useful for diagnostic purposes, but we do not recommend using this model to generate the final tracking output, since rare (but real) poses may lie outside the Pose PCA subspace, while the output of this smoother is restricted to lie within this subspace (the span of *B*) by construction.

### 4.13 Canonical correlations analysis (CCA)

In Supplementary Fig. 2 and Supplementary Fig. 4 we use canonical correlations analysis to compute the directions of motion that should match in the left and right cameras and top and bottom cameras, respectively. (These canonical correlations directions are orthogonal to the epipolar lines familiar from multiple view geometry [15].) In this subsection we provide details of this computation.

Let 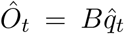 be the output of the multi-camera Ensemble Kalman Smoother at time step *t*, projected back onto the camera planes. We can further decompose *Ô*_*t*_ as 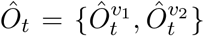, where 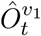 is the two-dimensional prediction for the first camera and 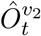 is the two-dimensional prediction for the second camera. Now, we compute 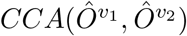 to find the one-dimensional linear projection of the outputs for each camera that maximizes their correlation. Since *Ô*_*t*_ is generated from a lower-dimensional set of latents *q*_*t*_, the projection of 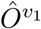 and 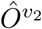 onto the first canonical component will be perfectly correlated. We can then project the original model predictions for each camera onto the first canonical component for each camera.

Any frames where the two camera-views do not have the same projected value will most likely be outliers. This can be seen in Supplementary Figs. 2 and 4, where outlier frames due to paw switching and paw occlusions cause the model predictions for the two camera views to have different CCA projections.

### 4.14 Neural decoding

We performed neural decoding using cross-validated linear regression with L2 regularization (the Ridge module in scikit-learn [24]), following [25]. The decoding targets – pupil diameter or paw speed – are binned into non-overlapping 20 ms bins. For each successful trial, we select an alignment event – reward delivery for pupil diameter and wheel movement onset for paw speed – and decode the target starting 200 ms before and ending at 1000 ms after the alignment event. We bin spike counts similarly using all recorded neurons in each session. The target value for a given bin (ending at time *t*) is decoded from spikes in a preceding (causal) window spanning *R* bins (ending at times *t*, …, *t*-*R*+1). Therefore, if decoding from *N* neurons, there are *RN* predictors of the target variable in a given bin. In practice we use *R* = 10.

To improve decoding performance, we smoothed the target variables. For pupil diameter, both the DeepLab-Cut (DLC) and Lightning Pose (LP) predictions of pupil diameter were smoothed using a Savitzky-Golay filter that linearly interpolates over low-confidence time points (confidence <0.9). The filter window is set to 31 frames (500 ms) for the left video (we did not decode pupil diameter from the lower-spatial-resolution right video). For more details of this method, see [3]. We did not apply additional smoothing to the output of the Ensemble Kalman Smoother (LP+EKS) model. For paw speed, small errors in the paw position will be magnified when taking the derivative. To compensate for this we lightly smoothed the paw position estimates using a Savitzky-Golay filter after linearly interpolating over low-confidence time points (confidence <0.9), and then computed paw speed. The right video filter window is set to 13 frames (87 ms) and the left is set to 7 frames (117 ms). This smoothing was applied to the outputs of all three models (DLC, LP, LP+EKS).

All decoding results use nested cross-validation. Each of five cross-validation folds is based on a training/validation set comprising 80% of the trials and a test set of the remaining 20% of trials. Trials are selected at random (in an “interleaved” manner). The training/validation set of a fold is itself split into five sub-folds using an interleaved 80%/20% partition. A model is trained on the 80% training set using various regularization coefficients (*{*10^*−*5^, 10^*−*4^, 10^*−*3^, 10^*−*2^, 10^*−*1^, 10^0^, 10^1^*}*, denoted as input parameter *α* by scikit-learn), and evaluated on the held-out validation set. This procedure is repeated for all five sub-folds. The coefficient which achieves the highest *R*^2^ value, averaged across all five validation sets, is selected as the “best” coefficient and used to train a new model across all trials in the 80% training/validation set. The model is then used to produce predictions for each trial in the 20% test set. This train/validate/test procedure is repeated five times, each time holding out a different 20% of test trials such that, after the five repetitions, 100% of trials have a held-out decoding prediction. The final reported decoding score is the *R*^2^ computed across all held-out predictions. Code for performing this decoding analysis can be found at https://github.com/int-brain-lab/paper-brain-wide-map.

### 4.15 Lightning Pose software package

We built Lightning Pose with the following philosophy. To begin with, computer vision is a vast field, of which animal pose estimation is a small part. The thriving deep learning software ecosystem offers well-engineered and well-tested solutions for every stage of the pose estimation pipeline. We can therefore outsource code to these frameworks to a large degree, leaving us with a smaller code base to maintain.

We start with Lightning Pose’s core components, which are depicted in the innermost purple box in Extended Data Fig. 8A.

First, an algorithmic signature of Lightning Pose is training with two data streams, labeled images and unlabeled videos (as depicted in Fig. 2A), which have to be loaded and “augmented” in tandem. This requirement led us to develop a generic class of so-called “data modules” supporting flexible semi-supervised training.

Most computer vision systems are built to ingest images, not videos; raw videos are rarely used during training. The standard approach converts raw videos into formatted (“augmented”) images using CPUs. The CPU approach is inefficient and may cause the network to spend most of its time idly waiting for data instead of predicting or training (“data bottleneck”; [26]). Therefore, we built high-performance video readers using NVIDIA’s data loading library (DALI; https://github.com/NVIDIA/DALI; leftmost box inside innermost purple box in Extended Data Fig. 8A). DALI uses the native capabilities of Graphics Processing Units (GPUs) to both read (“decode”) and augment videos (resize, crop, scale, etc) to greatly accelerate video handling at training and prediction time.

Moreover, Lightning Pose decouples network architectures from datasets and training losses (center and right boxes, respectively, inside innermost purple box in Extended Data Fig. 8A). As part of our own experiments, we realized that users need flexibility to compose a set of supervised and unsupervised losses without making any code changes. We therefore built a “loss factory” that enables developers to experiment with existing losses easily and also quickly prototype new losses. Losses can be applied at any level of representation in the network, ranging from the time series of predicted keypoints, through heatmaps, to hidden network features. New losses require minimal extra code, are automatically logged during training, and can contain their own trainable parameters and even trainable sub-networks.

Having established how we handle data, design networks, and select losses, we still need a procedure for training networks. We offload this task to PyTorch Lightning ([27]; middle box in Extended Data Fig. 8A), which is an increasingly popular wrapper around the PyTorch deep learning framework [13]. This enables us to use the latest strategies for training models, logging the results, and distributing computation across multiple GPUs, without having to modify any of our core modules described above as new training techniques emerge.

In addition, we use Hydra [28] to configure, launch, and log network training jobs (Extended Data Fig. 8A, outermost purple box). This eliminates a substantial amount of “boilerplate” code while increasing the reproducibility of training, which often depends on choices of random number generator, batch sizes, etc.

Finally, we developed a suite of interactive training diagnostics and model comparison tools, facilitating hyperparameter sensitivity analyses (Extended Data Fig. 8A, right gray box). During training, we provide online access to TensorBoard (https://www.tensorflow.org/tensorboard) to monitor the individual losses. After training, we use a Streamlit (https://streamlit.io) user interface to visualize per-keypoint diagnostics for both labeled frames and unlabeled videos. We also use a FiftyOne user interface (https://voxel51.com) for viewing images and videos along with multiple models’ predictions, enabling users to filter body parts and models, and browse moments of interest in predicted videos.

### 4.16 A cloud-hosted application for pose estimation as a service

More and more laboratories have access to the accelerated computers needed for running deep learning pipelines. But unfortunately, installing, executing, and maintaining deep learning pipelines on them remains a hurdle even for experienced software developers.

We built a browser application that uses cloud computers and allows users with no coding expertise to estimate animal pose using any computer with access to internet. Our app (Extended Data Fig. 8B) supports the full life cycle of animal pose estimation, from data annotation via LabelStudio (https://labelstud.io) to model training to video prediction and diagnostic visualization (via the open-source ecosystem introduced above). When launched by a user, the app starts a number of cloud machines equipped with the necessary hardware and software, which will turn off when idle. Our app is built on Lightning.ai’s (https://lightning.ai) infrastructure for cloud-hosted deep learning applications, removing technical obstacles related to resource provisioning, secure remote access, and software dependency management.

To conclude, as argued in [29], the cloud-centric approach we take serves to democratize analysis tools, improving scalability, code maintenance requirements, and computation time and cost. Our app enables developers who have created new losses or network architectures within the Lightning Pose software package to easily make these advances available to the broader audience through the cloud-based app. This ability significantly accelerates the process of moving model development from the prototyping to production stage.

For up-to-date installation instructions and a walk-through of the app, we refer the reader to the app’s documentation website (https://pose-app.readthedocs.io).

## 5 Data availability

We have made all labeled data used in this manuscript publicly available.

mirror-mouse

https://figshare.com/articles/dataset/Lightning_Pose_dataset_mirror-mouse/24993315

mirror-fish

https://figshare.com/articles/dataset/Lightning_Pose_dataset_mirror-fish/24993363

CRIM13

https://figshare.com/articles/dataset/Lightning_Pose_dataset_CRIM13/24993384

IBL-paw

https://ibl-brain-wide-map-public.s3.amazonaws.com/aggregates/Tags/2023_Q1_Biderman_Whiteway_et_al/_ibl_videoTracking.trainingDataPaw.7e79e865-f2fc-4709-b203-77dbdac6461f.zip

IBL-pupil

https://ibl-brain-wide-map-public.s3.amazonaws.com/aggregates/Tags/2023_Q1_Biderman_Whiteway_et_al/_ibl_videoTracking.trainingDataPupil.27dcdbb6-3646-4a50-886d-03190db68af3.zip

All of the model predictions on labeled frames and unlabeled videos are available at https://figshare.com/articles/dataset/Lightning_Pose_results_Nature_Methods_2024/25412248. These results, along with the labeled data, can be used to reproduce the main figures of the paper.

To access the IBL data analyzed in Figs. 6 and Extended Data Fig. 7, see the documentation at https://int-brain-lab.github.io/ONE/FAQ.html#how-do-i-download-the-datasets-cache-for-a-specific-ibl-pa and use the tag 2023_Q1_Biderman_Whiteway_et_al. This will provide access to spike sorted neural activity, trial timing variables (stimulus onset, feedback delivery, etc.), the original IBL DeepLabCut traces, and the raw videos.

## 6 Code availability

The code for Lightning Pose is available at https://github.com/danbider/lightning-pose under the MIT license. The repository also contains a Google Colab tutorial notebook that trains a model, forms predictions on videos, and visualizes the results. From the command-line interface, running `pip install lightning-pose` will install the latest release of Lightning Pose via the Python Package Index (PyPI).

The code for the Ensemble Kalman Smoother is available at https://github.com/paninski-lab/eks under the MIT license. The repository contains the core EKS code as well as scripts demonstrating how to use the code on several example datasets.

The code for the cloud-hosted application is available at https://github.com/Lightning-Universe/Pose-app under the Apache-2.0 license. This code enables launching our app locally or on cloud resources by creating a Lightning.ai account.

Code for reproducing the figures in the main text is available at https://github.com/themattinthehatt/lightning-pose-2024-nat-methods under the MIT license. This repository also includes a script for downloading all required data from the proper repositories.

The hardware and software used for IBL video collection is described in [8]. The protocols used in the mirror-mouse and mirror-fish datasets (both have the same video acquisition pipeline) is described in [1].

We used the following packages in our data analysis: CUDA toolkit (12.1.0), cuDNN (8.5.0.96), deeplabcut (2.3.5 for runtime benchmarking, 2.2.3 for everything else), ffmpeg (3.4.11), fiftyone (0.23.4), h5py (3.9.0), hydra-core (1.3.2), ibllib (2.32.3), imgaug (0.4.0), kaleido (0.2.1), kornia (0.6.12), lightning (2.1.0), lightning-pose (1.0.0), matplotlib (3.7.5), moviepy (1.0.3), numpy (1.24.4), nvidia-dali-cuda120 (1.28.0), opencv-python (4.9.0.80), pandas (2.0.3), pillow (9.5.0), plotly (5.15.0), scikit-learn (1.3.0), scipy (1.10.1), seaborn (0.12.2), streamlit (1.31.1), tensorboard (2.13.0), torchvision (0.15.2)

## 7 Extended Data Figures

**Extended Data Figure 1:**
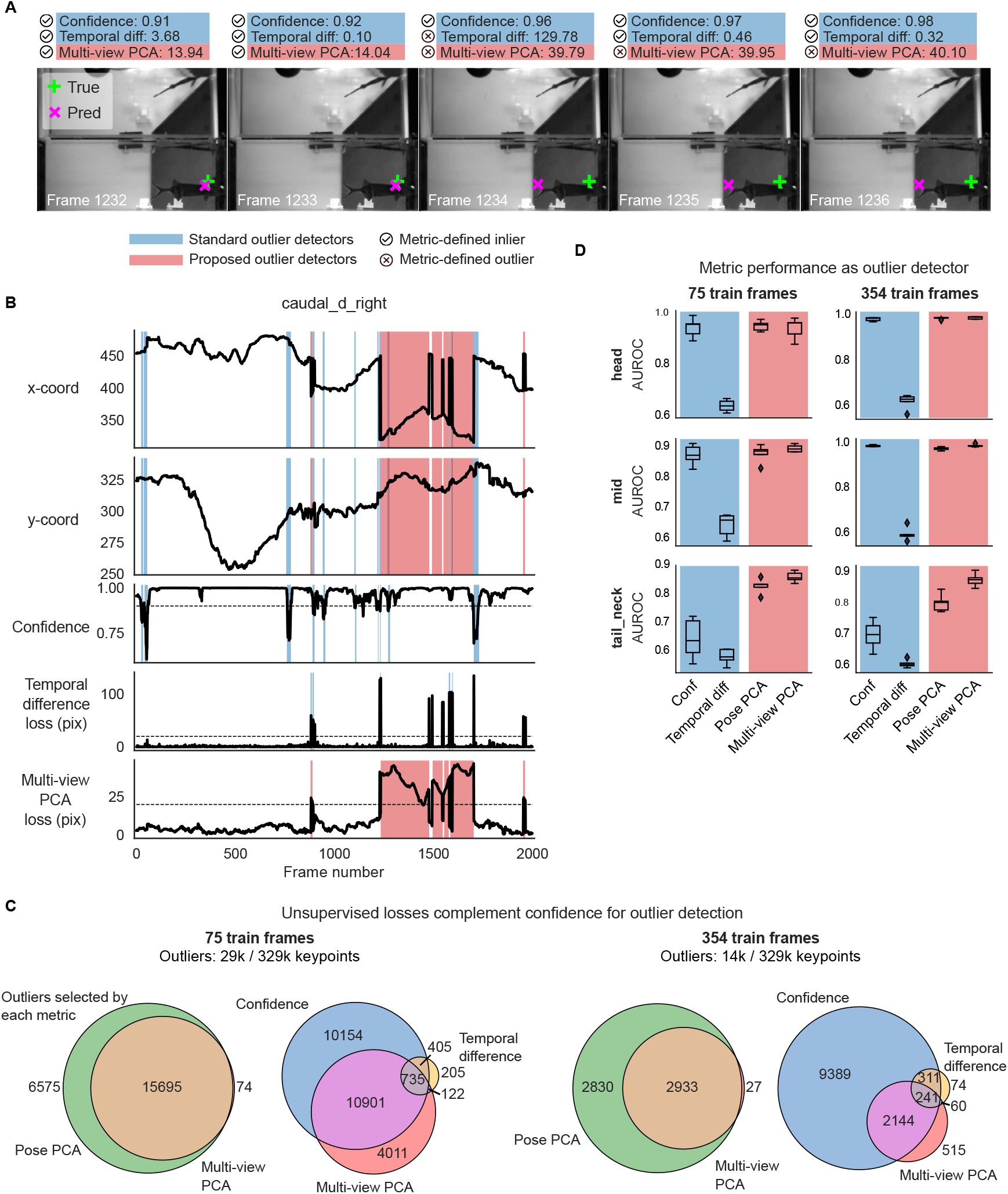
Unsupervised losses complement model confidence for outlier detection on mirror-fish dataset. Example traces, unsupervised metrics, and predictions from a DeepLabCut model (trained on 354 frames) on held-out videos. Conventions for panels A-D as in Fig. 3. **A**: Example frame sequence. **B**: Example traces from the same video. **C**: Total number of keypoints flagged as outliers by each metric, and their overlap. **D**: Area under the receiver operating characteristic curve for several body parts. We define a “true outlier” to be frames where the horizontal displacement between top and bottom predictions *or* the vertical displacement between top and right predictions exceeds 20 pixels. AUROC values are only shown for the three body parts that have corresponding keypoints across all three views included in the Pose PCA computation (many keypoints are excluded from the Pose PCA subspace due to many missing hand labels). AUROC values are computed across frames from 10 test videos; boxplot variability is over n=5 random subsets of training data. The same subset of keypoints is used for panel C. Boxes in panel D use 25th/50th/75th percentiles for min/center/max; whiskers extend to 1.5 * IQR.

**Extended Data Figure 2:**
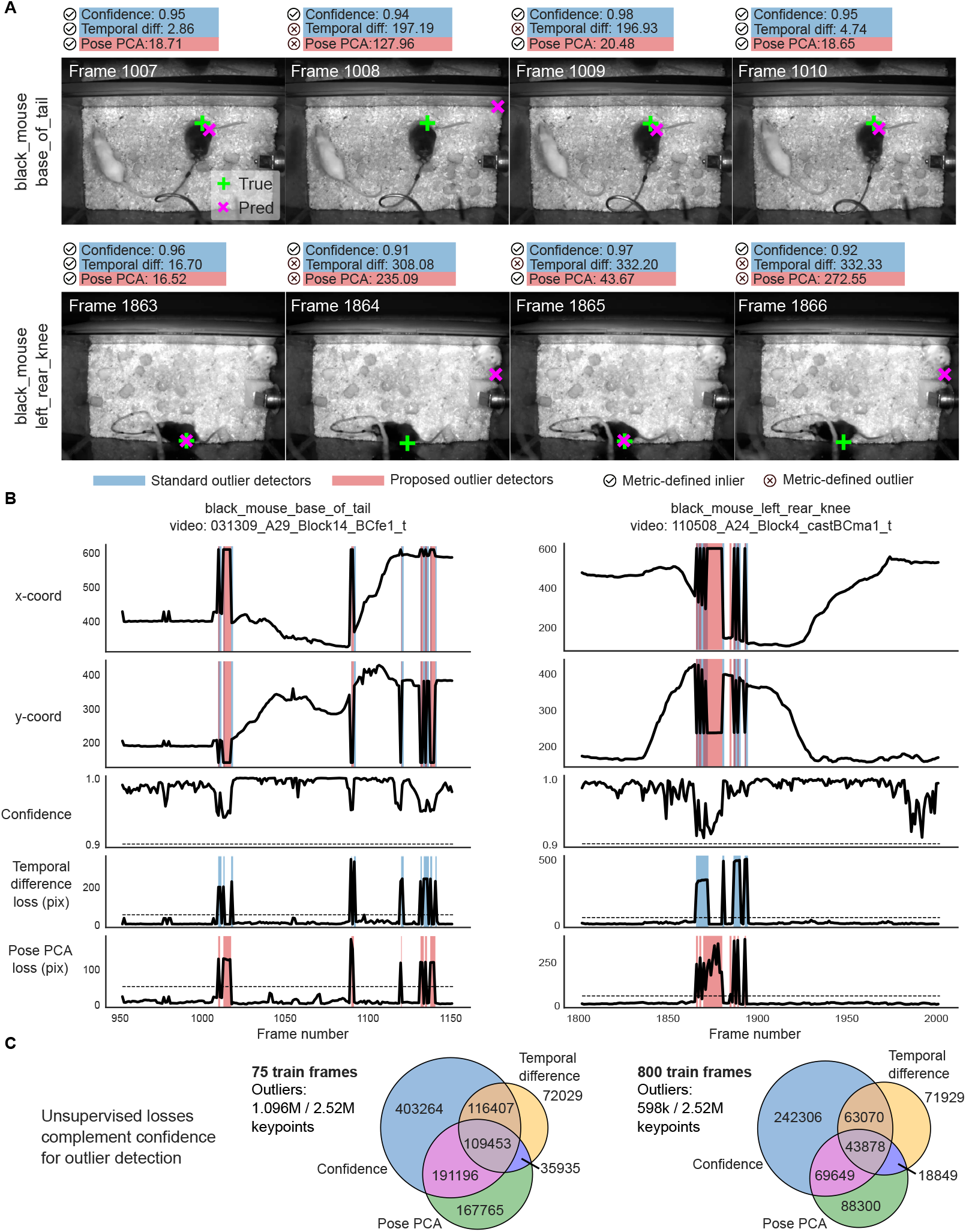
Unsupervised losses complement model confidence for outlier detection on CRIM13 dataset. Example traces, unsupervised metrics, and predictions from a DeepLabCut model (trained on 800 frames) on held-out videos. Conventions for panels A-C as in Fig. 3. **A**: Example frame sequence. In the first row, note that the second prediction jumps to the cage wall with high confidence, but is flagged as problematic by the Pose PCA loss. In the second row, the prediction again jumps back and forth between the mouse and the cage wall, and only the Pose PCA metric properly captures which predictions are outliers across all frames. **B**: Example traces from the same video. Because the size of CRIM13 frames are larger than those of the mirror-mouse and mirror-fish datasets, we use a threshold of 50 pixels instead of 20 to define outliers through the unsupervised losses. **C**: Total number of keypoints flagged as outliers by each metric, and their overlap. Outliers are collected from predictions across frames from 18 test videos and across predictions from five different networks trained on random subsets of labeled data.

**Extended Data Figure 3:**
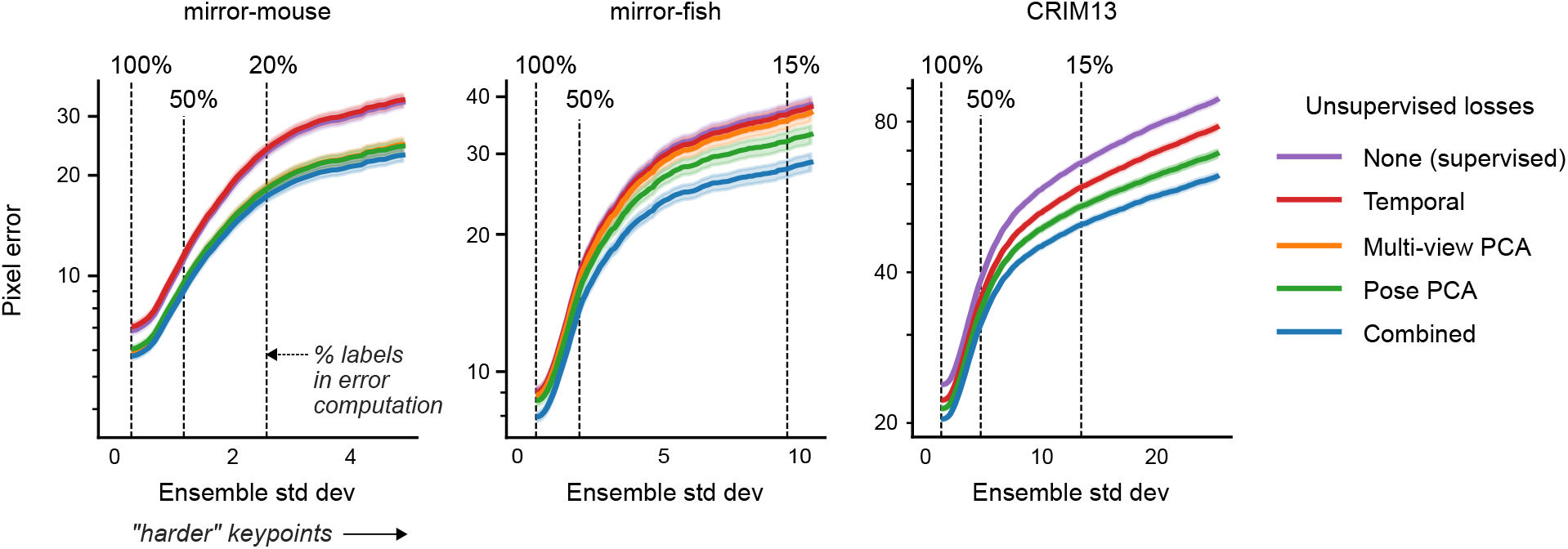
PCA-derived losses drive most improvements in semi-supervised models. For each model type we train three networks with different random seeds controlling the data presentation order. The models train on 75 labeled frames and unlabeled videos. We plot the mean pixel error and 95% CI across keypoints and OOD frames, as a function of ensemble standard deviation, as in Fig. 4. At the 100% vertical line, n=17150 keypoints for mirror-mouse, n=18180 for mirror-fish, and n=89180 for CRIM13.

**Extended Data Figure 4:**
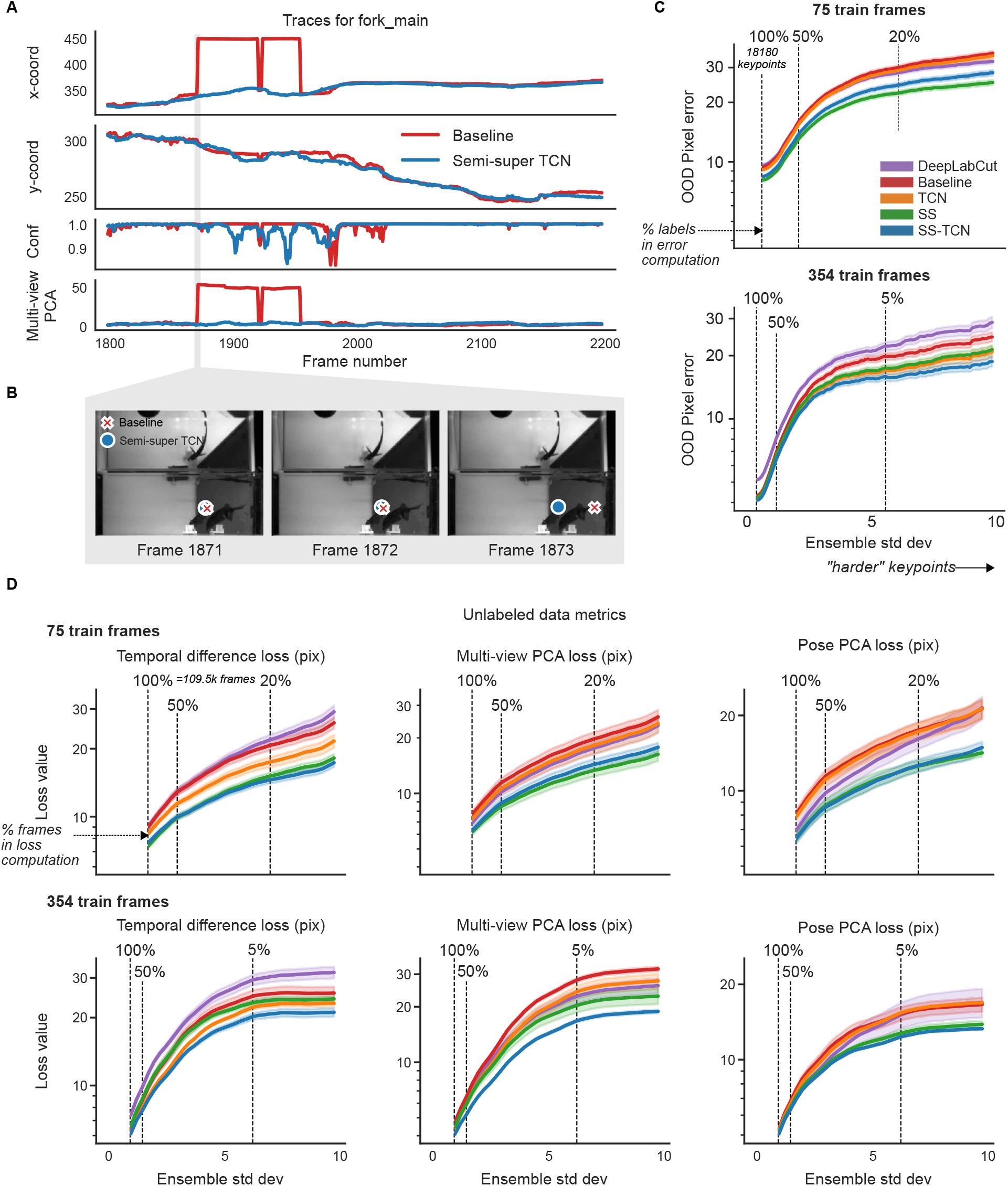
Unlabeled frames improve pose estimation in mirror-fish dataset. Conventions as in Fig. 4. Also see Supplementary Video 6.

**Extended Data Figure 5:**
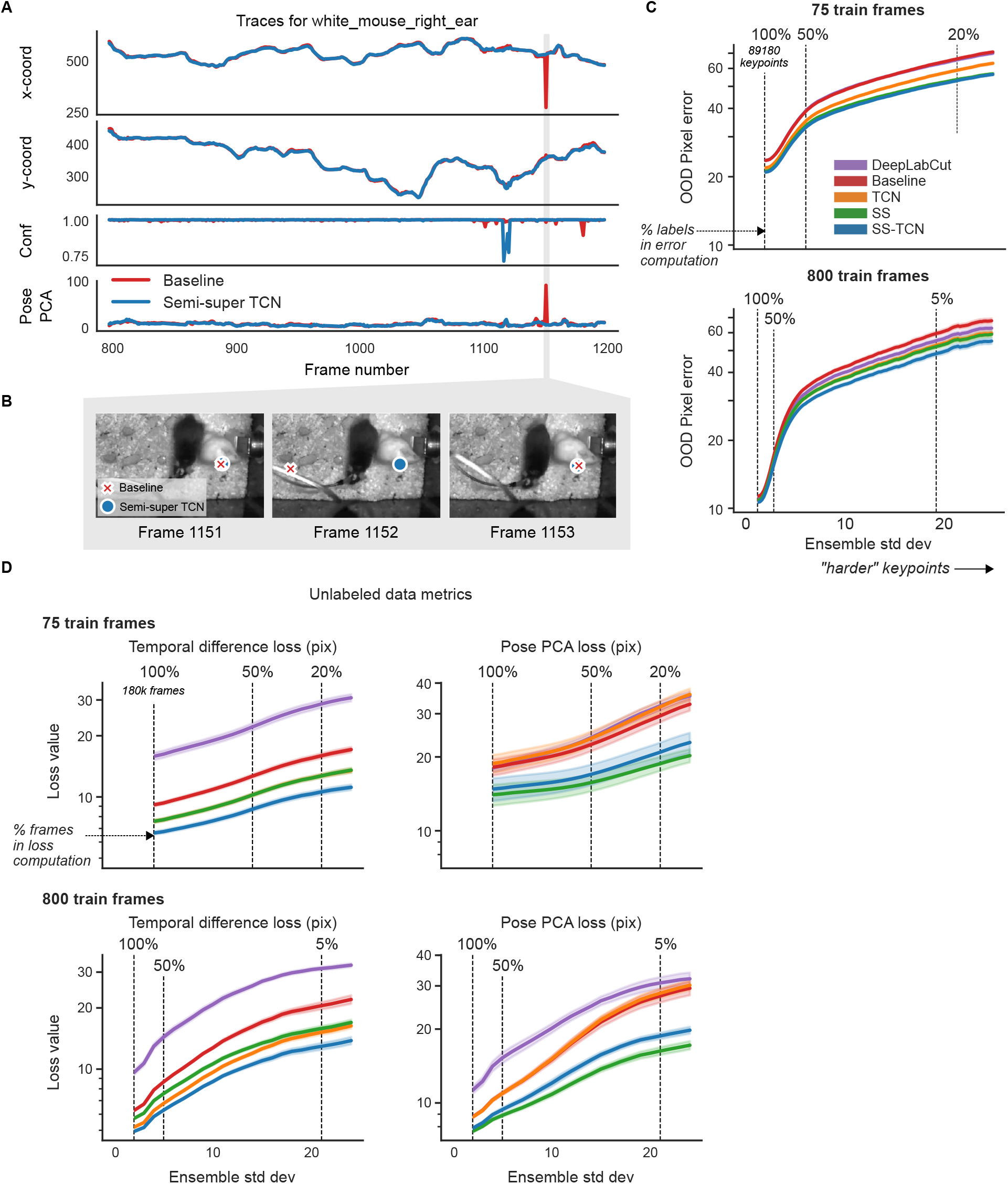
Unlabeled frames improve pose estimation in CRIM13 dataset. Models in panel A use 800 training frames. Remaining conventions as in Fig. 4. Also see Supplementary Video 7.

**Extended Data Figure 6:**
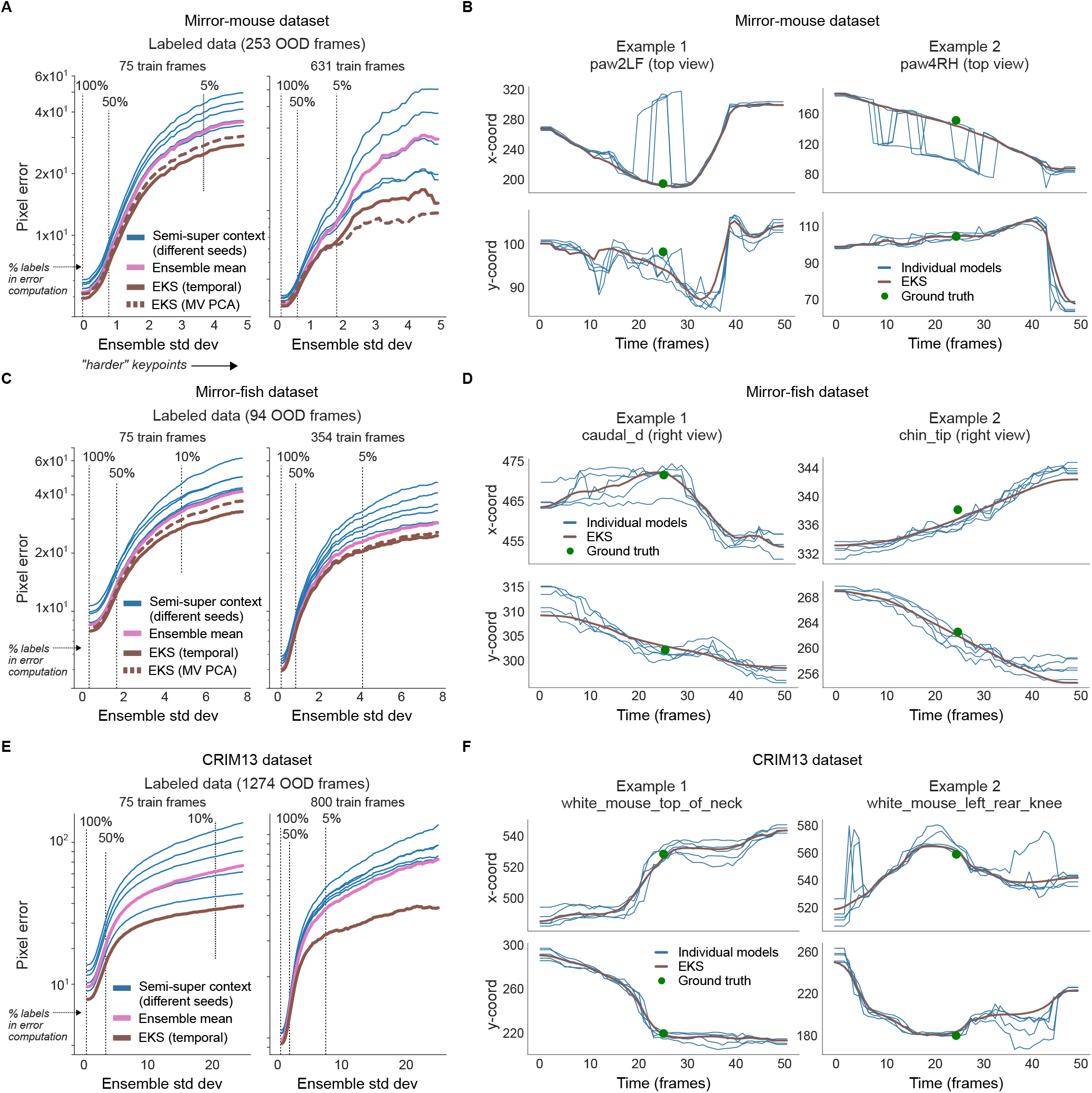
The Ensemble Kalman Smoother improves pose estimation across datasets. We trained an ensemble of five semi-supervised TCN models on the same training data. The networks differed in the order of data presentation and in the random weight initializations for their “head.” This figure complements Fig. 5 which uses an ensemble of DeepLabCut models as input to EKS, illustrating the flexibility of our method. **A**. Mean OOD pixel error over frames and keypoints as a function of ensemble standard deviation (as in Fig. 4). When the ensemble predictions disagree more strongly, EKS increasingly outperforms individual ensemble members and their mean. **B**. Time series of predictions (*x* and *y* coordinates on top and bottom, respectively) from the five individual semi-supervised TCN models (75 labeled training frames; blue lines) and EKS-temporal (brown lines). Ground truth labels are shown as green dots. **C**,**D**. Identical to A,B but for the mirror-fish dataset. **E**,**F**. Identical to A,B but for the CRIM13 dataset.

**Extended Data Figure 7:**
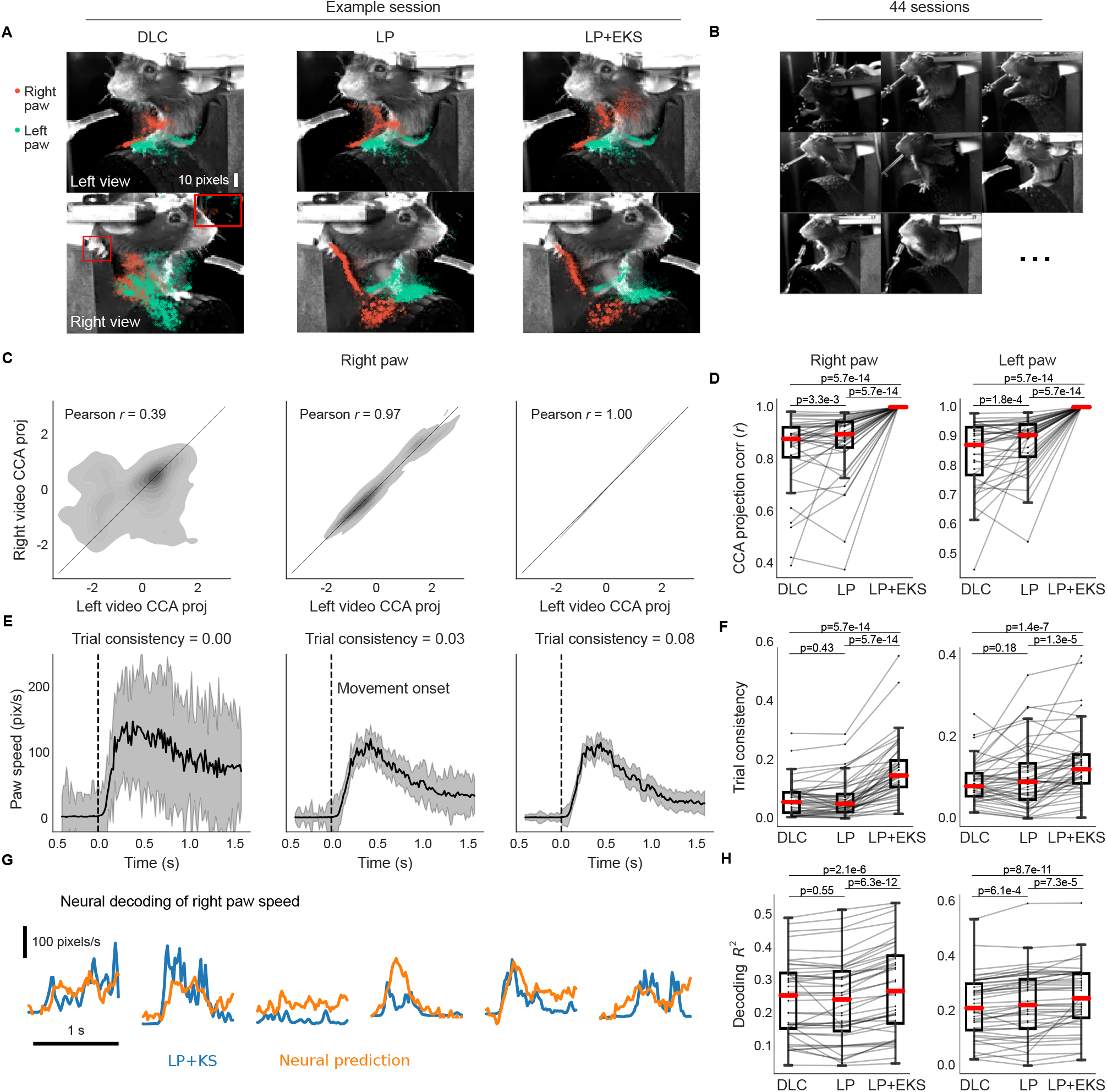
Lightning Pose models and ensemble smoothing improve pose estimation on IBL paw data. **A**. Sample frames from each camera view overlaid with a subset of paw markers estimated from DeepLabCut (*left*), Lightning Pose using a semi-supervised TCN model (*center*), and a 5-member ensemble using semi-supervised TCN models (*right*). **B**. Example left view frames from a subset of 44 IBL sessions, illustrating the diversity of imaging conditions in the dataset. **C**. As discussed in Supplementary Fig. 4, the right paw position in the right view should be highly correlated with right paw position in the left view; the 1D subspace of maximal correlation is found with canonical correlation analysis (CCA). Panel shows the empirical distribution of the right paw position projected onto this dimension from each view. Column arrangement as in A. The LP+EKS model imposes a low-dimensional model that enforces perfectly correlated projections, by construction. **D**. Correlation in the CCA subspace is computed across n=44 sessions for each model and paw. The LP+EKS model has a correlation of 1.0 by construction. **E**. Median right paw speed plotted across correct trials aligned to first movement onset of the wheel; error bars show 95% confidence interval across n=273 trials. The same trial consistency metric from Fig. 6 is computed. See Supplementary Video 14. **F**. Trial consistency computed across n=44 sessions. **G**. Example traces of Kalman smoothed right paw speed (blue) and predictions from neural activity (orange) for several trials using cross-validated, regularized linear regression (Methods). **H**. Neural decoding performance across n=44 sessions. Panels D, F, and H use a one-sided Wilcoxon signed-rank test; boxes use 25th/50th/75th percentiles for min/center/max; whiskers extend to 1.5 * IQR. See Supplementary Table 2 for further quantification of boxes.

**Extended Data Figure 8:**
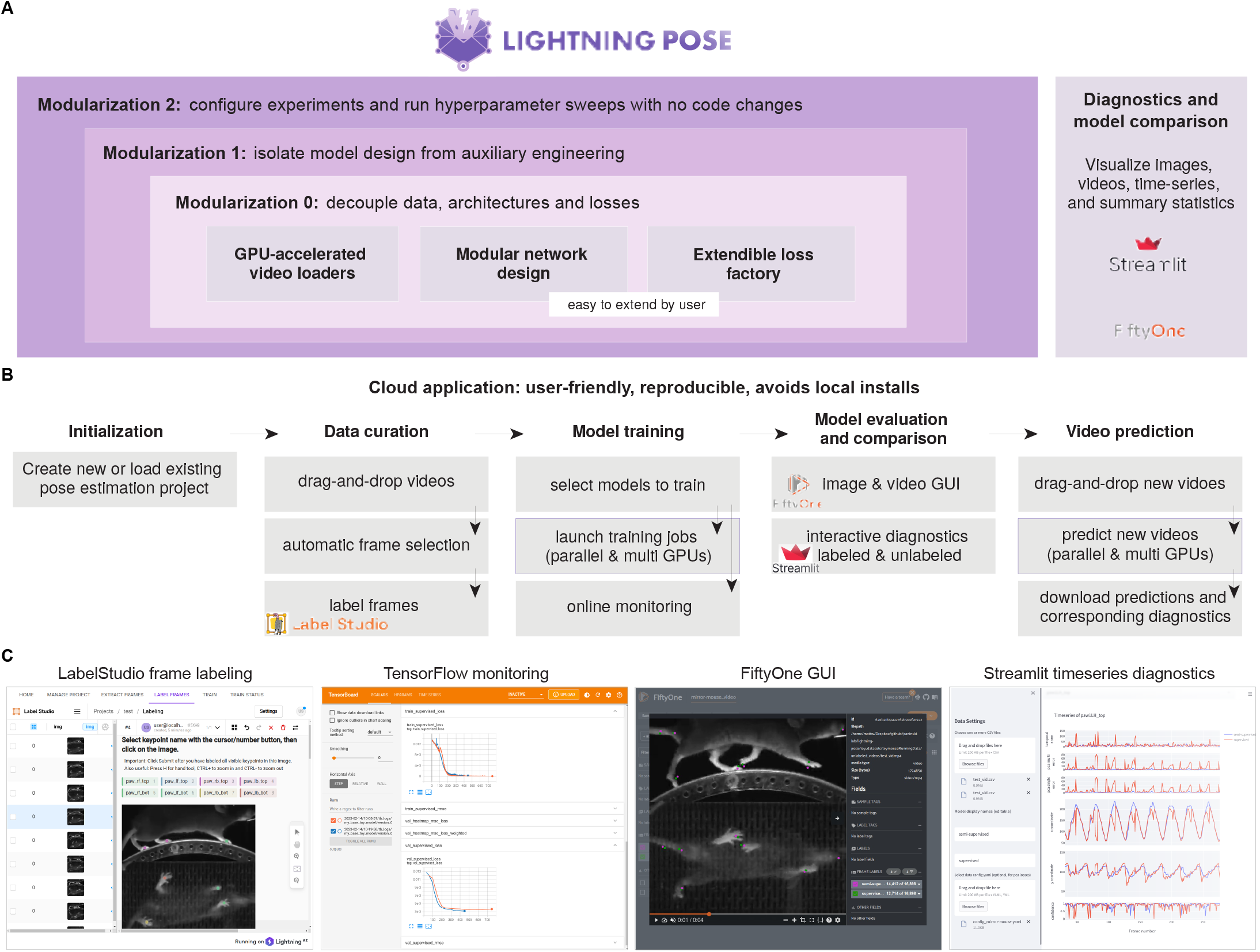
Lightning Pose enables easy model development, fast training, and is accessible via a cloud application. **A**. Our software package outsources many tasks to existing tools within the deep learning ecosystem, resulting in a lighter, modular package that is easy to maintain and extend. The innermost purple box indicates the core components: accelerated video reading (via NVIDIA DALI), modular network design, and our general-purpose loss factory. The middle purple box denotes the training and logging operations which we outsource to PyTorch Lightning, and the outermost purple box denotes our use of the Hydra job manager. The right box depicts a rich set of interactive diagnostic metrics which are served via Streamlit and FiftyOne GUIs. **B**. A diagram of our cloud application. The application’s critical components are dataset curation, parallel model training, interactive performance diagnostics, and parallel prediction of new videos. **C**. Screenshots from our cloud application. From left to right: LabelStudio GUI for frame labeling, TensorFlow monitoring of training performance overlaying two different networks, FiftyOne GUI for comparing these two networks’ predictions on a video, and a Streamlit application that shows these two networks’ time series of predictions, confidences, and spatiotemporal constraint violations.

## 8 Supplementary Video Captions

**Supplementary Video 1** DeepLabCut model predictions (631 labeled frames), mirror-mouse dataset, corresponding to traces in Fig. 1B. See Supplementary Fig. 5 for a detailed caption of Supplementary Videos 1, 5-7. [link]

**Supplementary Video 2** Selected frames with high Pose PCA errors, mirror-mouse dataset. DLC model trained with 631 labeled frames. See Supplementary Fig. 6 for a detailed caption of Supplementary Videos 2-4. [link]

**Supplementary Video 3** Selected frames with high Pose PCA errors, mirror-fish dataset. DLC model trained with 354 labeled frames. [link]

**Supplementary Video 4** Selected frames with high Pose PCA errors, CRIM13 dataset. DLC model trained with 800 labeled frames. [link]

**Supplementary Video 5** Baseline vs semi-supervised TCN model predictions (75 labeled frames), mirror-mouse dataset, corresponding to traces in Fig. 4A. [link]

**Supplementary Video 6** Baseline vs semi-supervised TCN model predictions (75 labeled frames), mirror-fish dataset, corresponding to traces in Extended Data Fig. 4A. [link]

**Supplementary Video 7** Baseline vs semi-supervised TCN model predictions (800 labeled frames), CRIM13 dataset, corresponding to traces in Extended Data Fig. 5A. [link]

**Supplementary Video 8** Ensemble Kalman Smoother predictions (multi-view PCA), mirror-mouse dataset, corresponding to the session in Supplementary Fig. 2. See Supplementary Fig. 7 for a detailed caption of Supplementary Videos 8-12. [link]

**Supplementary Video 9** Ensemble Kalman Smoother predictions (Pose PCA), IBL-pupil dataset, corresponding to the session in Supplementary Fig. 3. [link]

**Supplementary Video 10** Ensemble Kalman Smoother predictions (multi-view PCA), IBL-paw dataset, corresponding to the session in Supplementary Fig. 4. [link]

**Supplementary Video 11** Ensemble Kalman Smoother predictions (multi-view PCA), mirror-fish dataset. [link]

**Supplementary Video 12** Ensemble Kalman Smoother predictions (temporal), CRIM13 dataset. [link]

**Supplementary Video 13** Trial-by-trial markers and traces for IBL-pupil dataset, corresponding to the example session in Fig. 6. See Supplementary Fig. 8 for a detailed caption of Supplementary Videos 13-14. [link]

**Supplementary Video 14** Trial-by-trial markers and traces for IBL-paw dataset, corresponding to the example session in Extended Data Fig. 7. [link]

## Lightning Pose: Supplementary Information

### 1 Supplementary Methods

#### 1.1 Runtime benchmarking

We used the mirror-mouse dataset (two views, 17 total keypoints) and measured training and inference time. We compared each of our model variants – supervised baseline, TCN, semi-supervised (combining the three losses presented in the main text), semi-supervised TCN model – and DeepLabCut (Mathis *et al*. [1]; version 2.3.5). For all models, we used a ResNet-50 backbone. All experiments ran on an NVIDIA A10 GPU (using 16 CPU threads and 24 GB GPU memory, costing 1.624 USD per hour on-demand on Amazon Web Services as of September 2023).

##### 1.1.1 Training time

We measured the time it takes to load a batch of data from disk and complete a stochastic gradient descent step, an operation that forms the building block of neural network training. We varied two parameters: image resizing dimensions (128 *×* 128 and 256 *×* 256; images are read and resized “on the fly”), and labeled batch size *B* (16 and 32 images). We timed 100 training batches after discarding the first 15 “warmup” training batches. For DeepLabCut, we used the Tensorpack accelerated data loading package (henceforth DLC+tensorpack). As expected, the supervised model’s training time is on par with DLC+tensorpack (Supplementary Fig. 9). All other models train on a significant number of additional unlabeled frames and are therefore slower to train. The TCN (with *J* = 5 context frames) operates on a total of 5*B* frames and is 2x-5x slower; the semi-supervised model appends an additional unlabeled video clip of length 2*B* frames (for a total of 3*B* frames) and is 2x-4x slower; their combination operates on 7*B* frames and is 3x-5x slower. To put these estimates in perspective, a typical network will perform at minimum 300 passes over the entire labeled dataset (a.k.a. “epochs”). With 631 labeled InD images of the mirror-mouse dataset (each 256 *×* 256), and a batch size of 32, each epoch will include *I*631*/*32*1* = 20 batches. Throughout training, the semisupervised model will ingest additional 20 batches *×* 64 video frames *×* 300 epochs = 384*K* unlabeled video frames, and will train in approximately (0.44*s ×* 20 *×* 300)*/*60*s* = 44 minutes on a single NVIDIA A10 GPU (excluding logging operations) which will cost approximately 1.2 USD.

##### 1.1.2 Inference time

Next, we calculate the speed at which a trained network predicts a video (in frames-per-second, FPS). We compare three models: Lightning Pose without context (LP; either supervised or semi-supervised, since the training strategy does not affect inference time), LP with context (LP+TCN), and DLC. We calculate FPS for increasing image sizes (128x128, 256x256, 384x384, 512x512) and sequence lengths (16, 32, 64, 128, 256, 512). LP and LP+TCN use NVIDIA-DALI video loading which includes an additional resizing operation “on the fly,” which is not done by DeepLabCut.

Supplementary Fig. 10 summarizes the results. LP (orange) tends to have the highest throughput, with DLC (blue) 5% slower, and LP+TCN (green) 50% slower when averaged across all sequence lengths and frame sizes. LP speedups become more pronounced with longer sequence lengths and larger frame sizes, where GPU utilization increases. For smaller image sizes and sequence lengths, LP and LP+TCN are slower due to a paradoxical “data bottleneck”: since NVIDIA-DALI decodes and augments the videos entirely on the GPU, using very small batch sizes is inefficient as it “seeks” the entire video to find each batch. The seeking operation occupies the GPU instead of performing backward and forward passes through the network. In this unconventional regime, and with many CPUs available, standard CPU dataloaders like DeepLabCut’s are preferable. To summarize, when GPUs are properly utilized, LP’s inference is faster, as expected.

### 1.2 EKS’ Relationship to previous work

We are far from the first to notice that the output of pose tracking networks can contain “glitches,” and a number of strategies have been proposed for post-processing the network output to remove these glitches.

The simplest and perhaps most commonly applied strategy [2–4] is to detect “bad” keypoints and frames and remove them, followed by simple temporal interpolation to fill in the resulting gaps in the estimated pose traces. A number of criteria have been proposed to find bad frames, e.g., low network confidence, large temporal jumps, and/or multi-camera inconsistency.

While attractively simple, this “remove-then-interpolate” strategy is suboptimal for several reasons. First, it can be challenging to automatically and reliably choose thresholds to determine which bad frames should be dropped. Second, using simple temporal interpolation to replace removed keypoints ignores useful spatial constraints (such as the multi-view PCA loss) which, as we have seen, can significantly improve the estimation of uncertain keypoints. Third, networks often make errors confidently that may not be corrected with this simple strategy. Finally, even error frames often contain partial information about keypoint location (for example, a keypoint may be near the estimated value, but not match the estimated value exactly), and removing error frames completely discards this useful partial information.

A number of more complex denoising strategies have been proposed in the single-animal pose tracking literature [5–8], in addition to post-processing strategies for the multi-animal tracking case [9, 10] that are beyond the scope of this paper. These advanced techniques vary widely in their complexity, computational demands, assumptions, generality, outlier-handling logic, etc., but they all operate on the output of a single network. As we have seen, the output of a single network (particularly a fully-supervised network) can be highly unreliable, and moreover the reliability (as measured by the ensemble variance) can vary sharply from frame to frame. Without well-calibrated information about the reliability of each frame, it can be difficult to correct network errors.

Previous work [11] showed that training an ensemble of multiple networks and using the mean prediction leads to better pose estimates. Our Ensemble Kalman Smoother (EKS) denoises the ensemble means and leads to further drastic improvements in pixel errors (main text Fig. 5). EKS leverages the per-frame and per-keypoint ensemble variance to determine the degree of smoothing by our spatiotemporal priors. To reiterate, EKS extracts information even from “bad” predictions, all without the need for the user to set any manual thresholds to detect these “bad” points. Once the ensemble has been run our method is simpler, faster, more interpretable, and easier to tune than the more complex strategies discussed in [5–7].

However, it is important to note that these more complex methods are complementary to ours: future work could combine our ensembling strategy with the more realistic nonlinear constraints and non-Gaussian observation models developed in these previous papers, to potentially obtain further accuracy improvements (at the cost of longer computational post-processing times).

Finally, a note on terminology: the Ensemble Kalman Smoother we use here is different from the Ensemble Kalman filter commonly used e.g. in weather prediction [12]. The two approaches differ in whether ensembling is performed in the dynamics step or the observation step of the Kalman filter model. In our case, the ensemble is used to generate the observation model (which is then smoothed with a simple linear-Gaussian dynamical system model), whereas in the weather prediction context ensembling is performed over multiple instances of nonlinear dynamics models, which are then combined with a simple Kalman-like observation update.

### 1.3 Differences between DeepLabCut and the Baseline model

#### Backbones

Both packages support multiple backbones. In the results we use AnimalPose10K for all datasets but mirror-fish (where we use ImageNet), whereas DeepLabCut defaults to ImageNet.

#### Heatmap peak finding

As described in the Methods section, Lightning Pose operates on normalized heatmaps, that are valid 2D probability distributions for each keypoint. We find the heatmaps’ peaks (i.e., the predicted width-height coordinates) via a soft argmax. Crucially, this process is differentiable, which is necessary when defining downstream unsupervised loss functions on the width-height coordinates. DeepLabCut, on the other hand, uses a location refinement strategy that requires a hard argmax which is not differentiable, and therefore incompatible with unsupervised losses.

#### Supervised heatmap losses

For the supervised loss, we use the mean-squared error between the target heatmap and the predicted heatmap. DeepLabCut uses a cross-entropy loss for each pixel.

#### Training

Though both use the Adam optimizer, DeepLabCut trains for a certain number of “iterations” (typically 50K) independent of the dataset size, while we use the more common “epochs” method which counts full passes over the labeled dataset, typically 300 at a minimum. As a result there are differences in the learning rate schedule as well.

### 1.4 IBL-paw results

Here, we report results on the “IBL-paw” dataset. As in “IBL-pupil”, we compare the following three approaches: DeepLabCut with custom post-processing (DLC), Lightning Pose’s semi-supervised TCN model followed by the same post-processing (LP; using temporal difference and Pose PCA losses), and a multiview EKS variant which uses an ensemble of *m* = 5 LP models (LP+EKS). Here too, we obtain performance gains (Extended Data Fig. 7 and Supplementary Fig. 4).

We use two cameras to track the paws and, therefore, we can use the multi-view PCA loss to help quantify the rate of paw tracking errors. (Since the dataset does not contain synchronous labeled frames from both cameras, we did not train with multi-view PCA and only use it post-hoc; see the Methods section.)

Specifically, we use canonical correlations analysis (CCA) to find directions of motion that must match in the left and right cameras (see Methods), and then we quantify the correlation values of these two directions of motion. We find that the correlation values obtained by DeepLabCut can be fairly low in some sessions (*r*=0.83*±*0.02, mean*±*sem over 44 sessions), largely due to occlusions or to frames in which one paw is confused for the other. LP networks improve these correlations slightly (*r*=0.86*±*0.02), and the EKS that enforces the multi-view PCA loss pushes these correlation values to 1.0, by construction (Extended Data Fig. 7C,D; see Methods for multi-view EKS details). Next we align trials to movement onset and compute the trial consistency metric introduced above, finding improvements with LP+EKS (Extended Data Fig. 7E,F; DLC 0.07*±*0.01; LP 0.07*±*0.01; LP+EKS 0.17*±*0.02). As in IBL-pupil, we also quantify the correlation between paw speed and neural activity using a simple decoding analysis, and find that LP+EKS but not LP leads to greater decoding accuracy (Extended Data Fig. 7H; DLC *R*^2^=0.23*±*0.02; LP 0.22*±*0.02; LP+EKS 0.26*±*0.02). All quantities reported here refer to the right paw; see Supplementary Table 2 for left paw values, which are qualitatively similar.

## 2 Supplementary Tables

**Supplementary Table 1:** Dataset details. In-distribution (InD) frames are selected from one set of animals/videos; out-of-distribution (OOD) frames are selected from a non-overlapping subset of animals/videos. The number of videos is generally larger than the number of animals, indicating multiple experimental sessions from some animals. The total number of unlabeled frames is also included under “Videos/frames” (rounded to the nearest thousand). See text for IBL details. Frame size is (height *×* width).

|  | FPS | Size | In-distribution (Train) |  |  | Out-of-distribution (Test) |  |  |
| --- | --- | --- | --- | --- | --- | --- | --- | --- |
|  |  |  | Labeled frames | Animals | Videos/frames | Labeled frames | Animals | Videos/frames |
| Mirror-mouse | 250 | 406×396 | 789 | 10 | 17/510k | 253 | 3 | 5/150k |
| Mirror-fish | 300 | 384×512 | 373 | 10 | 28/55k | 94 | 3 | 10/22k |
| CRIM13 | 30 | 480×640 | 3986 | 4 | 37/314k | 1274 | 4 | 19/155k |
| IBL-paw | 60/150 | 102×128 | 6071 | 35 | 84/187k | 1446 | 10 | 19/90k |
| IBL-pupil | 60/150 | 100×100 | 2619 | 26 | 52/279k | 1012 | 8 | 8/72k |

**Supplementary Table 2:** Model performance on IBL data. We quantify pose estimation performance on the IBL data using a range of metrics. For more details on how the metrics are computed, see main text Fig. 6 (pupil) and Extended Data Fig. 7 (paw). An asterisk (^*∗*^) indicates values that are fixed to 1.0 by model construction. Values are mean and standard error of the mean computed over n=65 sessions for the pupil, and n=44 sessions for the paws.

| | | Vert vs Horz diameter $r$ | Trial consistency | Decoding $R^2$ |
| --- | --- | --- | --- | --- |
| IBL-pupil | DLC | 0.36±0.03 | 0.35±0.06 | 0.27±0.02 |
|  | LP | 0.88±0.01 | 0.62±0.07 | 0.33±0.02 |
|  | LP+EKS | 1.00±0.00* | 0.74±0.08 | 0.35±0.02 |
| | | CCA projection $r$ | Trial consistency | Decoding $R^2$ |
| IBL-paw (left) | DLC | 0.85±0.02 | 0.10±0.01 | 0.22±0.02 |
|  | LP | 0.88±0.01 | 0.11±0.01 | 0.23±0.02 |
|  | LP+EKS | 1.00±0.00* | 0.14±0.01 | 0.25±0.02 |
| IBL-paw (right) | DLC | 0.83±0.02 | 0.07±0.01 | 0.23±0.02 |
|  | LP | 0.86±0.02 | 0.07±0.01 | 0.22±0.02 |
|  | LP+EKS | 1.00±0.00* | 0.17±0.02 | 0.26±0.02 |

**Supplementary Table 3:**
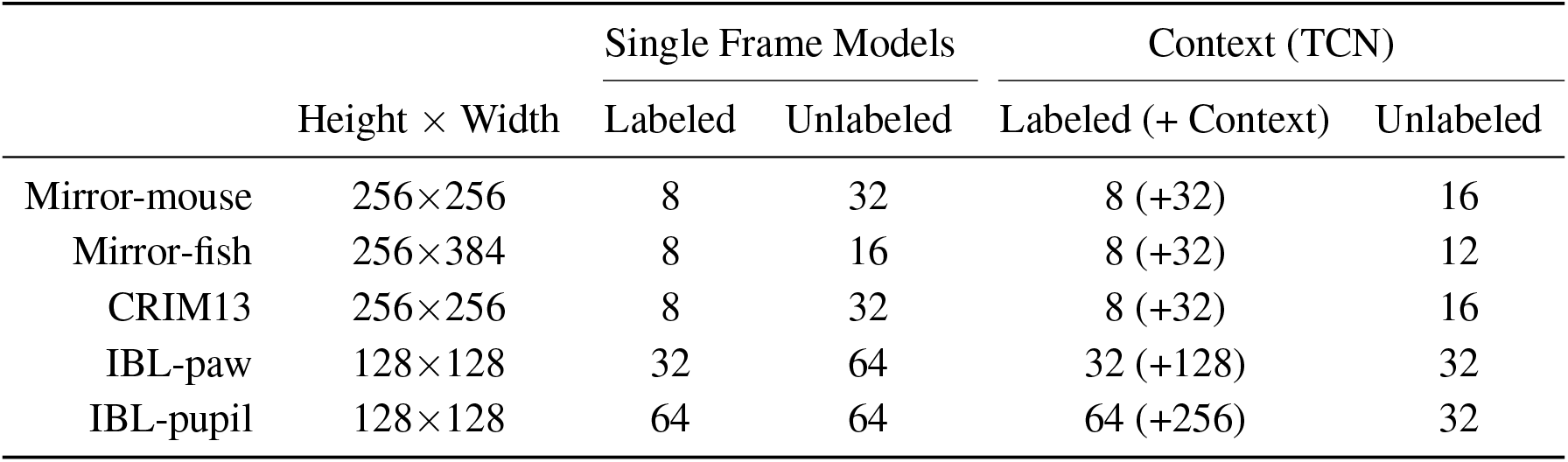
Batch size details. Height *×* Width column shows the dimensions of the *resized* images fed to the network. TCN models with a window of (2*J* + 1) frames (we use *J* = 2) will result in a 5x larger effective labeled batch size (number of unlabeled context frames in parentheses). In general, TCN’s effective labeled batch size is (2*J* + 1)*B* without unlabeled videos and (2*J* + 1)*B* + *T* with unlabeled videos. We intentionally kept labeled batch sizes small so that simple consumer GPUs could train a Single Frame and TCN models with an identical labeled batch size.

| | Height × Width | Single Frame Models | | Context (TCN) | |
| --- | --- | --- | --- | --- | --- |
|  |  | Labeled | Unlabeled | Labeled (+ Context) | Unlabeled |
| Mirror-mouse | 256×256 | 8 | 32 | 8 (+32) | 16 |
| Mirror-fish | 256×384 | 8 | 16 | 8 (+32) | 12 |
| CRIM13 | 256×256 | 8 | 32 | 8 (+32) | 16 |
| IBL-paw | 128×128 | 32 | 64 | 32 (+128) | 32 |
| IBL-pupil | 128×128 | 64 | 64 | 64 (+256) | 32 |

**Supplementary Table 4:** Hyperparameters for unsupervised losses. The table presents the log-weights 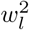 for each of the individual losses *ℒ*_*l*_, where the total loss is parameterized as sum of terms proportional to Gaussian likelihoods, as in [13]: 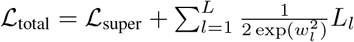

| Train frames | Temporal |  | Multi-view PCA |  | Pose PCA |  |
| --- | --- | --- | --- | --- | --- | --- |
|  | 75 | All | 75 | All | 75 | All |
| Mirror-mouse | 4.75 | 4.75 | 4.5 | 4.5 | 4.5 | 5.0 |
| Mirror-fish | 5.5 | 5.0 | 6.0 | 6.0 | 5.0 | 6.5 |
| CRIM13 | 4.5 | 4.5 | - | - | 4.5 | 5.0 |
| IBL-paw | 6.0 | 6.0 | - | - | - | - |
| IBL-pupil | 4.5 | 4.5 | - | - | 4.0 | 4.0 |

## 3 Supplementary Figures

**Supplementary Figure 1:**
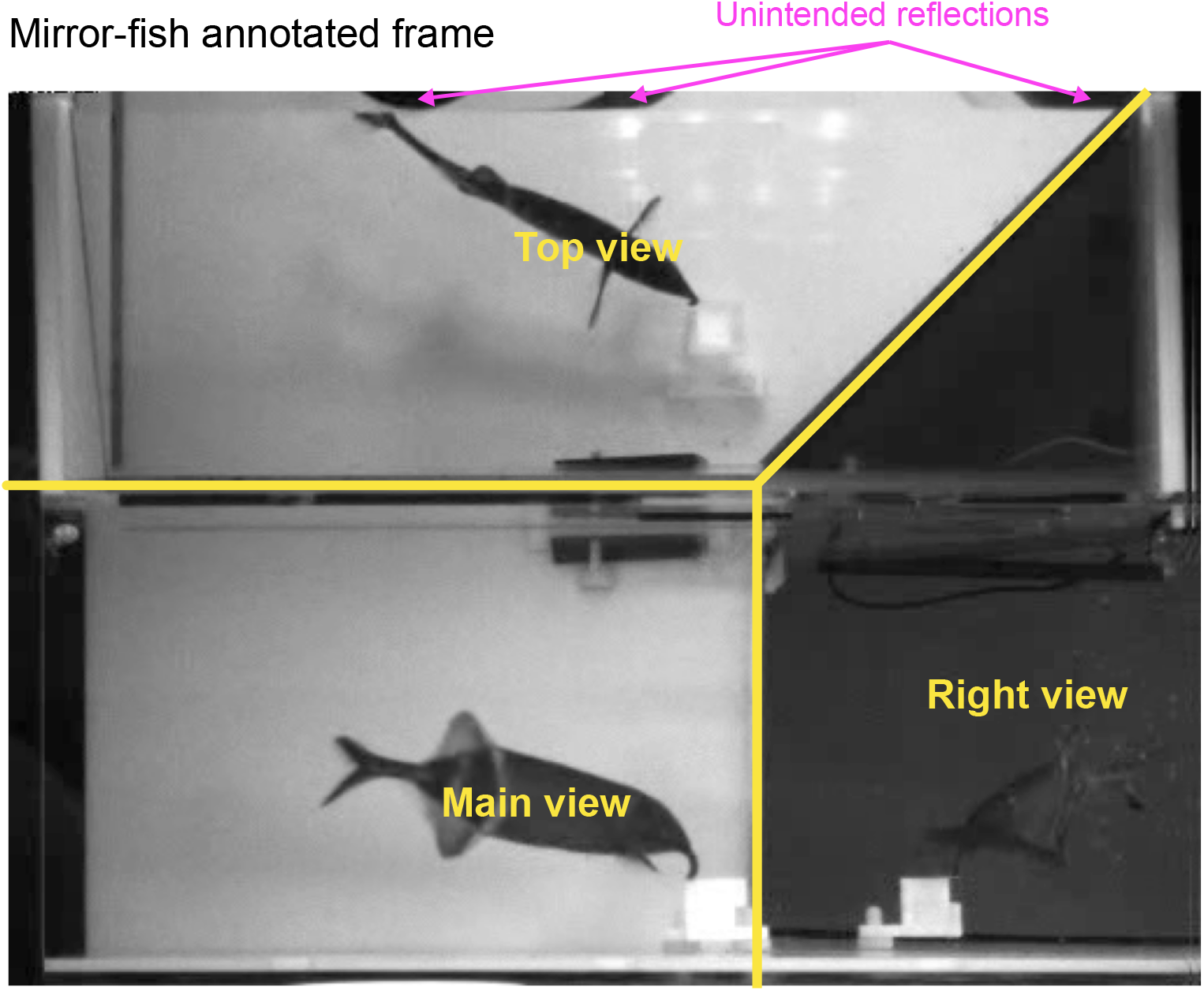
Annotated frame from mirror-fish dataset. The mirror-fish dataset uses a single camera pointed at a tank containing a single fish. The tank contains two mirrors at 45^*°*^, allowing the the camera to capture three roughly orthogonal views. Occasionally, a combination of mirror placement and water levels lead to unintended reflections on the top of the frame.

**Supplementary Figure 2:**
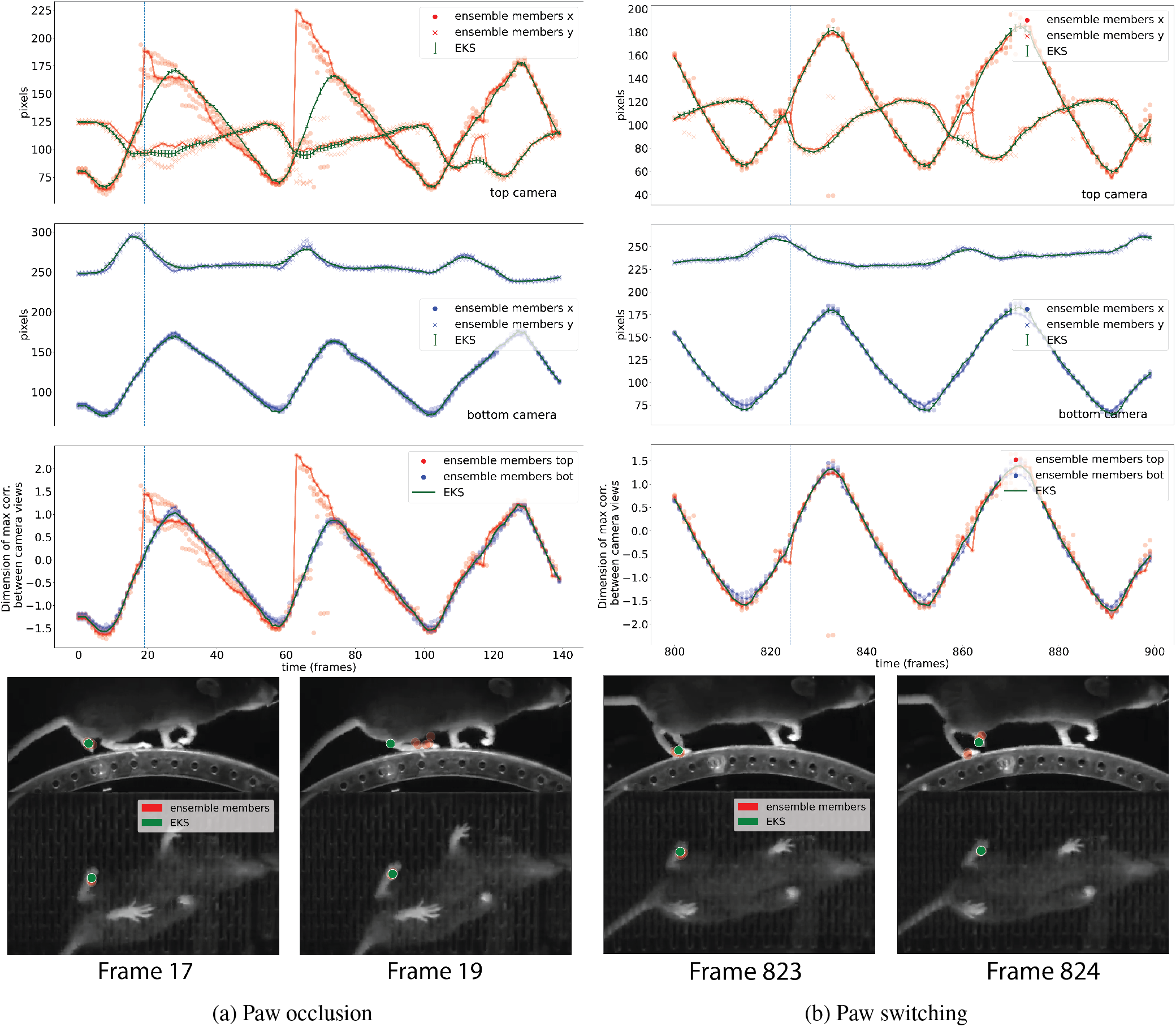
Ensemble Kalman smoothing example with the 75 training frame mirror-mouse dataset. We use ensembles of *m* = 5 semi-supervised TCN models as input to EKS. The two columns show two illustrative examples. Top panels: x and y coordinates of the left hind paw viewed on the top camera. Conventions as in Supplementary Fig. 3. Second from top: x and y coordinates of the left paw viewed on the bottom camera. Third from top: CCA coordinates computed from the top and bottom camera views. Similarly to the IBL-paw dataset (Supplementary Fig. 4), these CCA coordinates should be equal at each frame. The top view is more challenging (the camera is facing the side of the mouse rather than the bottom, and we are tracking the distant paw here, so more occlusions occur), and the ensemble variance is correspondingly larger for the top view; therefore the EKS tracks the bottom view more closely here. Bottom: example frames (indicated with the vertical dashed line above). In the left column, we see an example of paw occlusion - when the left hind paw goes behind the back of the animal all members of the ensemble jump to the nearest visible keypoint. Tracking is accurate in frame 17 and then the occlusion and ensemble confusion is visible in frame 19. Note that the EKS accurately tracks the correct paw here, since it uses information from the more confident camera view to resolve confusion in the more challenging camera view. In the right column, we see an example in which the EKS is able to correct an error due to paw switching in the indicated frames (823-24). For the full video used here, see Supplementary Video 8.

**Supplementary Figure 3:**
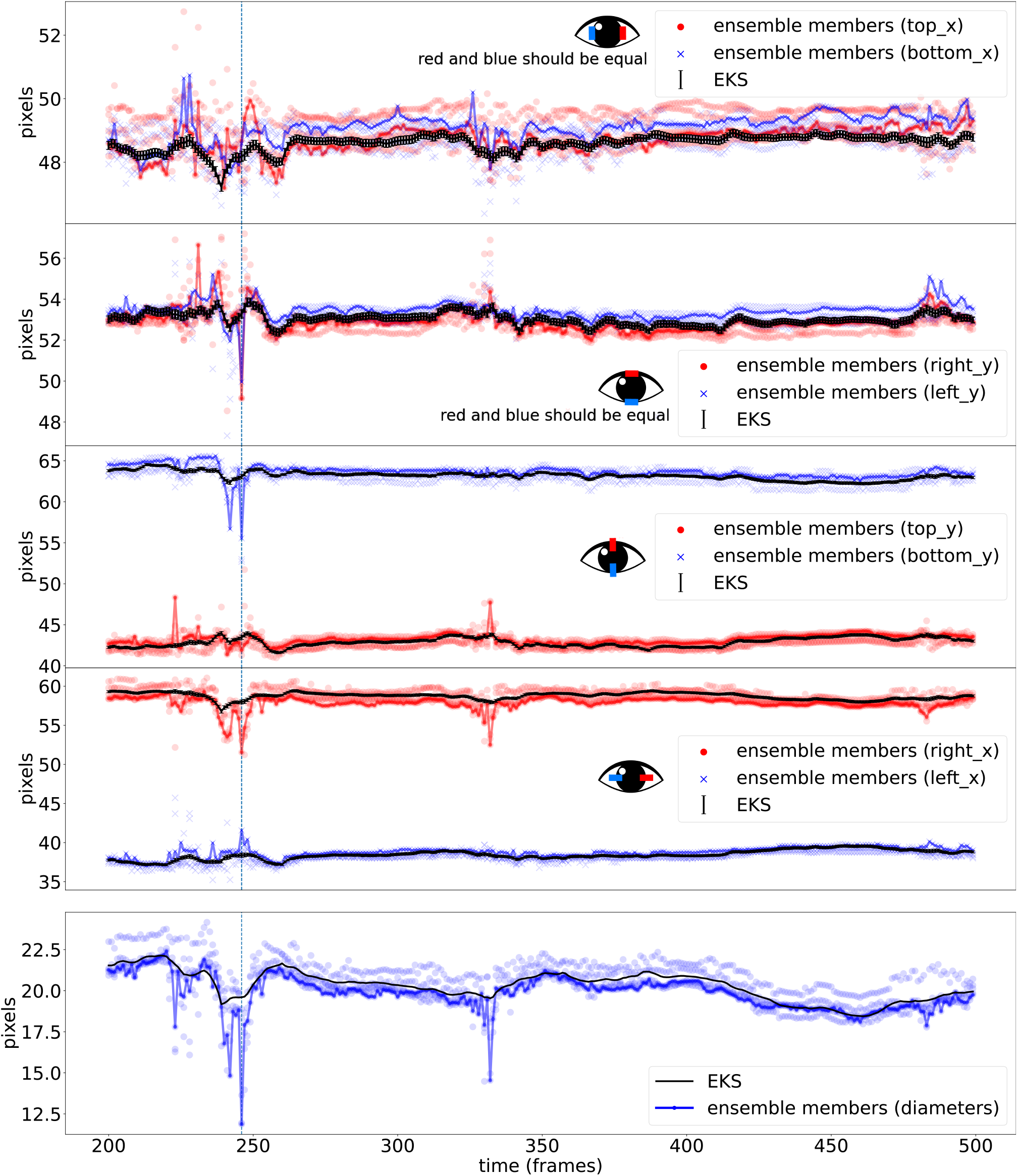
Ensemble Kalman smoothing example with pupil traces. The top four panels show the eight tracked coordinates, while the bottom panel shows the estimated diameter. In each panel, the *×*’s indicate the output of individual ensemble members (semi-supervised TCN models); we have connected the output of a single ensemble member with a solid line across time. The black trace indicates the EKS output. As the pupil keypoints are arranged in a diamond shape, they are paired naturally, e.g. the x coordinate of the top and bottom keypoints should be equal, as should the y coordinate of the left and right keypoint (as indicated by the insets at the bottom right of each of the first four panels). This pairing makes it easy to detect errors: for example, in the first and second panels, any divergence between the red and blue traces are due to tracking errors. Note that tracking errors are common here but are largely resolved by the EKS (an example error is shown using dotted line at frame 246). For the full video used here, see Supplementary Video 9.

**Supplementary Figure 4:**
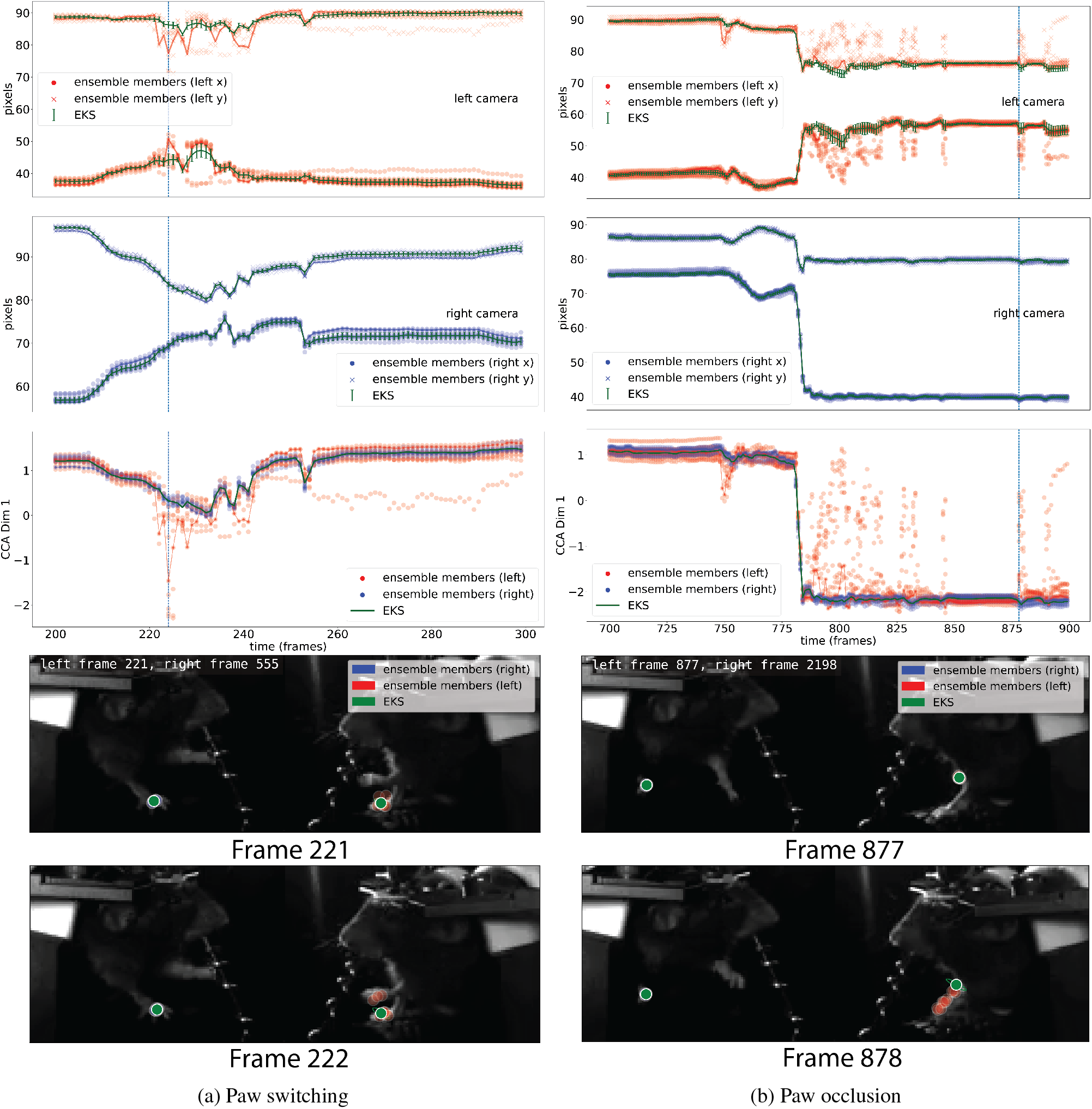
Ensemble Kalman smoothing of baseline supervised models improves pose estimation on IBL-paw data. The two columns show two illustrative examples. Top panels: x and y coordinates of the left paw viewed on the left camera. Conventions as in Supplementary Fig. 3. Second from top: x and y coordinates of the left paw viewed on the right camera. Third from top: canonical correlation analysis (CCA) coordinates computed from the left and right camera views (see Methods). Due to the geometry of multiple cameras recording the same body parts from different directions, these CCA coordinates should be equal at each frame (this is true for the EKS by construction). The left view is more challenging (the paw is further from the camera and the sampling rate of the video is lower), and the ensemble variance is correspondingly larger for the left view; therefore the EKS tracks the right view more closely here. Bottom: example frames (indicated with the vertical dashed line above). In the left column, we see an example of paw confusion - some members of the ensemble (correctly) track one paw, and some (mistakenly) track the other. Tracking is accurate in frame 221 and then confusion between the two paws is visible in frame 222. Note that the EKS accurately tracks the correct paw here, since it uses information from the more confident camera view to resolve confusion in the more challenging camera view. In the right column, we see an example in which the EKS is able to correct an error due to paw occlusion in the indicated frames (877-8). The right ensemble is highly confident so it is tightly packed behind the ensemble-kalman prediction. For the full video used here, see Supplementary Video 10.

**Supplementary Figure 5:**
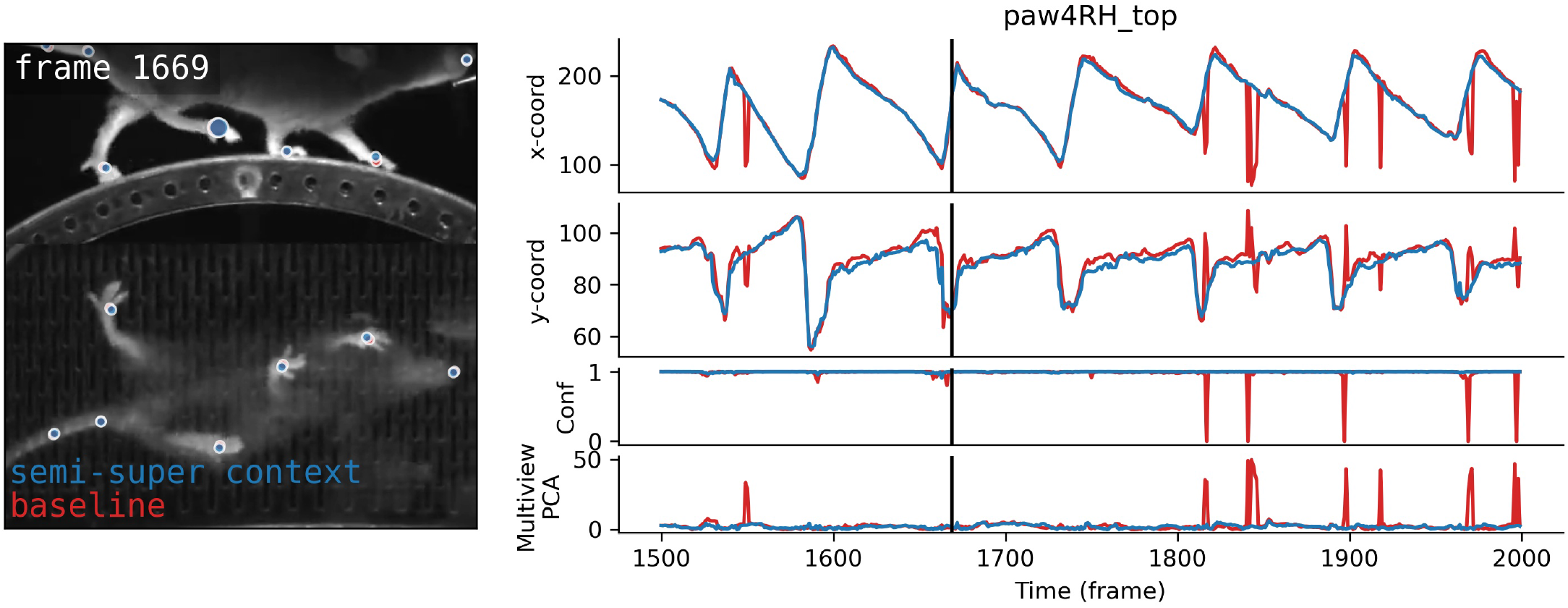
Baseline vs semi-supervised context model predictions. *Left*: Model predictions (with no confidence filtering) for a snippet of OOD video, for both the fully supervised baseline model (red) and the semisupervised context model (blue). The larger marker is the one depicted in the traces. *Right*: x- and y-coordinates of the larger marker, along with confidence values and the multi-view PCA loss. Vertical black bar marks the current frame in the video.

**Supplementary Figure 6:**
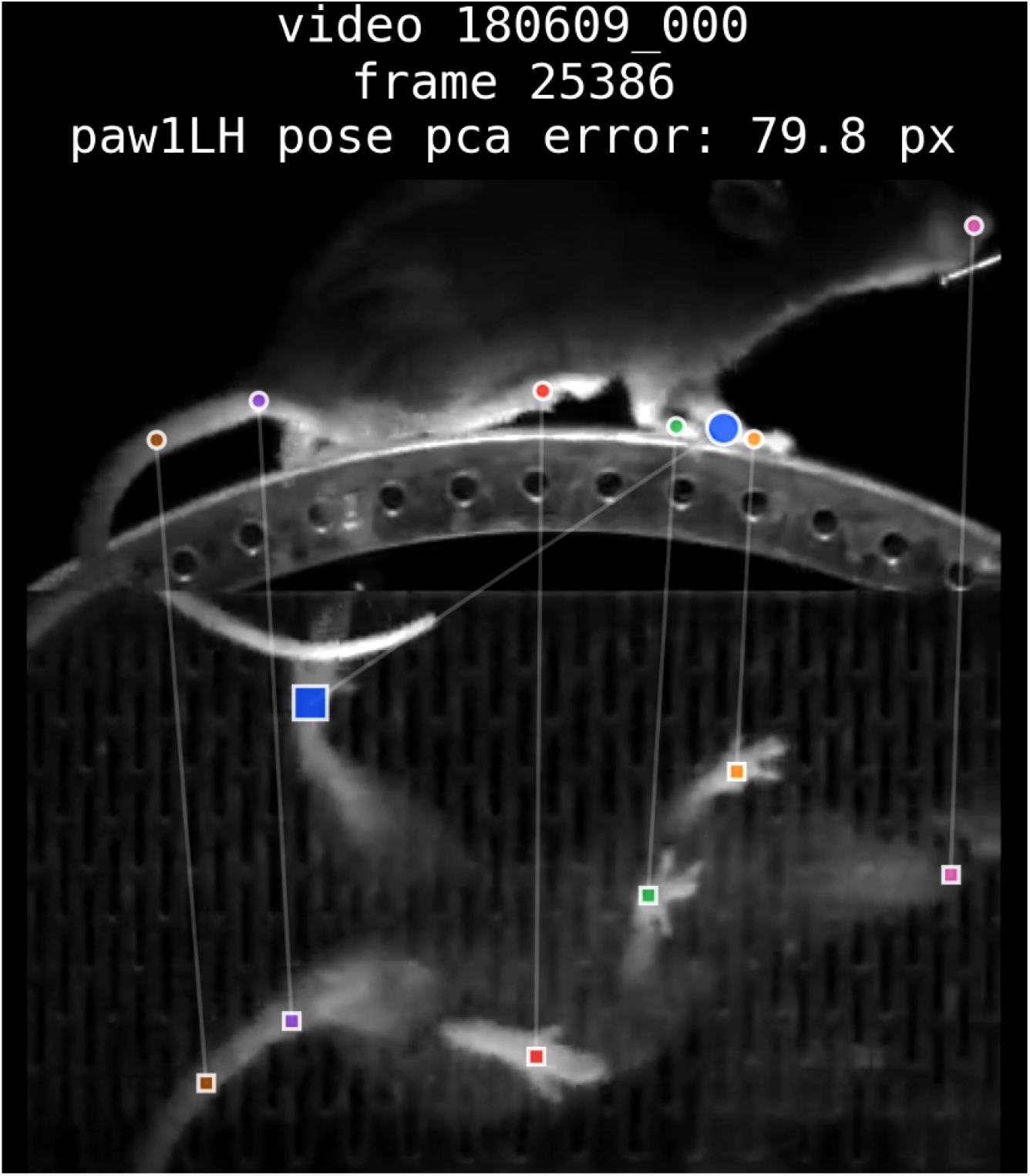
Selected frames with high Pose PCA errors. The video is composed of 100 unlabeled video frames with high single-body-part Pose PCA errors, averaged across all views in the mirrored datasets (DLC models using all available training frames). The body part in question is indicated in the text of the title, as well as the enlarged markers in the frame. In the mirrored datasets the keypoint is enlarged in all views. Each body part is given a unique marker color, which is shared across views (mirror-mouse, mirror-fish) or animals (CRIM13); marker shape is unique for each view/animal. Skeleton lines highlight prediction errors. The videos demonstrate that high values of the Pose PCA error generally capture erroneous predictions; however, high values are occasionally associated with correct predictions on rare poses that are not included in the PCA subspace. To avoid redundant frames, we choose the 1000 frames with highest Pose PCA errors, then fit k-means on these keypoints using 100 clusters, and select one frame from each cluster to display in the video.

**Supplementary Figure 7:**
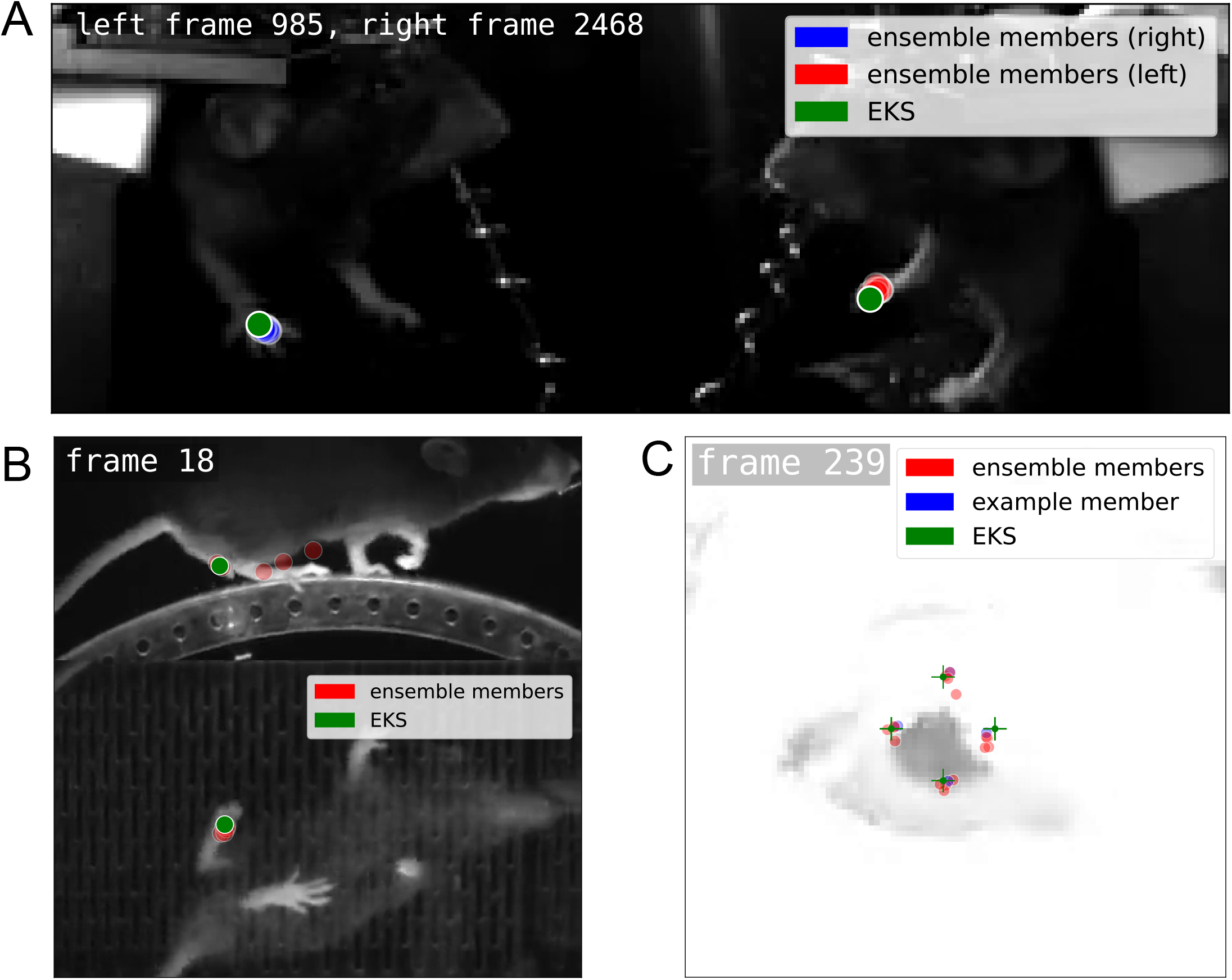
Ensemble kalman model predictions. **A**: Video frames for IBL-paw dataset from two cameras (right and left) overlaid with predicted markers from ten supervised baseline models (blue and red) and the Ensemble Kalman Smoother (green). The cameras for this dataset have different sampling frequencies so the closest frames are visualized together for this video. All blue points are closely packed together behind the green point. **B**: Video frames for 75 frame mirror-mouse dataset with two camera views (top and bottom) overlaid with predicted markers from five semi-supervsied baseline models (red) and the Ensemble Kalman Smoother (green). **C**: Video frame for IBL-pupil dataset with predicted markers from ten supervised baseline models (red), one example supervised baseline model (blue), and the Ensemble Kalman Smoother (green).

**Supplementary Figure 8:**
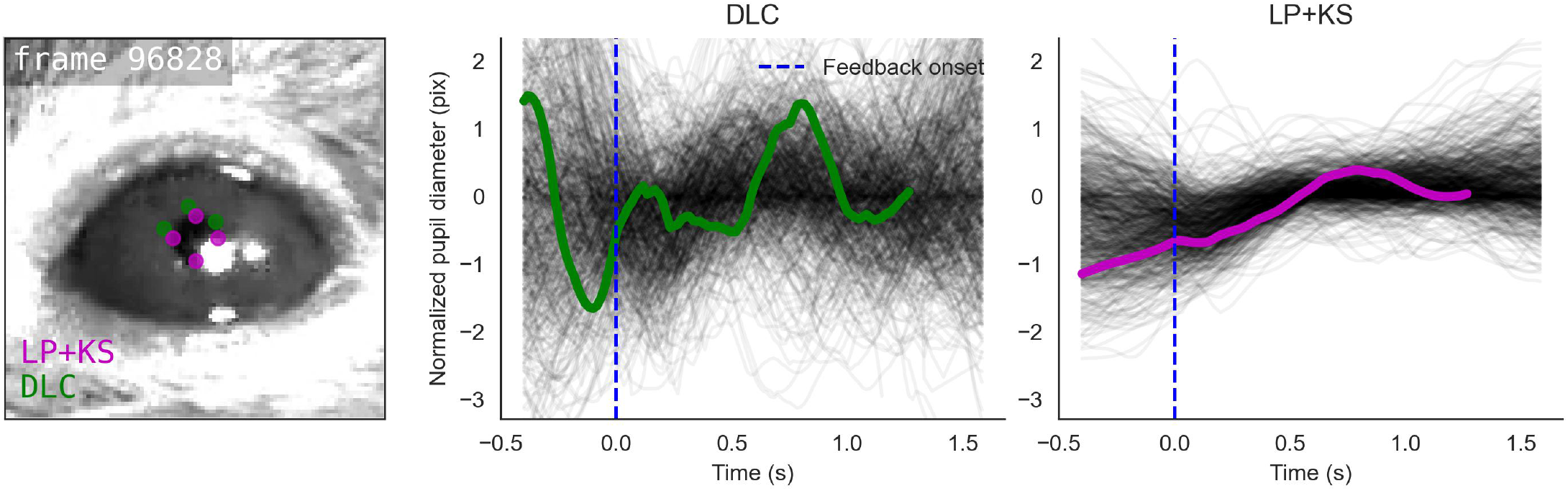
Trial-by-trial markers and traces for IBL datasets. Left: Video frame overlaid with predicted markers from the DLC model (green) and the ensemble Kalman smoother (LP+EKS; magenta). Markers with confidence <0.9 are not displayed. **Center**: Black traces show smoothed DLC pupil diameter from a subset of correct trials, aligned to feedback onset (reward delivery). The green trace highlights the pupil diameter of the current trial displayed on the left. **Right**: Pupil diameters from the LP+EKS model. For the paw video, instead of displaying per-trial traces in black we display the mean and 95% confidence interval.

**Supplementary Figure 9:**
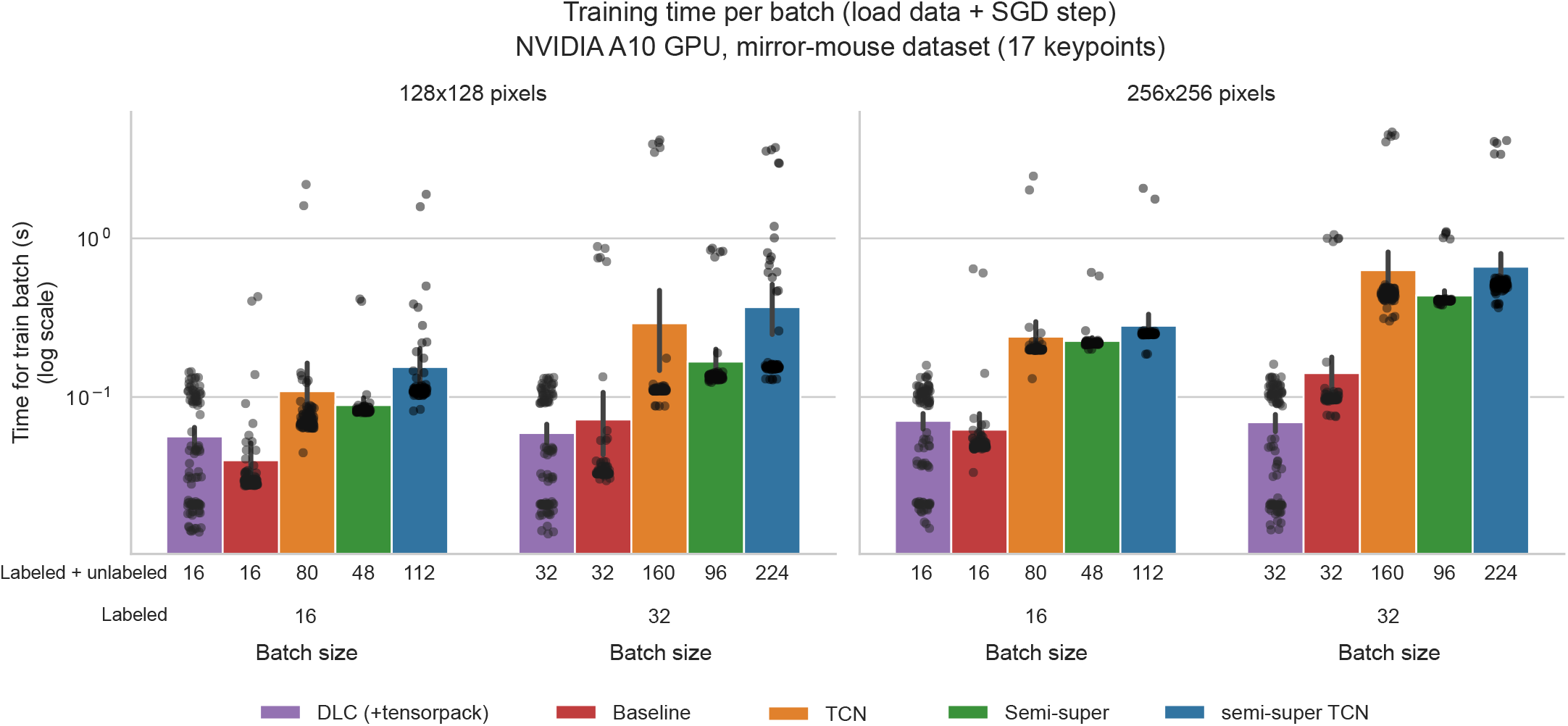
Training time per batch. Each bar depicts the mean batch processing time (in seconds) and 95% CI over n=100 batches, with each of the batches overlaid as a point, using a log-scale spacing for the y-axis. Left panel: 128 *×* 128 images; right panel: 256 *×* 256 images. Each panel is divided into a labeled batch size of 16 (left) and 32 (right). The upper x-axis label denotes the total batch size, comprised of both labeled and unlabeled frames.

**Supplementary Figure 10:**
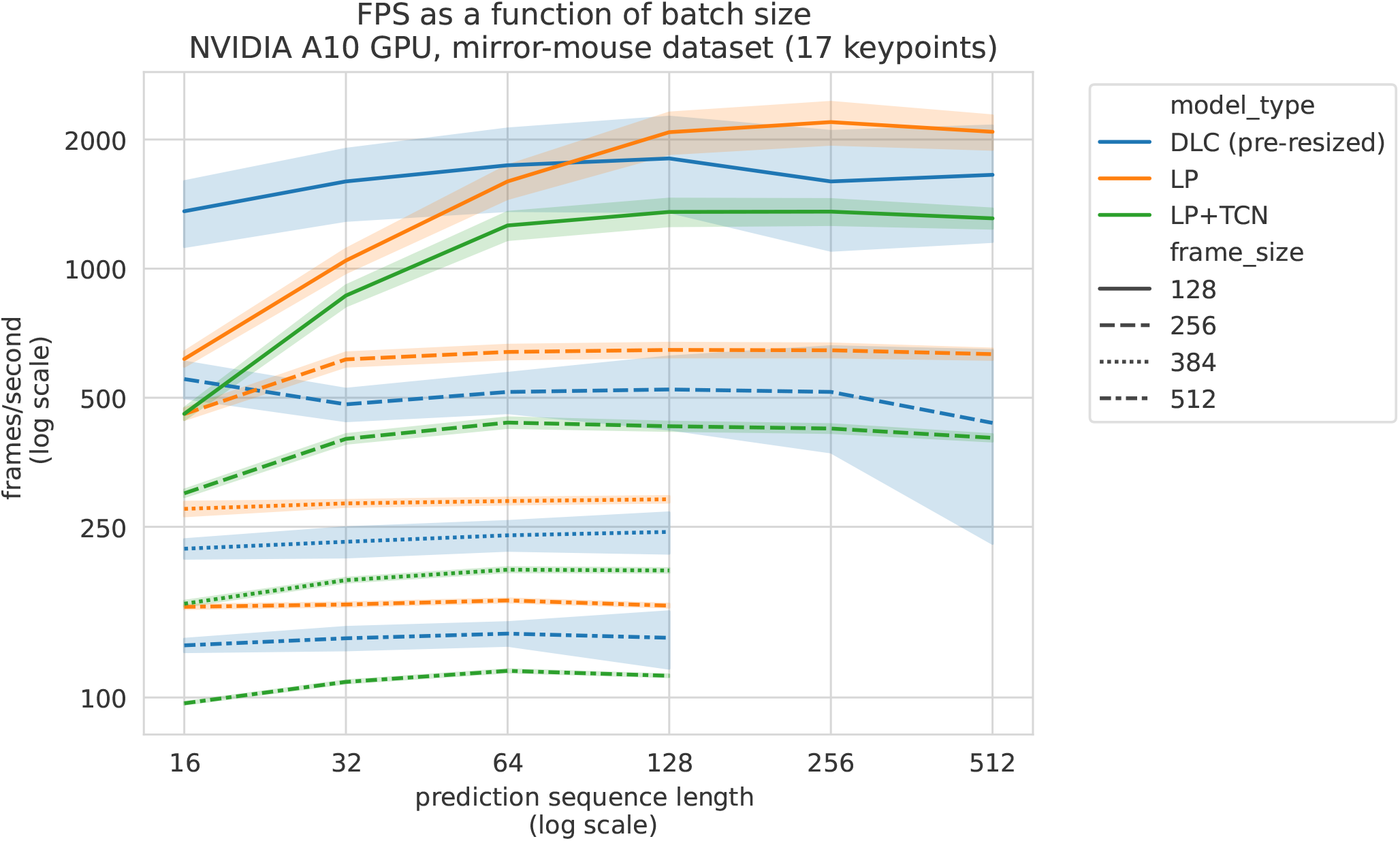
Prediction throughput. x-axis: prediction sequence length (frames), log-spaced. y-axis: frames per second, log-spaced. Colors indicate model types, line styles indicate the frame sizes. For image sizes 384 and 512, sequence lengths did not exceed 128 due to GPU memory constraints. Plotted are means *±* standard errors across n=5 videos.

### 4 The International Brain Laboratory consortium members

Larry Abbot, Luigi Acerbi, Valeria Aguillon-Rodriguez, Mandana Ahmadi, Jaweria Amjad, Dora Angelaki, Jaime Arlandis, Zoe C Ashwood, Kush Banga, Hailey Barrell, Hannah M Bayer, Brandon Benson, Julius Benson, Jai Bhagat, Dan Birman, Niccolò Bonacchi, Kcenia Bougrova, Julien Boussard, Sebastian A Bruijns, Robert Campbell, Matteo Carandini, Joana A Catarino, Fanny Cazettes, Gaelle A Chapuis, Anne K Churchland, Yang Dan, Felicia Davatolhagh, Peter Dayan, Sophie Denève, Eric EJ DeWitt, Ling Liang Dong, Tatiana Engel, Michele Fabbri, Mayo Faulkner, Robert Fetcho, Ila Fiete, Charles Findling, Laura Freitas-Silva, Surya Ganguli, Berk Gercek, Naureen Ghani, Ivan Gordeliy, Laura M Haetzel, Kenneth D Harris, Michael Hausser, Naoki Hiratani, Sonja Hofer, Fei Hu, Felix Huber, Julia M Huntenburg, Cole Hurwitz, Anup Khanal, Christopher S Krasniak, Sanjukta Krishnagopal, Michael Krumin, Debottam Kundu, Agnès Landemard, Christopher Langdon, Christopher Langfield, Inês Laranjeira, Peter Latham, Petrina Lau, Hyun Dong Lee, Ari Liu, Zachary F Mainen, Amalia Makri-Cottington, Hernando Martinez-Vergara, Brenna McMannon, Isaiah McRoberts, Guido T Meijer, Maxwell Melin, Leenoy Meshulam, Kim Miller, Nathaniel J Miska, Catalin Mitelut, Zeinab Mohammadi, Thomas Mrsic-Flogel, Masayoshi Murakami, Jean-Paul Noel, Kai Nylund, Farideh Oloomi, Alejandro Pan-Vazquez, Liam Paninski, Alberto Pezzotta, Samuel Piccard, Jonathan W Pillow, Alexandre Pouget, Florian Rau, Cyrille Rossant, Noam Roth, Nicholas A Roy, Kamron Saniee, Rylan Schaeffer, Michael M Schartner, Yanliang Shi, Carolina Soares, Karolina Z Socha, Cristian Soitu, Nicholas A Steinmetz, Karel Svoboda, Marsa Taheri, Charline Tessereau, Anne E Urai, Erdem Varol, Miles J Wells, Steven J West, Matthew R Whiteway, Charles Windolf, Olivier Winter, Ilana Witten, Lauren E Wool, Zekai Xu, Han Yu, Anthony M Zador, Yizi Zhang

